# EpiFoundation: A Foundation Model for Single-Cell ATAC-seq via Peak-to-Gene Alignment

**DOI:** 10.1101/2025.02.05.636688

**Authors:** Juncheng Wu, Changxin Wan, Zhicheng Ji, Yuyin Zhou, Wenpin Hou

**Affiliations:** Department of Computer Science and Engineering, UC Santa Cruz; Department of Biostatistics and Bioinformatics, Duke University; Department of Biostatistics, Mailman School of Public Health, Columbia University

## Abstract

Foundation models exhibit strong capabilities for downstream tasks by learning generalized representations through self-supervised pre-training on large datasets. While several foundation models have been developed for single-cell RNA-seq (scRNA-seq) data, there is still a lack of models specifically tailored for single-cell ATAC-seq (scATAC-seq), which measures epigenetic information in individual cells. The principal challenge in developing such a model lies in the vast number of scATAC peaks and the significant sparsity of the data, which complicates the formulation of peak-to-peak correlations. To address this challenge, we introduce **EpiFoundation**, a foundation model for learning cell representations from the high-dimensional and sparse space of peaks. EpiFoundation relies on an innovative cross-modality pre-training procedure with two key technical innovations. First, EpiFoundation exclusively processes the non-zero peak set, thereby enhancing the density of cell-specific information within the input data. Second, EpiFoundation utilizes dense gene expression information to supervise the pretraining process, aligning peak-to-gene correlations. EpiFoundation can handle various types of downstream tasks, including cell-type annotation, batch correction, and gene expression prediction. To train and validate EpiFoundation, we curated **MiniAtlas**, a dataset of 100,000+ single cells with paired scRNA-seq and scATAC-seq data, along with diverse test sets spanning various tissues and cell types for robust evaluation. EpiFoundation demonstrates state-of-the-art performance across multiple tissues and diverse downstream tasks.

## 1. Introduction

Single-cell ATAC-seq (Assay for Transposase-Accessible Chromatin using sequencing) (Buenrostro et al., 2015) provides unprecedented resolution in understanding the regulatory landscape of individual cells by profiling chromatin accessibility. This technology enables the identification of active regulatory elements such as promoters, enhancers, and transcription factor binding sites at a single-cell level, offering valuable insights into gene regulation and epigenomic heterogeneity across complex biological systems (Zhang et al., 2021; Zu et al., 2023; Cusanovich et al., 2018). This technology is particularly effective in distinguishing cell types, states, and lineages within heterogeneous tissues, as well as uncovering dynamic changes in chromatin accessibility during processes like development, differentiation, **EpiFoundation: A Foundation Model for Single-Cell ATAC-seq via Peak-to-Gene Alignment** and disease progression (Kim et al., 2024; Buenrostro et al., 2015). By linking regulatory elements to gene expression and integrating multi-omics data, single-cell ATAC-seq has become a critical tool for elucidating the mechanisms underlying cellular identity and function, advancing our understanding of gene regulation in both health and disease.

Recent advances in foundation models have revolutionized single-cell analysis by leveraging large-scale pre-training on extensive datasets. Models such as Geneformer (Theodoris et al., 2023), scGPT (Cui et al., 2024a), scBERT (Yang et al., 2022), and scFoundation (Hao et al., 2024a) utilize the self-supervised learning strategy akin to Masked Language Modeling (MLM) employed in BERT (Kenton & Toutanova, 2019). In particular, these models conceptualize a single cell as *“a sentence of genes”*, wherein certain gene expressions are randomly masked, and the model is trained to predict the masked expressions based on the expressions of the remaining genes, thereby capturing gene-to-gene correlations. These models can subsequently be fine-tuned for a variety of downstream applications, providing greater adaptability and efficacy in comparison to approaches tailored to specific tasks. Nonetheless, contemporary foundation models predominantly target scRNA-seq data and lack optimization for encoding scATAC-seq data. While most existing methods for single-cell ATAC-seq data are task-specific (Lal et al., 2021; Ji et al., 2020; Xiong et al., 2019; Ashuach et al., 2023), foundation models have the potential to significantly enhance these methods and enable the extraction of information from a broader perspective.

However, these scRNA-seq solutions cannot be directly applied to scATAC-seq due to the unique challenges associated with modeling scATAC-seq data. The data typically comprises a vast number of peaks (accessible chromatin regions), often ranging from 10^5^ – 10^6^, and suffers from high sparsity due to the limited DNA molecules available for sequencing, typically only two copies per chromosome in diploid cells (Ji et al., 2020). Given the huge scale of peak numbers, encoding all peaks results in unacceptable computational costs. Furthermore, modeling peak-to-peak correlations from such sparse data presents additional difficulties. These challenges necessitate the development of innovative methodologies to effectively analyze and interpret single-cell ATAC-seq data.

In this paper, we introduce EpiFoundation, a foundational model specifically designed for single-cell ATAC-seq data. The model addresses the aforementioned challenges by incorporating the following technique contributions: (1) We argue that determining “which peaks are expressed” suffices for cell representation modeling and propose to model single cells using their non-zero peaks set. This approach enhances the density of cell-specific information within the input data, thereby improving the model’s efficiency and its capacity to capture meaningful regulatory signals. (2) We utilize paired gene expression signals as the training supervision, facilitating the peak-to-gene alignment and ensuring that cell representations are accurately linked to phenotypes, which are typically defined by transcriptomic data.

Moreover, to provide paired transcriptomic and epigenomic information, we curated the **MiniAtlas**, a high-quality single-cell multi-omics dataset with both scRNA-seq and scATAC-seq measurements per cell. As shown in Figure 1, the MiniAtlas spans 19 tissues and 56 cell types, with uniformly called peaks to ensure comparability across samples, serving as the foundation for training and evaluating EpiFoundation. In addition, we also curate heterogeneous test sets from distinct samples to validate our model, including three datasets from bone marrow mononuclear cells (BMMC), kidney, and peripheral blood mononuclear cells (PBMC) tissues, as well as an ALLTissue test set encompassing all tissues in the MiniAtlas.

**Figure 1.**
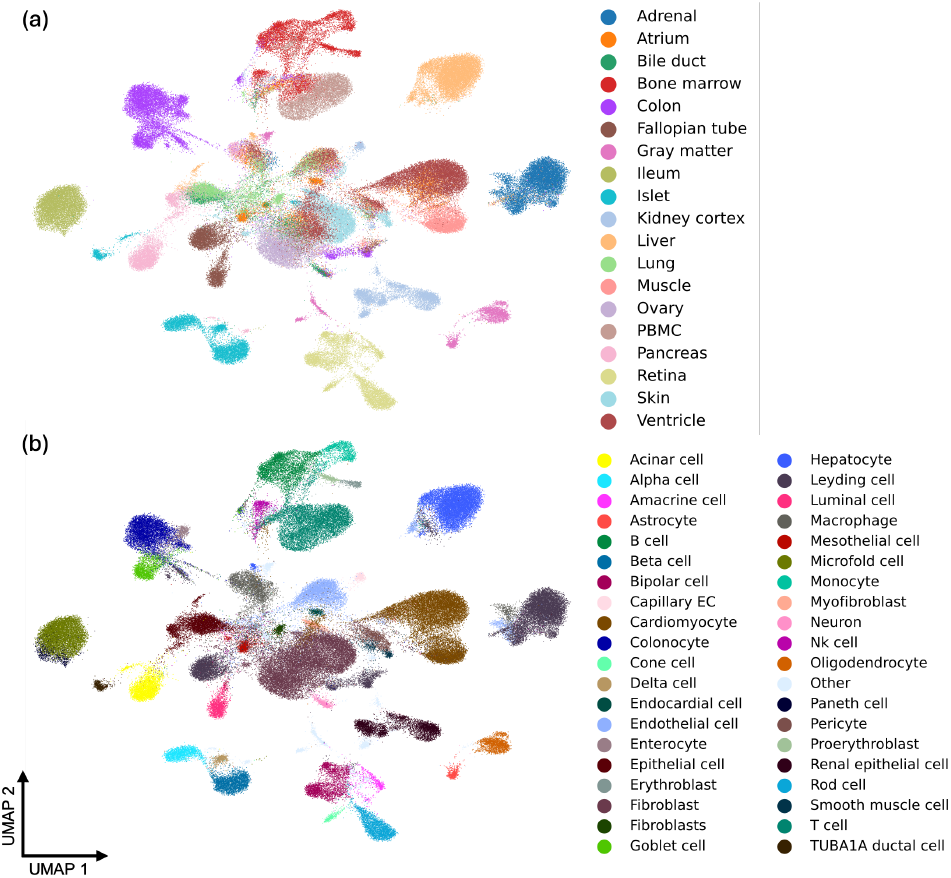
Pre-training data of proposed EpiFoundation. We propose the **MiniAtlas** dataset, containing more than 100,000 scATAC-seq with paired scRNA-seq as training data, across 19 tissues and 56 cell types, facilitating the training of foundation models. We cluster all cells using embedding extracted by EpiFoundation, and color each cell by ground-truth (a) *tissue* and (b) *cell-type* label. EpiFoundation enables modeling cell representation while preserving tissue and cell-type specific information.

EpiFoundation is tailored for crucial scATAC-seq data analysis tasks, encompassing cell type annotation, batch correction, and gene expression prediction. In the domains of cell type annotation and batch correction, the model integrates chromatin accessibility measurements per cell, enabling precise identification of cell identity and exceeding the accuracy attained by state-of-the-art methodologies. Gene expression prediction is inherently supported by the model’s architecture, where gene expression serves as a supervised signal during pre-training. Subsequently, the model is further fine-tuned to predict more fine-grained gene expression. We compare EpiFoundation with Gene Activity (Stuart et al., 2021), a widely applied gene expression prediction methodology. Our model demonstrates state-ofthe-art performance, significantly outperforming existing methods across multiple datasets and metrics.

## 2. Related Works

### 2.1. Foundation models for scRNA-seq data

Geneformer (Theodoris et al., 2023), scGPT (Cui et al., 2024a), scBERT (Yang et al., 2022), and scFoundation (Hao et al., 2024a) are foundation models pre-trained on extensive datasets comprising millions of scRNA-seq profiles. These models exhibit promising performance in a variety of tasks, including cell type annotation, batch integration, perturbation modeling, and gene network inference. Additionally, GenePT (Chen & Zou, 2024) employs GPT-3.5 to generate gene embeddings based on textual descriptions, demonstrating comparable performance. GPT-4 itself can also be viewed as a foundation model and can be applied to downstream tasks such as cell type annotation (Hou & Ji, 2024) and answering genomic questions (Hou & Ji, 2023). LangCell (Zhao et al., 2024) and ZerOmics (Anonymous, 2025) combine the cell encoder with text encoders describing cell metadata, further expanding its applications. Nonetheless, these models lack specific technical design tailored to the challenges in modeling scATAC-seq data.

### 2.2. Foundation models for gene regulation

General Expression Transformer (GET) (Fu et al., 2025) models pseudobulk scATAC-seq signals and incorporates transcription factor information to identify cell-type-specific gene regulation. While effective for regulatory program prediction, this approach sacrifices single-cell resolution, constraining its ability to capture cellular heterogeneity. A recent preprint, CREformer (Yang et al., 2024), integrates bulk epigenetic data with single-cell paired RNA-seq and ATAC-seq for epigenetic regulation tasks, such as predicting master regulators, enhancers, and functional variants. However, both approaches focus on paired data and regulation-related tasks rather than exclusively addressing scATAC-seq data analysis.

### 2.3. Methods for analyzing scATAC-seq data

scCLIP (Xiong et al., 2023) integrates data from two single modalities, SCATE (Ji et al., 2020) and AtacWorks (Lal et al., 2021) to enhance signal quality. SCALE (Xiong et al., 2019) extracts latent features for denoising and cell clustering. BAVARIA (Kopp et al., 2022) uses variational autoencoders for dimension reduction and batch correction. MultiVI (Ashuach et al., 2023), a deep generative model, is designed for multi-omics analysis and single-modality data integration. These task-specific models highlight the need for a foundation model specifically tailored to scATAC-seq data to support a broader range of downstream analyses.

## 3. Method

### 3.1. Problem Formulation

The proposed EpiFoundation aims to address the following problem: consider a matrix 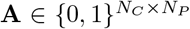 that represents the binarized accessibility signals of peaks. *A*_*i,j*_ = 1 indicates that peak *j* is expressed within cell *i*, and conversely. Herein, *N*_*C*_ and *N*_*P*_ correspond to the total number of cells and the number of peaks in the dataset, respectively. 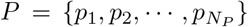 denotes all peaks within the dataset. For each cell *i*, our objective is to construct its cellular representation 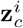 by aligning the peak-to-gene correlations during the pre-training of the model, based on **A**[*i*,:], which represents the accessibility of each peak within *i*. Specifically, the model is trained to predict the paired binary expression of genes within the same single cell (denoted as **B**^*binary*^[*i*,:]), where 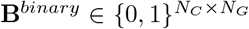 is the binary expression matrix of genes, with *N*_*G*_ indicating the total number of genes. And **B**^*binary*^ is obtained from raw gene expression counts **B**^*raw*^ by:

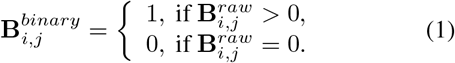

For downstream applications, we extract 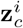 for each cell utilizing the pre-trained weights, and train distinct decoders to predict the cell type label *t*^*i*^ and the fine-grained expression of each gene using 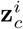. Furthermore, 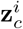 can be viewed as an unbiased representation of cells and employed in the task of batch correction.

### 3.2. Data Collection

In this section, we provide specifics regarding how we collect datasets encompassing the aforementioned data, essential for the training of EpiFoundation. As shown in Figure 1, we collect a 10X Multiome **MiniAtlas** (scATACseq and scRNA-seq coassay) of over 100,000 cells across 19 tissues and 56 cell types. To collect such data, 10X Multiome samples were collected from GEO (Clough & Barrett, 2016) and ENCODE (Snyder et al., 2020) as raw sequencing files in FASTQ format. Raw sequencing data is processed with 10x Cell Ranger ARC software (version 2.0.1) to align the reads to the human GRCh38 genome (10x version 2020-A-2.0.0), which produced a gene-cell count matrix for RNA-seq and a fragment file for ATACseq. All fragment files for ATAC-seq were pooled to call peaks **P** using MACS2 (version 2.2.7.1) (Zhang et al., 2008). The peak cell count matrix **A**^*raw*^ for the ATAC-seq was calculated using the `feature matrix` function provided by the R package Signac (version 1.8.0) (Stuart et al., 2021). The binarized peak-cell count matrix **A** was constructed from **A**^*raw*^ by setting counts to 1 for values greater than 1. The RNA count matrix was normalized and logtransformed using the `NormalizeData` function to obtain **B**^*raw*^. For each sample, cells were clustered based on the information from both RNA and ATAC modalities using `FindMultiModalNeighbors` function provided by Seurat (version 4.3.0) (Hao et al., 2024b). We then computed the Spearman correlation coefficient between cell cluster and cell type expression profiles provided in the DISCO database (Li et al., 2022) to assign a cell type label **t** to each cluster. We provide more details regarding the data collection in the Appendix A.

### 3.3. Model Pre-training

Due to the extensive and sparse characteristics of the peak dataset, embedding all peaks is inefficient. In this paper, we hypothesize that (1) only determining *which peaks are expressed* within the cell *i* suffices to construct its cell-level representation, and (2) the *alignment of peak-to-gene correlations* facilitates cell modeling. To formulate cell representation, as shown in Figure 2(a), we initially transform the set of non-zero peaks alongside their respective chromosomes into **input embedding** (Section 3.3.1). Subsequently, we employ transformer blocks to process the input embedding for generating **cell representation** (Section 3.3.2), and ultimately perform **peak-to-gene alignment** as the pre-training objective (Section 3.3.3).

**Figure 2.**
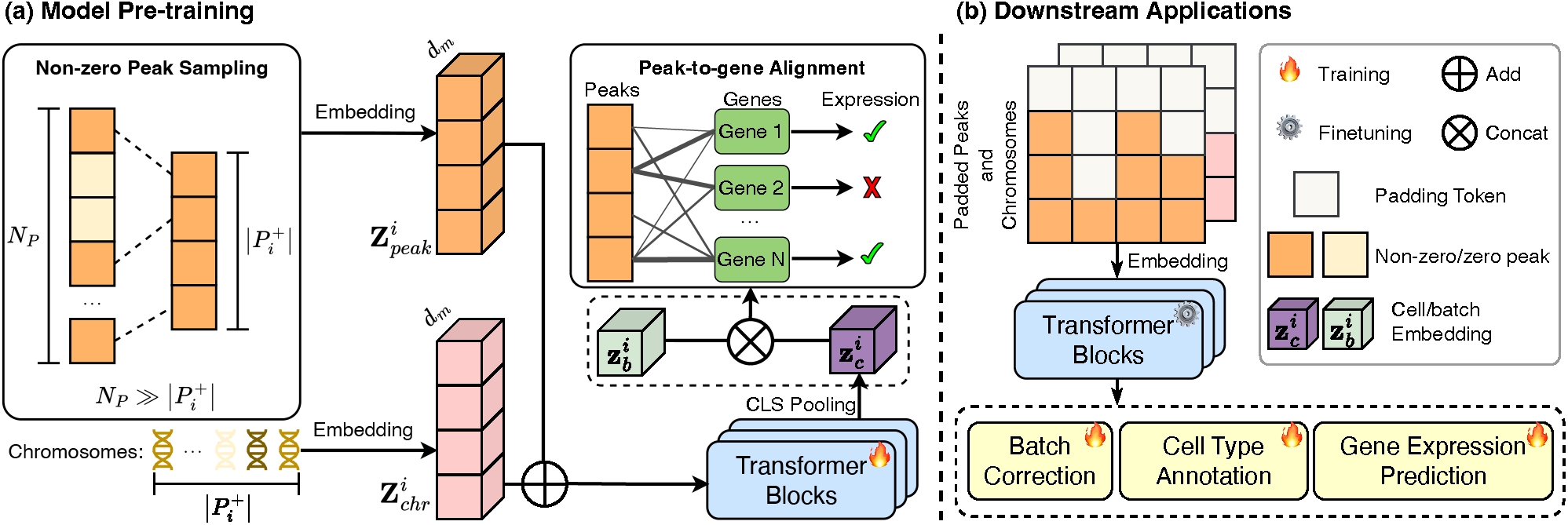
The overview of EpiFoundation. (a) **Model pre-training** with paired ATAC and RNA sequence data. For each single cell, embedding of non-zero peak sequence 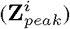 and corresponding chromosomes 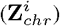 is processed using Transformer blocks to obtain the cell embedding 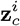. After that, **z**_*c*_ is concatenated with batch embedding 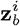 to remove batch effect. We use fused cell embedding to predict binary expression of each genes as training objective. (b) **Downstream application** of EpiFoundation. Pre-trained foundation model can be fine-tuned into downstream tasks including *cell type annotation, batch correction*, and *gene expression prediction*.

#### 3.3.1. Input Embedding

Input embedding of EpiFoundation is composed of two parts: non-zero peaks embedding alongside their corresponding chromosome embedding. Firstly, non-zero peaks embedding for cell *i* can be formulated as:

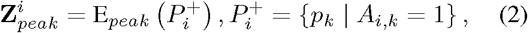

where 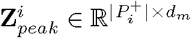 denotes the peak embedding, *d*_*m*_ represents the embedding dim, and 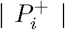 is the number of non-zero peaks within cell *i*. 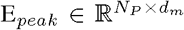 is the embedding layer for peak modeling. For most of the cells, 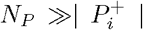.Our proposed non-zero peak embedding improves the density of cell-specific information within the input sequence and facilitates more effective cell modeling. If the number of non-zero peaks 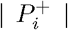 in cell *i* is greater than the pre-defined maximum sequence *L*_*peak*_, we randomly sample *L*_*peak*_ non-zero peaks. In all of our experiments, we set *L*_*peak*_ = 12, 000 to make sure that for more than 95% of cells, all non-zero peaks are contained in the input sequence.

Additionally, we find the corresponding chromosome for each peak in 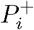, obtaining the chromosome list of cell *i* as:

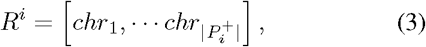

where 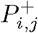 belongs to chromosome *chr*_*j*_ for 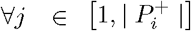. Then we formulate the chromosome embedding as:

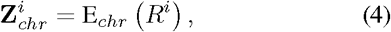

where 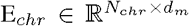 is the chromosome embedding layer. Here, *N*_*chr*_ denotes the total number of chromosomes, including 22 human autosomes and the sex chromosomes X and Y. Finally, the input embedding for cell *i* is formulated as:

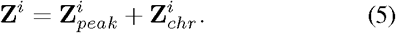

#### 3.3.2. Cell Representation

The input embedding 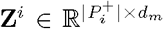 in Equation (5) is then encoded into cell representation by the Transformer blocks. Specifically, we add a [*CLS*] token at the beginning of the input peaks sequence. After *N*_*L*_ layers of Transformer blocks, we obtain the embedding of [*CLS*] token as the representation for cell *i*:

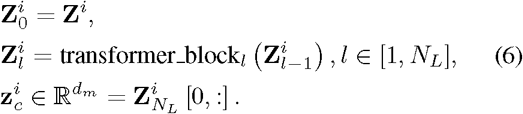

Following scGPT (Cui et al., 2024b), we incorporate separate batch information during the pre-training process to mitigate the bias introduced by different batches of cells. Specifically, for cell *i* which belongs to batch 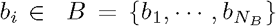 where *N*_*B*_ is the number of different batches in the training data. The batch embedding is generated through an independent embedding layer E_*batch*_. Subsequently, the corrected cell representation 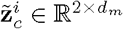 is obtained by concatenating the 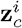 from Equation (6) and 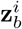. This procedure can be represented by the following formulation:

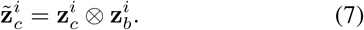

Where 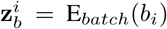 and ⊗ denote the concatenation operation. Note that batch information is only used during the model pre-training to make sure that the cell representation obtained from the non-zero peak set by Equation (6) is unbiased during fine-tuning and evaluation.

#### 3.3.3. Peak-TO-Gene Alignment

During the pre-training stage, EpiFoundation is trained to learn the internal peak-to-gene alignment within the foundation model by predicting binary gene expression. This process is aimed at formulating cellular representations that facilitate the integration of these two modalities. For cell *i*, a gene set 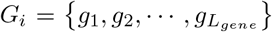 containing *L*_*gene*_ genes is randomly sampled from the gene sets *G*, which encompass a total of *N*_*G*_ genes. In our experiments, *L*_*gene*_ is configured at 8,000, encompassing the majority of non-zero genes across all cells, thereby facilitating dense and effective pre-training supervision. Moreover, each *G*_*i*_ is curated to possess an equal distribution of genes with and without expression, thus guaranteeing that the model is trained without bias. Then, the ground-truth expression of cell *i* on gene set *G*_*i*_ can be denoted as 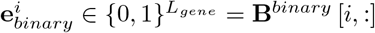 To predict 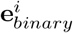, we first get the embedding of genes in *G*_*i*_ by:

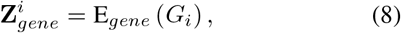

where 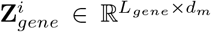 and 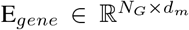 we broadcast 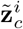 to 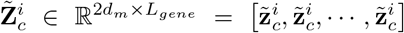 which is concatenated with the gene embedding. This combined representation serves as the input of a simple decoder D_*pre*_ to predict binary expression. In summary, we formulate the binary gene expression prediction process as follows:

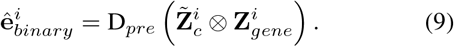

Finally, the loss function for the EpiFoundation model pretraining is formulated as:

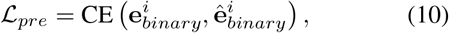

where CE denotes the cross-entropy loss.

### 3.4. Downstream Applications

The pre-trained EpiFoundation model enables the generation of high-quality cell representations by modeling the correlation between ATAC and gene modalities. Consequently, as shown in Figure 2(b), this pre-trained model can be adapted for various downstream applications in single-cell analysis via supervised fine-tuning, including batch correction, cell type annotation, and gene expression prediction.

For the cell type annotation and batch correction tasks, we compile fine-tuning datasets comprising binary peak accessibility counts alongside the corresponding ground-truth cell type labels for various tissues. For cell *i*, its ground-truth cell type label is denoted as *t*^*i*^. We regress the cell representation 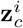 from Equation (6) into the prediction of cell types, and the loss function for cell type annotation fine-tuning is formulated as:

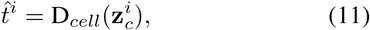

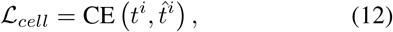

where D_*cell*_ is the cell type decoder, and 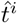 is the cell type prediction.

For the gene expression prediction, our objective is to refine the model to predict fine-grained gene expression values, as opposed to the peak-to-gene alignment the pretraining phase. We normalize and categorize the raw gene expression counts **B**^*raw*^ into *N*_*bin*_ = 10 of expression levels. Categorized gene expression counts are represented as 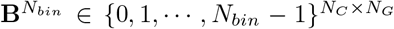. Similarly, we predict categorized gene expression from 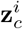, and use mean square error as the fine-tuning loss:

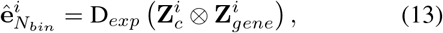

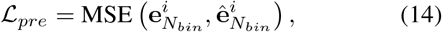

where 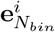 and 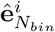 denote ground-truth and predicted _e_ expression values respectively. 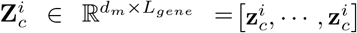 is the broadcasting of 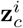, and D_*exp*_ is the categorized expression decoder.

## 4. Experiments

### 4.1. Experiment Settings

#### Training Parameters

The Transformer model within the EpiFoundation comprises 6 attention blocks based on Flashattention2 (Dao, 2023), with embedding dim *d*_*m*_ = 512. We train the model for 140 epochs, employing a batch size of 8 alongside gradient accumulation steps of 20. Additional training specifics are documented in Appendix B.1.

#### Evaluation Datasets

We collect datasets from three tissues for evaluation, including kidney, peripheral blood mononuclear cells (PBMC), and bone marrow mononuclear cells (BMMC) following the same method in Section 3.2. Each dataset is randomly divided into fine-tuning and testing sets. Additionally, we also collect an ALLTissue test set that encompasses all tissues of the training set. More details regarding the data collection can be found in Appendix A. All evaluation data used in this paper will also be made publicly available.

#### Comparing Methods

We select various competing methods for different tasks to validate the effectiveness of the proposed EpiFoundation. For the batch correction task, we compare our methods with state-of-the-art methods including scANVI (Xu et al., 2021), Harmony (Korsunsky et al., 2019), LIGER (Welch et al., 2019), and Principal Component Analysis (PCA) from binary accessibility counts of peaks. For the gene expression prediction task, we compare EpiFoundation with Gene Activity (Stuart et al., 2021). More details of competing methods are provided in Appendix B.2.

#### Evaluation Metrics

For batch correction task, we employ four widely recognized biological conservation metrics alongside two batch integration metrics. Biological conservation metrics are utilized to assess the preservation of meaningful biological variations inherent within a dataset, specifically: (1) Isolated Label Score (ISO) (Luecken et al., 2022), (2) Normalized Mutual Information (NMI), (3) Average Silhouette Width (ASW) (Luecken et al., 2022), and (4) Celltype Local Inverse Simpson Index score (cLISI) (Büttner et al., 2019). Batch correction metrics are designed to evaluate the efficacy of batch effect removal, including Graph Connectivity (GC) and Batch Average Silhouette Width (ASWb) (Luecken et al., 2022). For cell type annotation, we choose accuracy (ACC), F1-score (Macro F1 and Micro F1), and Receiver Operating Characteristic Area Under the Curve (ROC-AUC) as the evaluation metrics. Finally, for gene expression prediction task, we utilize the MSE, Spearman Correlation Coefficient (SRCC), and Pearson Correlation Coefficient (PRCC) between the model prediction and paired ground-truth expression levels as the evaluation metrics. Further details concerning the metrics employed can be found in Appendix B.3.

### 4.2. Cell Type Annotation

Cell type annotation is a crucial task for single-cell analysis, facilitating the comprehension of cellular composition and diversity within a given sample. Proposed EpiFoundation enables the assignment of cell types to individual cells based on the accessibility profile of peaks, demonstrating the potential for single-cell analysis from a novel dimension. As demonstrated in Table 2, we evaluate EpiFoundation on four datasets from different tissues. For each dataset, EpiFoundation is fine-tuned to predict the ground-truth cell-type label for each cell, as indicated in Equation (12). In all datasets, EpiFoundation yielded favorable results across various metrics, including accuracy, macro and micro F1 scores, and ROC-AUC, illustrating its effectiveness in predicting cell type from the non-zero peak set.

Additionally, we demonstrate the classification performance of EpiFoundation in Figure 3, where EpiFoundation demonstrates high classification accuracy, as indicated by the diagonal pattern of high-confidence predictions, highlighting the robustness of EpiFoundation in distinguishing complex cell types, including those with similar transcription profiles.

**Figure 3.**
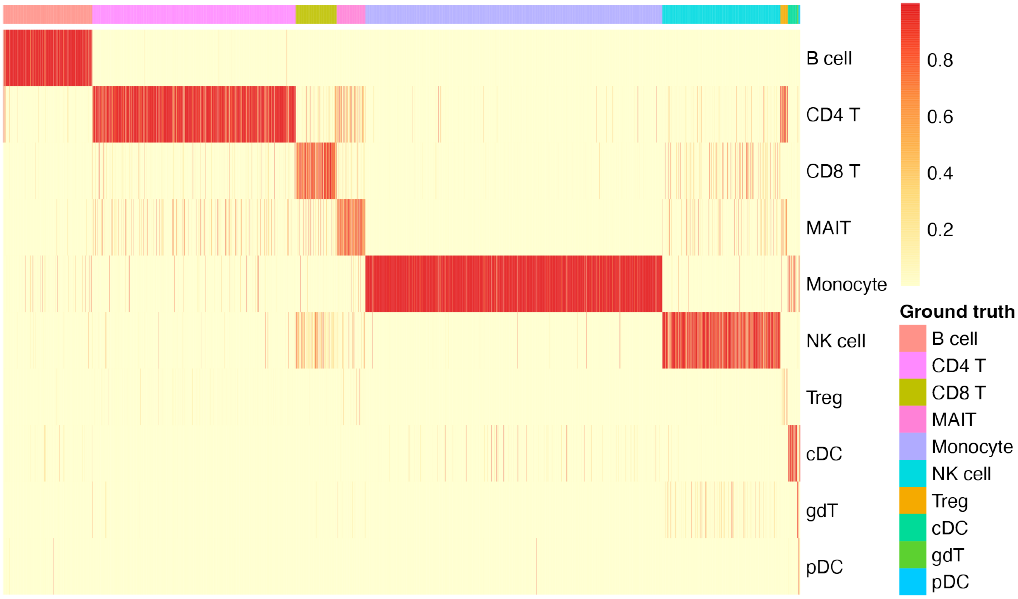
Classification performance of EpiFoundation on PBMC dataset. Each column represents a single cell colored by ground-truth cell type, while each row represents the predicted cell type. The colors in the heatmap refect the softmax score output from EpiFoundation, which indicates the confidence of the model in assigning a cell to a particular cell type.

### 4.3. Batch Correction

Batch effects refer to the variations observed in gene expression or peak accessibility data, which originate from technical discrepancies between distinct batches of samples processed at varying times or in separate laboratory environments, potentially obscuring the true biological differences among single cells. EpiFoundation facilitates the removal of batch effect by modeling robust representation for each individual cell, which conserves essential biological information necessary for aligning peak-to-gene correlations. We extract cell embedding using EpiFoundation fine-tuned on the cell-type annotation task, and compare our method against various state-of-the-art methods. On each dataset, we evaluate the biological conservation and batch integration capabilities of the extracted embedding.

According to the quantitative results presented in Table 1, EpiFoundation exhibits superior performance across the majority of datasets and evaluation metrics, demonstrating its capability to model meaningful and unbiased cell representations. Moreover, as shown in Figure 4, we cluster the cell embedding of different methods using UMAP (McInnes et al., 2018), and color each individual cell by its cell-type and batch labels respectively. The clustering outcomes of EpiFoundation achieve the highest normalized mutual information relative to the ground-truth cell-type labels and exhibit the best graph connectivity.

**Table 1.** Quantitative comparison on batch correction. We compete EpiFoundation with state-of-the-art batch correction methods on datasets from three tissues across four biological conservation metrics and 2 batch integration metrics. EpiFoundation achieves best performance in the majority of the evaluated metrics and datasets.

| Dataset | Method | Biological Conservation |  |  |  | Batch Integration |  |
| --- | --- | --- | --- | --- | --- | --- | --- |
|  |  | ISO↑ | NMI↑ | cASW↑ | cLISI↑ | bASW↑ | GC↑ |
| Kidney | PCA | 0.4568 | 0.3273 | 0.5346 | 0.9936 | 0.8504 | 0.4714 |
|  | scANVI (Xu et al., 2021) | **0.5668** | 0.2007 | 0.4743 | 0.9962 | 0.8890 | 0.7732 |
|  | Harmony (Korsunsky et al., 2019) | 0.4459 | 0.2964 | 0.5375 | 0.9934 | 0.8735 | 0.3995 |
|  | LIGER (Welch et al., 2019) | 0.5288 | 0.0942 | 0.2581 | 0.9700 | 0.7252 | 0.2911 |
|  | **EpiFoundation (ours)** | 0.4995 | **0.5681** | **0.6685** | **1.0000** | **0.9069** | **0.8267** |
| BMMC | PCA | 0.5310 | 0.5039 | 0.4491 | 0.9753 | 0.8240 | 0.4887 |
|  | scANVI (Xu et al., 2021) | 0.4836 | 0.4823 | 0.4742 | 0.9769 | 0.8623 | 0.6500 |
|  | Harmony (Korsunsky et al., 2019) | 0.5241 | 0.4760 | 0.4555 | 0.9598 | 0.8093 | 0.3739 |
|  | LIGER (Welch et al., 2019) | 0.5277 | 0.3942 | 0.4157 | 0.9482 | 0.7164 | 0.4919 |
|  | **EpiFoundation (ours)** | **0.5508** | **0.5773** | **0.5599** | **0.9876** | **0.8959** | **0.6856** |
| PBMC | PCA | **0.7462** | 0.6718 | 0.4461 | 0.9892 | 0.8821 | 0.3234 |
|  | scANVI (Xu et al., 2021) | 0.5798 | 0.4934 | 0.4985 | 0.9831 | 0.8782 | 0.6148 |
|  | Harmony (Korsunsky et al., 2019) | 0.7096 | 0.6355 | 0.4493 | 0.9880 | 0.8571 | 0.2807 |
|  | LIGER (Welch et al., 2019) | 0.5215 | 0.0644 | 0.4378 | 0.7587 | 0.8747 | 0.1868 |
|  | **EpiFoundation (ours)** | 0.6377 | **0.7378** | **0.5965** | **0.9991** | **0.9038** | **0.6837** |

**Table 2.** Performance of EpiFoundation on cell type annotation. We evaluate our model on three tissues, and Mini-atlas which integrate data from all tissues. Among all datasets, EpiFoundation demonstrates promising performance in determining the cell-type based on scATAC-seq.

| Dataset | ACC↑ | Macro F1↑ | Micro F1↑ | ROC-AUC↑ |
| --- | --- | --- | --- | --- |
| Kidney | 0.9135 | 0.7081 | 0.9135 | 0.9866 |
| PBMC | 0.8837 | 0.6299 | 0.8837 | 0.9764 |
| BMMC | 0.7615 | 0.5026 | 0.7615 | 0.9758 |
| ALLTissue | 0.8423 | 0.6934 | 0.8423 | 0.978 |

**Figure 4.**
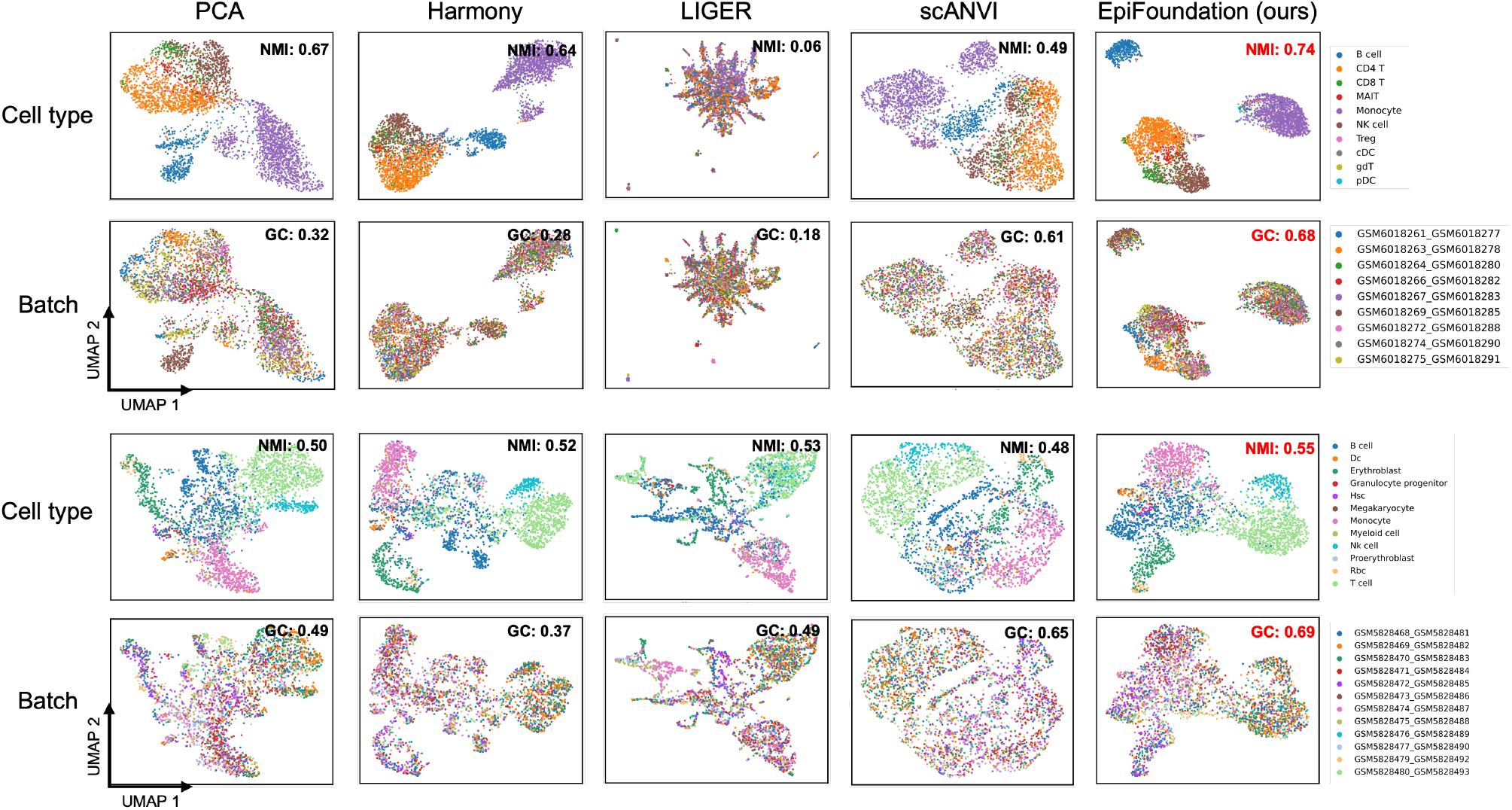
Qualitative comparison on batch correction. We cluster cells in testing set of PBMC and BMMC using embedding from state-of-the-arts methods including Harmony (Korsunsky et al., 2019), LIGER (Welch et al., 2019), scANVI (Xu et al., 2021), PCA of peaks expression, and proposed EpiFoundation. Our method demonstrates best performance across all competing methods, suggesting EpiFoundation can effectively remove batch efforts between different samples, while perserving the meaningful cell-specific variations.

### 4.4. Gene Expression Prediction

EpiFoundation formulates cross-modality correlation between peaks and genes, thus enabling the prediction of how active a specific gene will be within an individual cell from its non-zero peak set. As shown in Equation (14), we finetune the pre-trained EpiFoundation on datasets containing single tissue (PBMC, BMMC, and Kidney) and multiple tissues (ALLTissue), respectively. We compare EpiFoundation with Gene Activity (Stuart et al., 2021), which is widely applied to predict gene expression activity by summarizing the ATAC-seq reads near the transcription start sites of genes. The evaluation focuses specifically on protein-coding genes with results shown in Table 3. When compared to Gene Activity, our method exhibits significantly superior performance across all evaluation metrics and datasets, indicating that EpiFoundation achieves better alignment of peak-to-gene correlations.

**Table 3.** Quantitative comparison on gene expression prediction. We compare proposed EpiFoundation with Gene Activity (Stuart et al., 2021). Our model consistently performs better among all datasets, suggesting the efficacy of EpiFoundation in modeling peak-to-gene correlation.

| Metric | Dataset | Gene Activity | EpiFoundation (ours) |
| --- | --- | --- | --- |
| MSE↓ | PBMC | 10.2098 | **6.7642** |
|  | BMMC | 12.5869 | **8.7789** |
|  | Kidney | 11.5899 | **7.5959** |
|  | ALLTissue | 11.1423 | **9.0777** |
| SRCC↑ | PBMC | 0.1609 | **0.4221** |
|  | BMMC | 0.1766 | **0.3661** |
|  | Kidney | 0.1971 | **0.4030** |
|  | ALLTissue | 0.1772 | **0.3843** |
| PRCC↑ | PBMC | 0.1635 | **0.4776** |
|  | BMMC | 0.1779 | **0.3992** |
|  | Kidney | 0.2021 | **0.4422** |
|  | ALLTissue | 0.1803 | **0.4056** |

### 4.5. Ablation Studies

In this section, we examine the impact of two critical technical strategies employed in the training of EpiFoundation: the incorporation of batch labels to enhance batch correction, and the introduction of chromosome information, respectively. Upon the exclusion of the batch label, the cell embedding 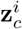 in Equation (6) is employed independently for peak-to-gene alignment, without concatenating it with batch embedding as illustrated in Equation (7). In this setting, the fine-tuning process is identical to baseline EpiFoundation following the Equation (11) and Equation (12).

Similarly, to remove the chromosome information, peak embedding 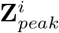 is utilized directly as the model’s input embedding **Z**^*i*^, without the incorporation of chromosome embedding as shown in Equation (5). The chromosome information will be removed during both the pre-training and the fine-tuning stages.

The results of the ablation experiments on the kidney dataset are presented in Table 4. We assess the models that have been pre-trained and fine-tuned using different strategies on the batch correction task, utilizing the NMI and ASWb metrics. A consistent decline in performance is observed upon the exclusion of both types of information, underscoring the efficacy of these strategies in supervising the model to learn high-quality cell representation with better preservation of cell-specific information.

**Table 4.** Ablation experiment on batch and chromosome label. We remove the batch and chromosome label in our pipeline, and evaluating the performance of batch correction on the kidney tissue. A decline performance is observed when removing both of two information, indicating the effectiveness of these settings.

| Batch Label | Chromosomes | NMI↑ | ASWb↑ |
| --- | --- | --- | --- |
| ✓ | ✗ | 0.4695 | 0.8891 |
| ✗ | ✓ | 0.4354 | 0.8986 |
| ✓ | ✓ | **0.5681** | **0.9069** |

## 5. Conclusion

In this paper, we introduce EpiFoundation, a foundational model for scATAC-seq. In order to address the challenge of modeling single cells from the high-dimensional sparse space of peaks, we propose representing cell embeddings using the set of non-zero peaks, alongside peak-to-gene alignment to guide the model to link the correlation between the peak and gene modalities. Furthermore, we have compiled a dataset comprising more than 100,000 scATACseq data with paired scRNA-seq, advancing the progress of research in this domain. Our proposed foundation model achieves state-of-the-art performance across various tasks including cell type annotation, batch correction, and gene expression prediction, presenting significant potential for enhanced single-cell modeling from scATAC-seq. In our future work, we will focus on the development of a more comprehensive single-cell foundation model based on the methodologies and data established in this work, with the objective of unifying multiple modalities including scRNAseq, scATAC-seq, and nucleotide sequences.

## Supporting information

Code for EpiFoundation

## Acknowledgment

We would like to thank the TPU Research Cloud (TRC) program and the Google Cloud Research Credits program for supporting our computing needs. W.H. and Z.J. are supported by the National Institute Of General Medical Sciences of the National Institutes of Health (NIH), under Award Number R35GM150887 and R35GM154865 respectively.

## A. Data Collection and Processing

### A.0.1. Data Download

Raw sequencing data of Multiome was downloaded from both GEO and ENCODE. For data from GEO, meta data and raw data URL can be obtained through R package GEOquery (version 2.62.2). Multiome samples from ENCODE were queried and downloaded directly from the ENCODE data portal (https://www.encodeproject.org/).

### A.0.2. Sequencing DATA PROCESSING

Sequencing reads files in FASTQ format downloaded from GEO and ENCODE were processed with 10x Cell Ranger ARC software (version 2.0.1) to align the reads to the human GRCh38 genome (10x version 2020-A-2.0.0). Cell Ranger generated gene-cell count matrix for RNA and fragments file for ATAC. All ATAC fragments files were merged to call peaks using MACS2 with non-standard and blacklist regions filtered out. Peak-cell count matrix was then calculated using `FeatureMatrix` function provided by Signac. Cells that met the following six criteria were retained: number of RNA reads greater than 1,000; number of RNA reads fewer than 25,000; number of ATAC reads greater than 1,000; number of ATAC reads fewer than 100,000; nucleosome signal (calculated by Signac’s `NucleosomeSignal` function) less than 2; and TSS enrichment score (calculated by Signac’s `TSSEnrichment` function) greater than 1. We also generated a binarized peak-cell count matrix, where counts were set to 1 for values greater than 1.

### A.0.3. Cell TYPE ANNOTATION

Seurat was used to further process the gene-cell count matrix in RNA modality. Specifically, the count matrix was normalized and log-transformed using the function `NormalizeData`. The top 2,000 variable genes were selected by the function `FindVariableGenes`. The normalized gene-cell matrix was scaled by `ScaleData`, and principal component analysis (PCA) was performed by `RunPCA`.

For ATAC modality, the raw peak-cell count matrix was processed by Signac. Specifically, top abundant features were selected using `FindTopFeatures` and kept for later data analysis. The count matrix was normalized using TF-IDF using `FunTFIDF` function, and performed dimension reduction using latent semantic indexing (LSI) provided by `RunSVD`.

RNA and ATAC modalities were integrated using `FindMultiModalNeighbors` with PCA of RNA and LSI of ATAC as the input to construct a weighted nearest neighbor (WNN) graph. Cell clustering was performed using the Louvain algorithm (`FindClusters`) with a resolution of 1. Average RNA expression of each cluster was then computed for cell type annotation.

To assign each cell cluster with a cell type, we downloaded the known cell type expression profile provided through the DISCO database. For each sample, we first selected the corresponding tissue in DISCO and obtained the log-normalized expression profile of each tissue. Cell type was assigned using the algorithm described by DISCO. Specifically, Spearman correlation was computed between each DISCO cell type-specific expression and each Multiome cluster expression using the top 3000 most variable genes. For each Multiome cluster, a cell type was assigned as the cell type in DISCO that has the highest correlation coefficient with the cluster.

## B. Technique Details

### B.1. Training Details

We provide all experiment configurations in the Table 5.

**Table 5.** Additional experiment details. Including (1) Model Configuration, and (2) Training Hyper-parameters

|  | Parameters | Value |
| --- | --- | --- |
| **Model Configuration** | Attention Blocks | 6 |
|  | Attention Heads | 8 |
|  | Max scATAC-seq Length | 12,000 |
|  | Max scRNA-seq Length | 8,000 |
|  | Embedding Dim | 512 |
| **Training Hyper-parameters** | Dropout | 0.15 |
|  | Epoches | 140 |
|  | Learning Rate | 1e-4 |
|  | LR Scheduling | Cosine Annealing |
|  | Gradient Accumulation Steps | 20 |
|  | Batch Size | 8 |
|  | Optimizer | Adam |

### B.2. Comparing Methods

#### B.2.1. Batch Correction

Multiple methods have been developed to correct batch effects for single-cell ATAC-seq. Here, we only include those shown (Luecken et al., 2022) to be top-ranked for single-cell ATAC-seq data integration.

- **PCA** (Principal Component Analysis) is a way to merge samples together without any batch correction. In theory, PCA result will keep the original batch variation of samples. Here we utilized the function `scib.integration.harmony` provided by Python package scIB (version 1.1.7) (Luecken et al., 2022) to obtain PCA embeddings of cells.
- **Harmony** (Korsunsky et al., 2019) is a single-cell batch correction method that utilizes an iterative soft clustering approach to align cells across different batches. It operates by projecting cells into a shared low-dimensional space using Principal Component Analysis (PCA), then iteratively adjusts cell embeddings to minimize batch effects while preserving biological variation. Here we use the function `scib.integration.harmony` provided by Python package scIB (version 1.1.7) (Luecken et al., 2022) with log-normalized binarized peak-cell count matrix as input to compute the latent spaces of Harmony.
- **LIGER** (Linked Inference of Genomic Experimental Relationships) (Welch et al., 2019) uses integrative non-negative matrix factorization (iNMF) to identify shared and dataset-specific factors across batches. It decomposes binarized peak-cell matrices from multiple datasets into shared latent factors that capture biological signals and unique factors that account for dataset-specific variation. The Python package pyliger (version 0.2.3) was adapted to compute LIGER embeddings.
- **scANVI** (single-cell annotation using variational inference) (Xu et al., 2021) is a batch correction and cell type annotation method based on variational autoencoder. It extends the scVI framework by integrating labeled and unlabeled single-cell data to harmonize batches while simultaneously learning cell type-specific latent representations. In this paper, we applied `scib.integration.scanvi` function with binarized raw ATAC-seq count matrix as input to compute scANVI embeddings.

#### B.2.2. RNA Prediction

**Gene activity** is widely applied as the replacement of gene expression in single-cell ATAC-seq data by summarizing the ATAC-seq reads near the transcription start sites of genes. Here, gene activity was calculated using `GeneActivity` function provided by Signac. Raw gene activity was then normalized and log-transformed using the function `NormalizeData` provided by Seurat.

### B.3. Metrics

#### B.3.1. Batch correction

We used two categories of metrics to evaluate the performance of models on batch correction (Luecken et al., 2022). The first category evaluates biological conservation after batch correction and includes the isolated label score (ISO), normalized mutual information (NMI), average silhouette width with respect to cell type (cASW), and cell-type local inverse Simpson index (cLISI). The second category focuses on batch integration and includes the average silhouette width with respect to batch (bASW) and graph connectivity (GC).

- **ISO**: Isolated Label Score (ISO) is a metric to quantify the capability of the integration method to retain meaningful biological structure across batches. For a given cell type *i* that occurs in *k*_*i*_ batches, the ILS is calculated by averaging the ASW values for cell types present in *k*_min_ batches, where *k*_min_ is the smallest number among all *k*_*i*_ values.
- **NMI**: Normalized Mutual Information (NMI) is a metric used to evaluate the similarity between two clusterings by quantifying the amount of information shared between them. It is derived from mutual information, a concept in information theory that measures the dependency between two variables. NMI is normalized to ensure the score ranges between 0 and 1, where 1 indicates perfect alignment between the clusterings, and 0 signifies no shared information. Mathematically, NMI is defined as:

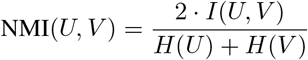

Where:

‐ *U* and *V* represent the two clustering results being compared.
‐ *I*(*U, V*) is the mutual information, calculated as:

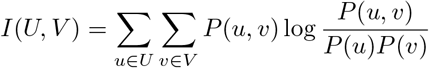

where *P* (*u, v*) is the joint probability of a data point belonging to cluster *u* in *U* and cluster *v* in *V*, and *P* (*u*), *P* (*v*) are the marginal probabilities.
‐ *H*(*U*) and *H*(*V*) are the entropies of *U* and *V*, respectively:

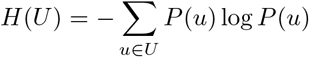 The final NMI is the maximum NMI when compare clustering result under different clustering resolutions to groundtruth cell type labels.

- **ASW**: Average Silhouette Width (ASW) is a metric used to evaluate the quality of clustering by measuring how well each data point lies within its assigned cell types (cASW) or batches (bASW). It is derived from the silhouette score, which assesses the cohesion and separation of clusters. The silhouette score for a single data point *i* is calculated as:

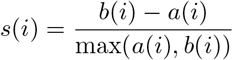

where:

‐ *a*(*i*) is the average distance between *i* and all other points within the same cluster (intra-cluster distance).
‐ *b*(*i*) is the minimum average distance between *i* and all points in the nearest neighboring cluster (inter-cluster distance).

The value of original ASW should be [− 1, 1] where higher value indicate better biological conservation in cASW and pool batch correction in bASW. To make the result consistency, cASW is scaled to the range of 0 to 1 by: 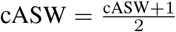. bASW is transformed to the range of 0 to 1 through bASW = 1 − |bASW| so that bigger bASW values indicate better batch correction.

- **cLISI**: Local Inverse Simpson’s Index (LISI) is a metric used to evaluate the performance of integration algorithms. It measures the local diversity of cells in a neighborhood, quantifying how well cells from cell types (cLISI) are mixed. Mathematically, LISI is derived from the Simpson’s Index, which measures diversity within a neighborhood. For a cell *i*, the local inverse Simpson’s index is calculated as:

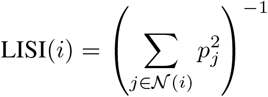

where:

‐ 𝒩 (*i*) is the neighborhood of cell *i* (defined by *k*-nearest neighbors in the embedding space).
‐ *p*_*j*_ is the proportion of cells in the neighborhood belonging to a cell type or batch *j*.

A higher LISI indicates between mixing of cell types or batches. To make value score of each metric consistent, we applied linear transformation to cLISI as LISI = (*L* − LISI)*/*(*L* − 1) where *L* is the number of unique cell types.

- **GC**: Graph connectivity measures how well cells of the same cell type are connected within a KNN graph. Mathematically, for a given cell *i* with group label *g*_*i*_, let 𝒩 (*i*) represent its set of *k*-nearest neighbors in the KNN. The connectivity score for *i* is defined as:

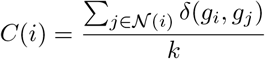

where:

‐ *δ*(*g*_*i*_, *g*_*j*_) = 1 if *g*_*i*_ = *g*_*j*_, and 0 otherwise.
‐ *k* is the number of nearest neighbors considered.

A high KNN connectivity score indicates that cells from the same cell type are tightly connected, reflecting good preservation of local structure and better mixing of batches.

#### B.3.2. Cell TYPE CLASSIFICATION

The performance of cell type classification was evaluated using accuracy(ACC), Macro F1 score, Micro F1 score, and ROC-AUC.

- **ACC**: Accuracy measures the proportion of correctly classified instances among the total instances in a dataset. This metric is calculated using the function `sklearn.metrics.accuracy score`.
- **Macro F1**: The Macro F1 score is calculated as the averaged F1 score for each class. This metric is computed through `sklearn.metrics.f1 score` with the parameter `average=‘macro’`.
- **Micro F1**: Micro F1 score is computed through `sklearn.metrics.f1 score` with the parameter `average=‘micro’`. In order to calculate Micro F1, it computes the overall precision and recall across all classes.

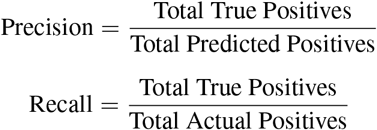

Using the global precision and global recall, the Micro F1 score is:

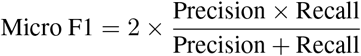

#### B.3.3. Gene PREDICTION

Mean square error (MSE), Spearman Correlation Coefficient (SRCC), and Pearson Correlation Coefficient (PRCC) were computed between predicted gene expression and the ground-truth gene expression from RNA modality. Specifically, each metrics were computed as below:

- **MSE**: The Mean Squared Error (MSE) is a common metric used to measure the average squared difference between predicted values (*ŷ*_*i*_) and actual values (*y*_*i*_). It is defined as:

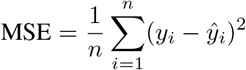

where:

‐ *n* is the number of observations,
‐ *y*_*i*_ is the true value of the *i*-th observation,
‐ *ŷ*_*i*_ is the predicted value of the *i*-th observation.

- **SRCC**: The Spearman correlation coefficient (*ρ* or *r*_*s*_) is a non-parametric measure of the strength and direction of the association between two ranked variables. It is calculated using the formula:

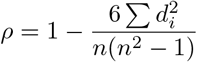

where:

‐ *d*_*i*_ is the difference between the ranks of the corresponding values in the two vectors,
‐ *n* is the length of each vector.

SRCC of each gene was calculated using the function `scipy.stats.spearmanr` and then averaged.

- **PRCC**: The Pearson correlation coefficient (*r*) measures the strength and direction of the linear relationship between two variables, *X* (predicted expression of a gene) and *Y* (ground truth expression of the same gene). It is calculated as:

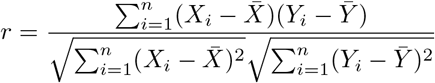

where:

‐ *n* is the number of data points,
‐ *X*_*i*_, *Y*_*i*_ are the individual data points of *X* and *Y*,
‐ 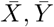 are the means of *X* and *Y*.

PRCC of each gene was calculated using the function `scipy.stats.pearsonr` and then averaged.

## C. Additional Experiment Results

Here we provide additional experimental results. Figure 5 compares the clustering map between EpiFoundation and other batch correction methods on the Kidney dataset. And Figure 6 demonstrates additional classification heat-maps for the cell type annotation task on BMMC and Kidney datasets.

**Figure 5.**
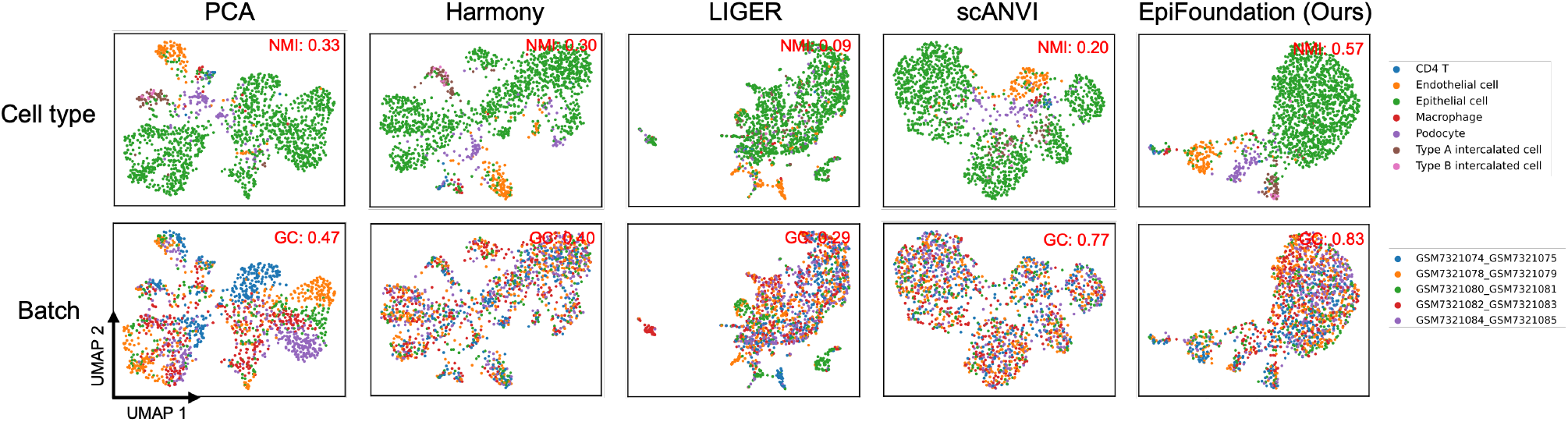
Qualitative comparison on batch correction on Kidney dataset.

**Figure 6.**
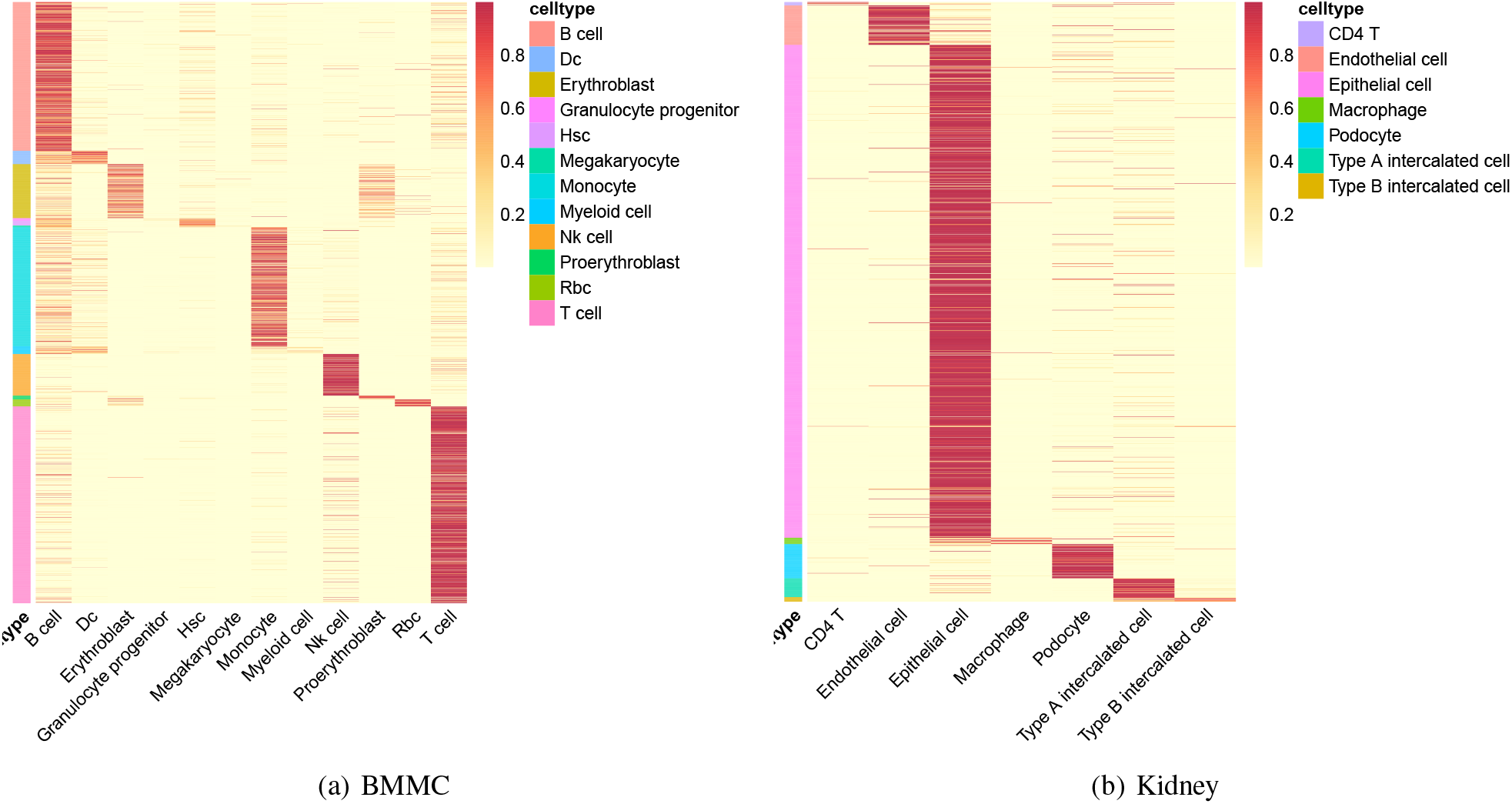
Classification performance of EpiFoundation on BMMC and Kidney dataset.

