## Supplementary material for "EpiFoundation: A Foundation Model for Single-Cell ATAC-seq via Peak-to-Gene Alignment": Code for EpiFoundation: mini_atlas.yml

```
task_name: mini_atlas_eval
train:
  # for ditributed training
  local_rank: 0
  # random seed
  seed: 2002

  # training hyperparameters
  batch_size: 8
  lr: 1e-4
  epochs: 150
  gradient_accumulation_steps: 20
  amp: True
  save_ckpt_freq: 20
  resume: True

  model:
    encoder: transformer
    pretrained: /home/jwu418/workspace/scMultiomics/result/mini_atlas/ckpts/Epoch_100_Step_187900_mini_atlas.pth # set to None if not using pretrained model
    # pretrained: null
    embedding_method: id_only
    atac_max_len: 12000
    rna_max_len: 8000
    embedding_dim: 512
    num_layers: 6
    head_num: 8
    head_dim: 1024
    dropout: 0.15
    additional_config_path: /path/to/additional_config.json
    cell_emb_style: cls
    mvc_arch_style: concat query
    use_batch_labels: False
    use_chr_labels: True
  
  cell_type_epochs: 0
  metric: True

valid:
  freq: 2

data:
  bin_num: &bn 2
  append_cls: True
  train:
    atac_path: /home/jwu418/workspace/data/ours/train/mini_atlas_atac_paired.h5ad
    atac_key: X
    rna_path: /home/jwu418/workspace/data/ours/train/mini_atlas_rna_binned_binning_2_reduced.h5ad
    rna_key: X
  test:
    atac_path: /home/jwu418/workspace/data/ours/valid/mini_atlas_atac_paired.h5ad
    atac_key: X
    rna_path: /home/jwu418/workspace/data/ours/valid/mini_atlas_rna_binned_binning_2_reduced.h5ad
    rna_key: X

vocab:
  rna_path: /home/jwu418/workspace/data/ours/vocab/rna_vocab.json
  atac_path: /home/jwu418/workspace/data/ours/vocab/atac_vocab.json
  cell_type_path: /home/jwu418/workspace/data/ours/vocab/mini_atlas_cell_vocab.json
  batch_path: /home/jwu418/workspace/data/ours/vocab/mini_atlas_batch_vocab.json
  chr_path: /home/jwu418/workspace/data/ours/vocab/chr_vocab.json
  gene2chr_path: /home/jwu418/workspace/data/ours/vocab/gene2chr.json
  special_tokens:
    pad: {token: <pad>, value: 2}
    # value of the mask is 1 plus bin_num
    mask: {token: <mask>, value: 3}
    cls: {token: <cls>, value: 0}
```
