## Supplementary material for "EpiFoundation: A Foundation Model for Single-Cell ATAC-seq via Peak-to-Gene Alignment": Code for EpiFoundation: csv2h5ad.py

```
import scanpy as sc
import scipy
import os

def convert_dense_csv_to_sparse_h5ad(input_file, output_floder=None):
    
    file_name = input_file.split('/')[-1].split('.')[0]
    cell_type = file_name.split(':')[1]
    batch = file_name.split(':')[0]
    output_file = f'{output_floder}/{file_name}.h5ad'

    adata = sc.read_csv(input_file).T
    adata.X = scipy.sparse.csr_matrix(adata.X)
    # add a obs column, traverse all the cells, and add the cell name to the obs column
    adata.obs['cell_id'] = adata.obs_names
    # traverse all cells, get the biggest value of each cell, and add it to the obs column
    for i in range(adata.shape[0]):
        # max expression value of each cell, used for binning
        adata.obs['max_exp'] = adata.X[i].max() 
        adata.obs['non_zero'] = adata.X[i].count_nonzero()
        adata.obs['cell_type'] = cell_type
        # add the batch information to the obs column
        # currently, I treat name of (such asENCSR008NTI) as batch information
        # Cells from different batches may have a different meaning of same expression value
        adata.obs['batch'] = batch
        
    adata.var['gene_id'] = adata.var_names
    adata.uns['binned'] = False
    adata.uns['HVG'] = False
    adata.uns['log1p'] = False
    
    adata.write(output_file)
    return adata
```
