## Supplementary material for "EpiFoundation: A Foundation Model for Single-Cell ATAC-seq via Peak-to-Gene Alignment": Code for EpiFoundation: dataloader.py

```
import torch
import os
from torch.utils.data import  Dataset, DataLoader
from torchtext.vocab import Vocab
from scanpy import AnnData
from typing import Dict, List, Tuple, Union
import numpy as np
from memory_profiler import profile
from tokenizer.gene_tokenizer import GeneVocab
from tqdm import tqdm
import psutil
import scanpy as sc
import json

    
class ATACDataset(Dataset):
    def __init__(self, 
                atac_file,
                atac_key: str,
                atac_vocab: GeneVocab,
                cell_vocab: GeneVocab,
                batch_vocab: GeneVocab,
                chr_vocab: GeneVocab,
                atac_max_len: int,
                pad_token: str,
                cls_token: str,
                reg_token: str = None,
                logger = None,
    ):

        self.atac_key = atac_key if atac_key != "X" else None
        self.atac_max_len = atac_max_len
        self.logger = logger
        self.pad_token = pad_token
        self.cls_token = cls_token
        
        # read raw data
        self.rna_raw = {}
        self.atac_raw = {}
        
        
        self.log(f"load atac data from {atac_file}")
        atac_raw_data = sc.read_h5ad(atac_file, backed='r')
        self.atac_raw['values'] = atac_raw_data.layers[self.atac_key] if self.atac_key is not None else atac_raw_data.X
        try:
            gene_names = atac_raw_data.var['features'].tolist()
        except:
            gene_names = atac_raw_data.var_names.tolist()
        # get atac chr, by spliting the gene name using '-', and get the first element
        gene_chr = [gene.split('-')[0] for gene in gene_names]
        
        self.atac_raw['chr_ids'] = np.array(chr_vocab(gene_chr))
        self.atac_raw['gene_ids'] = np.array(atac_vocab(gene_names))
        self.atac_raw['pad_id'] = atac_vocab[pad_token]
        self.log(f"atac_raw: {self.atac_raw['values'].shape}")
        
        self.atac_cls_id= atac_vocab[cls_token]
        if reg_token is not None:
            self.atac_reg_id = atac_vocab[reg_token]
            self.chr_reg_id = chr_vocab[reg_token]
        else:
            self.atac_reg_id = None
            self.chr_reg_id = None
        self.chr_cls_id = chr_vocab[cls_token]
        self.chr_pad_id = chr_vocab[pad_token]
        
        self.cell_ids = cell_vocab(atac_raw_data.obs['annot'].tolist())
        self.batch_ids = batch_vocab(atac_raw_data.obs['batch'].tolist())
        return

        
    def __len__(self):
        return len(self.cell_ids)
    
    def log(self, msg):
        if int(os.environ["LOCAL_RANK"]) == 0:
            self.logger.info(msg)
    
    
    def prepare_row_atac(self, row):
        non_zero_idx = row.indices
        peaks = self.atac_raw['gene_ids'][non_zero_idx]
        chrs = self.atac_raw['chr_ids'][non_zero_idx]
        
        if self.atac_reg_id is not None:
            peaks = np.insert(peaks, 0, self.atac_reg_id)
        peaks = np.insert(peaks, 0, self.atac_cls_id)
        peaks = torch.from_numpy(peaks).long()
        
        if self.chr_reg_id is not None:
            chrs = np.insert(chrs, 0, self.chr_reg_id)
        chrs = np.insert(chrs, 0, self.chr_cls_id)
        chrs = torch.from_numpy(chrs).long()
        
        num_special_tokens = 1 if self.atac_reg_id is None else 2
        
        if len(peaks) > self.atac_max_len:
            idx = np.random.choice(len(peaks) - num_special_tokens, self.atac_max_len - num_special_tokens, replace=False)
            idx = idx + num_special_tokens
            for i in range(num_special_tokens):
                idx = np.insert(idx, i, i)
            peaks = peaks[idx]
            chrs = chrs[idx]
        elif len(peaks) <= self.atac_max_len:
            peaks = torch.cat(
                [
                    peaks,
                    torch.full(
                        (self.atac_max_len - len(peaks),), self.atac_raw['pad_id'], dtype=peaks.dtype
                    ),
                ]
            )
            chrs = torch.cat(
                [
                    chrs,
                    torch.full(
                        (self.atac_max_len - len(chrs),), self.chr_pad_id, dtype=chrs.dtype
                    ),
                ]
            )
        return peaks, chrs
    
    def __getitem__(self, idx):
        atac_row = self.atac_raw['values'][idx]
        atac_ids, atac_chrs = self.prepare_row_atac(atac_row)
        
        cell_id = torch.tensor(self.cell_ids[idx]).long()
        batch_id = torch.tensor(self.batch_ids[idx]).long()
        
        return {
            "atac_ids": atac_ids,
            "atac_chrs": atac_chrs,
            "cell_ids": cell_id,
            "batch_ids": batch_id,
        }
        
    
class PairedSCDataset(Dataset):
    def __init__(self, 
                rna_file,
                atac_file,
                rna_key: str,
                atac_key: str,
                rna_vocab: GeneVocab,
                atac_vocab: GeneVocab,
                cell_vocab: GeneVocab,
                batch_vocab: GeneVocab,
                chr_vocab: GeneVocab,
                gene2chr_file: str,
                rna_max_len: int,
                atac_max_len: int,
                pad_token: str,
                rna_pad_value: int,
                cls_token: str,
                reg_token: str = None,
                logger = None,
                get_full_genes: bool = False,
    ):
        self.rna_key = rna_key if rna_key != "X" else None
        self.atac_key = atac_key if atac_key != "X" else None
        self.rna_max_len = rna_max_len
        self.atac_max_len = atac_max_len
        self.logger = logger
        self.pad_token = pad_token
        self.rna_pad_value = rna_pad_value
        self.cls_token = cls_token
        
        # read raw data
        self.rna_raw = {}
        self.atac_raw = {}
        
        self.get_full_genes = get_full_genes
        
        

        self.log(f"load rna data from {rna_file}")
        rna_raw_data = sc.read_h5ad(rna_file, backed='r')
        self.rna_raw['values'] = rna_raw_data.layers[self.rna_key] if self.rna_key is not None else rna_raw_data.X
        try:
            gene_names = rna_raw_data.var['features'].tolist()
        except:
            gene_names = rna_raw_data.var_names.tolist()
            
        # read gene2chr file as a dict
        with open(gene2chr_file, 'r') as file:
            gene2chr = json.load(file)
        gene_chr = [gene2chr[gene] for gene in gene_names]
        
        self.rna_raw['chr_ids'] = np.array(chr_vocab(gene_chr))
        self.rna_raw['gene_ids'] = np.array(rna_vocab(gene_names))
        self.rna_raw['pad_id'] = rna_vocab[pad_token]
        self.log(f"rna_raw: {self.rna_raw['values'].shape}")
        
        
        
        self.log(f"load atac data from {atac_file}")
        atac_raw_data = sc.read_h5ad(atac_file, backed='r')
        self.atac_raw['values'] = atac_raw_data.layers[self.atac_key] if self.atac_key is not None else atac_raw_data.X
        try:
            gene_names = atac_raw_data.var['features'].tolist()
        except:
            gene_names = atac_raw_data.var_names.tolist()
        # get atac chr, by spliting the gene name using '-', and get the first element
        gene_chr = [gene.split('-')[0] for gene in gene_names]
        
        self.atac_raw['chr_ids'] = np.array(chr_vocab(gene_chr))
        self.atac_raw['gene_ids'] = np.array(atac_vocab(gene_names))
        self.atac_raw['pad_id'] = atac_vocab[pad_token]
        self.log(f"atac_raw: {self.atac_raw['values'].shape}")
        
        self.atac_cls_id= atac_vocab[cls_token]
        if reg_token is not None:
            self.atac_reg_id = atac_vocab[reg_token]
            self.chr_reg_id = chr_vocab[reg_token]
        else:
            self.atac_reg_id = None
            self.chr_reg_id = None
        self.chr_cls_id = chr_vocab[cls_token]
        self.chr_pad_id = chr_vocab[pad_token]
        
        rna_cell_idx = rna_raw_data.obs_names.tolist()
        atac_cell_idx = atac_raw_data.obs_names.tolist()

        assert (rna_cell_idx == atac_cell_idx)
        self.cell_ids = cell_vocab(atac_raw_data.obs['annot'].tolist())
        self.batch_ids = batch_vocab(atac_raw_data.obs['batch'].tolist())
        
        return

        
    def __len__(self):
        return len(self.cell_ids)
    
    def log(self, msg):
        if int(os.environ["LOCAL_RANK"]) == 0:
            self.logger.info(msg)
    
    def prepare_row_rna_full(self, row):
        if not isinstance(row, np.ndarray):
            row = row.toarray().flatten()
        genes = self.rna_raw['gene_ids']
        values = row
        chrs = self.rna_raw['chr_ids']
        
        genes = torch.from_numpy(genes).long()
        values = torch.from_numpy(values).long()
        chrs = torch.from_numpy(chrs).long()
        
        return genes, values, chrs
            
            
    def prepare_row_rna(self, row, non_zero_prob=0.5):
        # if row is not a numpy array, convert it to numpy array
        if not isinstance(row, np.ndarray):
            row = row.toarray().flatten()
        non_zero_idx = np.nonzero(row)[0]
        zero_idx = np.nonzero(row == 0)[0]
        
        total_number = min(self.rna_max_len, int(len(non_zero_idx) / non_zero_prob))
        non_zero_number = int(total_number * non_zero_prob)
        zero_number = total_number - non_zero_number
            
        non_zero_idx = np.random.choice(non_zero_idx, non_zero_number, replace=False)
        zero_idx = np.random.choice(zero_idx, zero_number, replace=False)
        
        non_zero_genes = self.rna_raw['gene_ids'][non_zero_idx]
        zero_genes = self.rna_raw['gene_ids'][zero_idx]
        
        chr_non_zero = self.rna_raw['chr_ids'][non_zero_idx]
        chr_zero = self.rna_raw['chr_ids'][zero_idx]
        
        genes = np.concatenate([non_zero_genes, zero_genes])
        values = np.concatenate([row[non_zero_idx], row[zero_idx]])
        chrs = np.concatenate([chr_non_zero, chr_zero])
        
        # shuffle the genes
        idx = np.random.permutation(len(genes))
        genes = genes[idx]
        values = values[idx]
        chrs = chrs[idx]
        
        genes = torch.from_numpy(genes).long()
        values = torch.from_numpy(values).long()
        chrs = torch.from_numpy(chrs).long()
        
        if len(genes) < self.rna_max_len:
            genes = torch.cat(
                [
                    genes,
                    torch.full(
                        (self.rna_max_len - len(genes),), self.rna_raw['pad_id'], dtype=genes.dtype
                    ),
                ]
            )
            values = torch.cat(
                [
                    values,
                    torch.full((self.rna_max_len - len(values),), self.rna_pad_value, dtype=values.dtype),
                ]
            )
            chrs = torch.cat(
                [
                    chrs,
                    torch.full((self.rna_max_len - len(chrs),), self.chr_pad_id, dtype=chrs.dtype),
                ]
            )
        return genes, values, chrs
    
    def prepare_row_rna_binary(self, row):
        non_zero_idx = np.nonzero(row)[0]
        zero_idx = np.nonzero(row == 0)[0]
        
        if len(non_zero_idx) > self.rna_max_len // 2:
            non_zero_number = self.rna_max_len // 2
            zero_number = self.rna_max_len - non_zero_number
        else:
            non_zero_number = len(non_zero_idx)
            zero_number = non_zero_number
            
        non_zero_idx = np.random.choice(non_zero_idx, non_zero_number, replace=False)
        zero_idx = np.random.choice(zero_idx, zero_number, replace=False)
        
        non_zero_genes = self.rna_raw['gene_ids'][non_zero_idx]
        zero_genes = self.rna_raw['gene_ids'][zero_idx]
        
        chr_non_zero = self.rna_raw['chr_ids'][non_zero_idx]
        chr_zero = self.rna_raw['chr_ids'][zero_idx]
        
        genes = np.concatenate([non_zero_genes, zero_genes])
        values = np.concatenate([np.ones(non_zero_number), np.zeros(zero_number)])
        chrs = np.concatenate([chr_non_zero, chr_zero])
        
        # shuffle the genes
        idx = np.random.permutation(len(genes))
        genes = genes[idx]
        values = values[idx]
        chrs = chrs[idx]
        
        genes = torch.from_numpy(genes).long()
        values = torch.from_numpy(values).long()
        chrs = torch.from_numpy(chrs).long()
        
        if len(genes) < self.rna_max_len:
            genes = torch.cat(
                [
                    genes,
                    torch.full(
                        (self.rna_max_len - len(genes),), self.rna_raw['pad_id'], dtype=genes.dtype
                    ),
                ]
            )
            values = torch.cat(
                [
                    values,
                    torch.full((self.rna_max_len - len(values),), self.rna_pad_value, dtype=values.dtype),
                ]
            )
            chrs = torch.cat(
                [
                    chrs,
                    torch.full((self.rna_max_len - len(chrs),), self.chr_pad_id, dtype=chrs.dtype),
                ]
            )
        return genes, values, chrs
    
    def prepare_row_atac(self, row):
        non_zero_idx = row.indices
        peaks = self.atac_raw['gene_ids'][non_zero_idx]
        chrs = self.atac_raw['chr_ids'][non_zero_idx]
        
        if self.atac_reg_id is not None:
            peaks = np.insert(peaks, 0, self.atac_reg_id)
        peaks = np.insert(peaks, 0, self.atac_cls_id)
        peaks = torch.from_numpy(peaks).long()
        
        if self.chr_reg_id is not None:
            chrs = np.insert(chrs, 0, self.chr_reg_id)
        chrs = np.insert(chrs, 0, self.chr_cls_id)
        chrs = torch.from_numpy(chrs).long()
        
        num_special_tokens = 1 if self.atac_reg_id is None else 2
        
        if len(peaks) > self.atac_max_len:
            idx = np.random.choice(len(peaks) - num_special_tokens, self.atac_max_len - num_special_tokens, replace=False)
            idx = idx + num_special_tokens
            for i in range(num_special_tokens):
                idx = np.insert(idx, i, i)
            peaks = peaks[idx]
            chrs = chrs[idx]
        elif len(peaks) <= self.atac_max_len:
            peaks = torch.cat(
                [
                    peaks,
                    torch.full(
                        (self.atac_max_len - len(peaks),), self.atac_raw['pad_id'], dtype=peaks.dtype
                    ),
                ]
            )
            chrs = torch.cat(
                [
                    chrs,
                    torch.full(
                        (self.atac_max_len - len(chrs),), self.chr_pad_id, dtype=chrs.dtype
                    ),
                ]
            )
        return peaks, chrs
    
    def __getitem__(self, idx):
        
        rna_row = self.rna_raw['values'][idx]
        if  self.get_full_genes == True:
            rna_ids, rna_values, rna_chrs = self.prepare_row_rna_full(rna_row)
        else:
            rna_ids, rna_values, rna_chrs = self.prepare_row_rna(rna_row)
        atac_row = self.atac_raw['values'][idx]
        atac_ids, atac_chrs = self.prepare_row_atac(atac_row)
        
        cell_id = torch.tensor(self.cell_ids[idx]).long()
        batch_id = torch.tensor(self.batch_ids[idx]).long()
        
        return {
            "rna_ids": rna_ids,
            "rna_values": rna_values,
            "rna_chrs": rna_chrs,
            "atac_ids": atac_ids,
            "atac_chrs": atac_chrs,
            "cell_ids": cell_id,
            "batch_ids": batch_id,
        }
```
