## Supplementary material for "EpiFoundation: A Foundation Model for Single-Cell ATAC-seq via Peak-to-Gene Alignment": Code for EpiFoundation: preprocess.py

```
from typing import Dict, Optional, Union

import numpy as np
import torch
from scipy.sparse import issparse
import scanpy as sc
from scanpy.get import _get_obs_rep, _set_obs_rep
from anndata import AnnData
from tqdm import tqdm
from model import logger


class Preprocessor:
    """
    Prepare data into training, valid and test split. Normalize raw expression
    values, binning or using other transform into the preset model input format.
    """

    def __init__(
        self,
        use_key: Optional[str] = None,
        filter_gene_by_counts: Union[int, bool] = False,
        filter_cell_by_counts: Union[int, bool] = False,
        normalize_total: Union[float, bool] = 1e4,
        result_normed_key: Optional[str] = "X_normed",
        log1p: bool = False,
        result_log1p_key: str = "X_log1p",
        subset_hvg: Union[int, bool] = False,
        hvg_use_key: Optional[str] = None,
        hvg_flavor: str = "seurat_v3",
        binning: Optional[int] = 6,
        result_binned_key: str = "X_binned",
    ):
        r"""
        Set up the preprocessor, use the args to config the workflow steps.

        Args:

        use_key (:class:`str`, optional):
            The key of :class:`~anndata.AnnData` to use for preprocessing.
        filter_gene_by_counts (:class:`int` or :class:`bool`, default: ``False``):
            Whther to filter genes by counts, if :class:`int`, filter genes with counts
        filter_cell_by_counts (:class:`int` or :class:`bool`, default: ``False``):
            Whther to filter cells by counts, if :class:`int`, filter cells with counts
        normalize_total (:class:`float` or :class:`bool`, default: ``1e4``):
            Whether to normalize the total counts of each cell to a specific value.
        result_normed_key (:class:`str`, default: ``"X_normed"``):
            The key of :class:`~anndata.AnnData` to store the normalized data. If
            :class:`None`, will use normed data to replce the :attr:`use_key`.
        log1p (:class:`bool`, default: ``True``):
            Whether to apply log1p transform to the normalized data.
        result_log1p_key (:class:`str`, default: ``"X_log1p"``):
            The key of :class:`~anndata.AnnData` to store the log1p transformed data.
        subset_hvg (:class:`int` or :class:`bool`, default: ``False``):
            Whether to subset highly variable genes.
        hvg_use_key (:class:`str`, optional):
            The key of :class:`~anndata.AnnData` to use for calculating highly variable
            genes. If :class:`None`, will use :attr:`adata.X`.
        hvg_flavor (:class:`str`, default: ``"seurat_v3"``):
            The flavor of highly variable genes selection. See
            :func:`scanpy.pp.highly_variable_genes` for more details.
        binning (:class:`int`, optional):
            Whether to bin the data into discrete values of number of bins provided.
        result_binned_key (:class:`str`, default: ``"X_binned"``):
            The key of :class:`~anndata.AnnData` to store the binned data.
        """
        self.use_key = use_key
        self.filter_gene_by_counts = filter_gene_by_counts
        self.filter_cell_by_counts = filter_cell_by_counts
        self.normalize_total = normalize_total
        self.result_normed_key = result_normed_key
        self.log1p = log1p
        self.result_log1p_key = result_log1p_key
        self.subset_hvg = subset_hvg
        self.hvg_use_key = hvg_use_key
        self.hvg_flavor = hvg_flavor
        self.binning = binning
        self.result_binned_key = result_binned_key

    def __call__(self, adata: AnnData, batch_key: Optional[str] = None) -> Dict:
        """
        format controls the different input value wrapping, including categorical
        binned style, fixed-sum normalized counts, log1p fixed-sum normalized counts, etc.

        Args:

        adata (:class:`AnnData`):
            The :class:`AnnData` object to preprocess.
        batch_key (:class:`str`, optional):
            The key of :class:`AnnData.obs` to use for batch information. This arg
            is used in the highly variable gene selection step.
        """
        key_to_process = self.use_key
        # preliminary checks, will use later
        if key_to_process == "X":
            key_to_process = None  # the following scanpy apis use arg None to use X
        is_logged = self.check_logged(adata, obs_key=key_to_process)

        # step 1: filter genes
        if self.filter_gene_by_counts:
            logger.info("Filtering genes by counts ...")
            sc.pp.filter_genes(
                adata,
                min_counts=self.filter_gene_by_counts
                if isinstance(self.filter_gene_by_counts, int)
                else None,
            )

        # step 2: filter cells
        if (
            isinstance(self.filter_cell_by_counts, int)
            and self.filter_cell_by_counts > 0
        ):
            logger.info("Filtering cells by counts ...")
            sc.pp.filter_cells(
                adata,
                min_counts=self.filter_cell_by_counts
                if isinstance(self.filter_cell_by_counts, int)
                else None,
            )

        # step 3: normalize total
        if self.normalize_total:
            logger.info("Normalizing total counts ...")
            normed_ = sc.pp.normalize_total(
                adata,
                target_sum=self.normalize_total
                if isinstance(self.normalize_total, float)
                else None,
                layer=key_to_process,
                inplace=False,
            )["X"]
            key_to_process = self.result_normed_key or key_to_process
            _set_obs_rep(adata, normed_, layer=key_to_process)

        # step 4: log1p
        if self.log1p:
            logger.info("Log1p transforming ...")
            if is_logged:
                logger.warning(
                    "The input data seems to be already log1p transformed. "
                    "Set `log1p=False` to avoid double log1p transform."
                )
            if self.result_log1p_key:
                _set_obs_rep(
                    adata,
                    _get_obs_rep(adata, layer=key_to_process),
                    layer=self.result_log1p_key,
                )
                key_to_process = self.result_log1p_key
            sc.pp.log1p(adata, layer=key_to_process)


        # Select HVG in each batch if batch_key is provided
        # step 5: subset hvg
        if self.subset_hvg:
            logger.info("Subsetting highly variable genes ...")
            if batch_key is None:
                logger.warning(
                    "No batch_key is provided, will use all cells for HVG selection."
                )
            sc.pp.highly_variable_genes(
                adata,
                layer=self.hvg_use_key,
                n_top_genes=self.subset_hvg
                if isinstance(self.subset_hvg, int)
                else None,
                batch_key=batch_key,
                flavor=self.hvg_flavor,
                subset=True,
            )

        # step 6: binning
        if self.binning:
            logger.info("Binning data with {} bins ...".format(self.binning))
            if isinstance(self.binning, int):
                # raise ValueError(
                #     "Binning arg must be an integer, but got {}.".format(self.binning)
                # )
                n_bins = self.binning  # NOTE: the first bin is always a spectial for zero
                binned_rows = []
                bin_edges = []
                # layer to be preprocessed refer to unbinned data, such as X, X_pca, X_umap, etc.
                layer_data = _get_obs_rep(adata, layer=key_to_process)
                layer_data = layer_data.A if issparse(layer_data) else layer_data
                if layer_data.min() < 0:
                    raise ValueError(
                        f"Assuming non-negative data, but got min value {layer_data.min()}."
                    )
                for row in tqdm(layer_data):
                    if row.max() == 0:
                        logger.warning(
                            "The input data contains all zero rows. Please make sure "
                            "this is expected. You can use the `filter_cell_by_counts` "
                            "arg to filter out all zero rows."
                        )
                        binned_rows.append(np.zeros_like(row, dtype=np.int64))
                        bin_edges.append(np.array([0] * n_bins))
                        continue
                    non_zero_ids = row.nonzero()
                    non_zero_row = row[non_zero_ids]
                    # generate bins, represent a series of bin edges
                    # np.linspace(0, 1, n_bins - 1) is the cumulative distribution function
                    # return the quantile of the data, which is the inverse of the cumulative
                    bins = np.quantile(non_zero_row, np.linspace(0, 1, n_bins - 1))
                    # bins = np.sort(np.unique(bins))
                    # NOTE: comment this line for now, since this will make the each category
                    # has different relative meaning across datasets
                    non_zero_digits = _digitize(non_zero_row, bins, side='one')
                    assert non_zero_digits.min() >= 1
                    assert non_zero_digits.max() <= n_bins - 1
                    binned_row = np.zeros_like(row, dtype=np.int64)
                    binned_row[non_zero_ids] = non_zero_digits
                    binned_rows.append(binned_row)
                    bin_edges.append(np.concatenate([[0], bins]))
                    
                # Layer refer to the data type, such as X, X_pca, X_umap, etc.
                adata.layers[self.result_binned_key] = np.stack(binned_rows)
                adata.obsm["bin_edges"] = np.stack(bin_edges)
            elif isinstance(self.binning, list):
                # layer to be preprocessed refer to unbinned data, such as X, X_pca, X_umap, etc.
                layer_data = _get_obs_rep(adata, layer=key_to_process)
                layer_data = layer_data.A if issparse(layer_data) else layer_data
                if layer_data.min() < 0:
                    raise ValueError(
                        f"Assuming non-negative data, but got min value {layer_data.min()}."
                    )
                for bin_number in self.binning:
                    n_bins = bin_number
                    binned_rows = []
                    bin_edges = []
                    logger.info(f"Processing binning with {n_bins} bins")
                    for row in tqdm(layer_data):
                        if row.max() == 0:
                            logger.warning(
                                "The input data contains all zero rows. Please make sure "
                                "this is expected. You can use the `filter_cell_by_counts` "
                                "arg to filter out all zero rows."
                            )
                            binned_rows.append(np.zeros_like(row, dtype=np.int64))
                            bin_edges.append(np.array([0] * n_bins))
                            continue
                        non_zero_ids = row.nonzero()
                        non_zero_row = row[non_zero_ids]
                        # generate bins, represent a series of bin edges
                        # np.linspace(0, 1, n_bins - 1) is the cumulative distribution function
                        # return the quantile of the data, which is the inverse of the cumulative
                        bins = np.quantile(non_zero_row, np.linspace(0, 1, n_bins - 1))
                        # bins = np.sort(np.unique(bins))
                        # NOTE: comment this line for now, since this will make the each category
                        # has different relative meaning across datasets
                        non_zero_digits = _digitize(non_zero_row, bins, side='one')
                        assert non_zero_digits.min() >= 1
                        assert non_zero_digits.max() <= n_bins - 1
                        binned_row = np.zeros_like(row, dtype=np.int64)
                        binned_row[non_zero_ids] = non_zero_digits
                        binned_rows.append(binned_row)
                        bin_edges.append(np.concatenate([[0], bins]))
                    result_key = self.result_binned_key
                    bin_edges_key = n_bins
                    logger.info(f"Saving binned data with {n_bins} bins, result_key: {result_key}")
                    adata.layers[result_key] = np.stack(binned_rows)
                    adata.obsm[bin_edges_key] = np.stack(bin_edges)

    def check_logged(self, adata: AnnData, obs_key: Optional[str] = None) -> bool:
        """
        Check if the data is already log1p transformed.

        Args:

        adata (:class:`AnnData`):
            The :class:`AnnData` object to preprocess.
        obs_key (:class:`str`, optional):
            The key of :class:`AnnData.obs` to use for batch information. This arg
            is used in the highly variable gene selection step.
        """
        data = _get_obs_rep(adata, layer=obs_key)
        max_, min_ = data.max(), data.min()
        if max_ > 30:
            return False
        if min_ < 0:
            return False

        non_zero_min = data[data > 0].min()
        if non_zero_min >= 1:
            return False

        return True


def _digitize(x: np.ndarray, bins: np.ndarray, side="both") -> np.ndarray:
    """
    Digitize the data into bins. This method spreads data uniformly when bins
    have same values.

    Args:

    x (:class:`np.ndarray`):
        The data to digitize.
    bins (:class:`np.ndarray`):
        The bins to use for digitization, in increasing order.
    side (:class:`str`, optional):
        The side to use for digitization. If "one", the left side is used. If
        "both", the left and right side are used. Default to "one".

    Returns:

    :class:`np.ndarray`:
        The digitized data.
    """
    assert x.ndim == 1 and bins.ndim == 1

    left_digits = np.digitize(x, bins)
    if side == "one":
        return left_digits

    right_difits = np.digitize(x, bins, right=True)

    rands = np.random.rand(len(x))  # uniform random numbers

    digits = rands * (right_difits - left_digits) + left_digits
    digits = np.ceil(digits).astype(np.int64)
    return digits


def binning(
    row: Union[np.ndarray, torch.Tensor], n_bins: int
) -> Union[np.ndarray, torch.Tensor]:
    """Binning the row into n_bins."""
    dtype = row.dtype
    return_np = False if isinstance(row, torch.Tensor) else True
    row = row.cpu().numpy() if isinstance(row, torch.Tensor) else row
    # TODO: use torch.quantile and torch.bucketize

    if row.max() == 0:
        logger.warning(
            "The input data contains row of zeros. Please make sure this is expected."
        )
        return (
            np.zeros_like(row, dtype=dtype)
            if return_np
            else torch.zeros_like(row, dtype=dtype)
        )

    if row.min() <= 0:
        non_zero_ids = row.nonzero()
        non_zero_row = row[non_zero_ids]
        bins = np.quantile(non_zero_row, np.linspace(0, 1, n_bins - 1))
        non_zero_digits = _digitize(non_zero_row, bins)
        binned_row = np.zeros_like(row, dtype=np.int64)
        binned_row[non_zero_ids] = non_zero_digits
    else:
        bins = np.quantile(row, np.linspace(0, 1, n_bins - 1))
        binned_row = _digitize(row, bins)
    return torch.from_numpy(binned_row) if not return_np else binned_row.astype(dtype)
```
