## Supplementary material for "EpiFoundation: A Foundation Model for Single-Cell ATAC-seq via Peak-to-Gene Alignment": Code for EpiFoundation: env.yml

```
name: epiFoundation
channels:
  - pytorch
  - nvidia
  - defaults
  - conda-forge
  - https://repo.anaconda.com/pkgs/main
  - https://repo.anaconda.com/pkgs/r
dependencies:
  - _libgcc_mutex=0.1=main
  - _openmp_mutex=5.1=1_gnu
  - absl-py=2.1.0=pyhd8ed1ab_0
  - aom=3.6.0=h6a678d5_0
  - blas=1.0=mkl
  - bleach=4.1.0=pyhd3eb1b0_0
  - blosc=1.21.3=h6a678d5_0
  - bokeh=3.1.1=py38h2f386ee_0
  - bottleneck=1.3.7=py38ha9d4c09_0
  - brotli=1.0.9=h5eee18b_8
  - brotli-bin=1.0.9=h5eee18b_8
  - brotli-python=1.0.9=py38h6a678d5_8
  - brunsli=0.1=h2531618_0
  - bzip2=1.0.8=h5eee18b_6
  - c-ares=1.19.1=h5eee18b_0
  - ca-certificates=2024.9.24=h06a4308_0
  - certifi=2024.8.30=py38h06a4308_0
  - cfitsio=3.470=h5893167_7
  - charls=2.2.0=h2531618_0
  - charset-normalizer=3.3.2=pyhd3eb1b0_0
  - click=8.1.7=py38h06a4308_0
  - cloudpickle=3.0.0=py38h06a4308_0
  - colorama=0.4.6=pyhd8ed1ab_0
  - colorcet=3.1.0=py38h06a4308_0
  - cuda-cudart=12.1.105=0
  - cuda-cupti=12.1.105=0
  - cuda-libraries=12.1.0=0
  - cuda-nvrtc=12.1.105=0
  - cuda-nvtx=12.1.105=0
  - cuda-opencl=12.6.68=0
  - cuda-runtime=12.1.0=0
  - cuda-version=12.6=3
  - cycler=0.12.1=pyhd8ed1ab_0
  - cyrus-sasl=2.1.28=h52b45da_1
  - cytoolz=0.12.2=py38h5eee18b_0
  - dask-core=2023.4.1=py38h06a4308_0
  - datashader=0.15.2=py38h06a4308_0
  - datashape=0.5.4=py38h06a4308_1
  - dav1d=1.2.1=h5eee18b_0
  - dbus=1.13.18=hb2f20db_0
  - expat=2.6.3=h6a678d5_0
  - ffmpeg=4.3=hf484d3e_0
  - fontconfig=2.14.1=h55d465d_3
  - freetype=2.12.1=h4a9f257_0
  - giflib=5.2.1=h5eee18b_3
  - glib=2.78.4=h6a678d5_0
  - glib-tools=2.78.4=h6a678d5_0
  - gmp=6.2.1=h295c915_3
  - gmpy2=2.1.2=py38heeb90bb_0
  - gnutls=3.6.15=he1e5248_0
  - grpcio=1.62.2=py38h6a678d5_0
  - gst-plugins-base=1.14.1=h6a678d5_1
  - gstreamer=1.14.1=h5eee18b_1
  - holoviews=1.17.1=py38h06a4308_0
  - icu=73.1=h6a678d5_0
  - imagecodecs=2023.1.23=py38hc4b7b5f_0
  - imageio=2.33.1=py38h06a4308_0
  - importlib_metadata=8.5.0=hd8ed1ab_0
  - importlib_resources=6.4.0=py38h06a4308_0
  - intel-openmp=2023.1.0=hdb19cb5_46306
  - jinja2=3.1.4=py38h06a4308_0
  - joblib=1.4.2=py38h06a4308_0
  - jpeg=9e=h5eee18b_3
  - jxrlib=1.1=h7b6447c_2
  - krb5=1.20.1=h143b758_1
  - lame=3.100=h7b6447c_0
  - lazy_loader=0.4=py38h06a4308_0
  - lcms2=2.12=h3be6417_0
  - ld_impl_linux-64=2.38=h1181459_1
  - lerc=3.0=h295c915_0
  - libabseil=20240116.2=cxx17_h6a678d5_0
  - libaec=1.0.4=he6710b0_1
  - libavif=0.11.1=h5eee18b_0
  - libblas=3.9.0=1_h86c2bf4_netlib
  - libbrotlicommon=1.0.9=h5eee18b_8
  - libbrotlidec=1.0.9=h5eee18b_8
  - libbrotlienc=1.0.9=h5eee18b_8
  - libcblas=3.9.0=5_h92ddd45_netlib
  - libclang=14.0.6=default_hc6dbbc7_1
  - libclang13=14.0.6=default_he11475f_1
  - libcublas=12.1.0.26=0
  - libcufft=11.0.2.4=0
  - libcufile=1.11.1.6=0
  - libcups=2.4.2=h2d74bed_1
  - libcurand=10.3.7.68=0
  - libcurl=8.9.1=h251f7ec_0
  - libcusolver=11.4.4.55=0
  - libcusparse=12.0.2.55=0
  - libdeflate=1.17=h5eee18b_1
  - libedit=3.1.20230828=h5eee18b_0
  - libev=4.33=h7f8727e_1
  - libffi=3.4.4=h6a678d5_1
  - libgcc-ng=11.2.0=h1234567_1
  - libgfortran-ng=13.2.0=h69a702a_0
  - libgfortran5=13.2.0=ha4646dd_0
  - libglib=2.78.4=hdc74915_0
  - libgomp=11.2.0=h1234567_1
  - libgrpc=1.62.2=h2d74bed_0
  - libiconv=1.16=h5eee18b_3
  - libidn2=2.3.4=h5eee18b_0
  - libjpeg-turbo=2.0.0=h9bf148f_0
  - liblapack=3.9.0=5_h92ddd45_netlib
  - libllvm14=14.0.6=hecde1de_4
  - libnghttp2=1.57.0=h2d74bed_0
  - libnpp=12.0.2.50=0
  - libnvjitlink=12.1.105=0
  - libnvjpeg=12.1.1.14=0
  - libpng=1.6.39=h5eee18b_0
  - libpq=12.17=hdbd6064_0
  - libprotobuf=4.25.3=he621ea3_0
  - libssh2=1.11.0=h251f7ec_0
  - libstdcxx-ng=11.2.0=h1234567_1
  - libtasn1=4.19.0=h5eee18b_0
  - libtiff=4.5.1=h6a678d5_0
  - libunistring=0.9.10=h27cfd23_0
  - libuuid=1.41.5=h5eee18b_0
  - libwebp-base=1.3.2=h5eee18b_0
  - libxcb=1.15=h7f8727e_0
  - libxkbcommon=1.0.1=h097e994_2
  - libxml2=2.13.1=hfdd30dd_2
  - libzopfli=1.0.3=he6710b0_0
  - linkify-it-py=2.0.0=py38h06a4308_0
  - llvm-openmp=14.0.6=h9e868ea_0
  - locket=1.0.0=py38h06a4308_0
  - lz4-c=1.9.4=h6a678d5_1
  - markdown=3.6=pyhd8ed1ab_0
  - markdown-it-py=2.2.0=py38h06a4308_1
  - markupsafe=2.1.3=py38h5eee18b_0
  - mdit-py-plugins=0.3.0=py38h06a4308_0
  - mdurl=0.1.0=py38h06a4308_0
  - mkl=2023.1.0=h213fc3f_46344
  - mkl-service=2.4.0=py38h5eee18b_1
  - mkl_fft=1.3.8=py38h5eee18b_0
  - mkl_random=1.2.4=py38hdb19cb5_0
  - mpc=1.1.0=h10f8cd9_1
  - mpfr=4.0.2=hb69a4c5_1
  - mpmath=1.3.0=py38h06a4308_0
  - multipledispatch=0.6.0=py38_0
  - munkres=1.1.4=pyh9f0ad1d_0
  - mysql=5.7.24=h721c034_2
  - ncurses=6.4=h6a678d5_0
  - nettle=3.7.3=hbbd107a_1
  - networkx=3.1=py38h06a4308_0
  - numba=0.58.1=py38h6a678d5_0
  - numpy=1.24.3=py38hf6e8229_1
  - numpy-base=1.24.3=py38h060ed82_1
  - openh264=2.1.1=h4ff587b_0
  - openjpeg=2.5.2=he7f1fd0_0
  - openssl=3.0.15=h5eee18b_0
  - packaging=24.1=pyhd8ed1ab_0
  - pandas=2.0.3=py38h1128e8f_0
  - panel=1.2.3=py38h06a4308_0
  - param=1.13.0=py38h06a4308_0
  - partd=1.4.1=py38h06a4308_0
  - pcre2=10.42=hebb0a14_1
  - pillow=10.4.0=py38h5eee18b_0
  - pip=24.2=py38h06a4308_0
  - ply=3.11=py38_0
  - pyct=0.5.0=py38h06a4308_0
  - pynndescent=0.5.13=pyhff2d567_0
  - pyqt=5.15.10=py38h6a678d5_0
  - pyqt5-sip=12.13.0=py38h5eee18b_0
  - pysocks=1.7.1=py38h06a4308_0
  - python=3.8.19=h955ad1f_0
  - python-dateutil=2.9.0post0=py38h06a4308_2
  - python_abi=3.8=2_cp38
  - pytorch=2.1.1=py3.8_cuda12.1_cudnn8.9.2_0
  - pytorch-cuda=12.1=ha16c6d3_5
  - pytorch-mutex=1.0=cuda
  - pytz=2024.1=py38h06a4308_0
  - pyviz_comms=3.0.2=py38h06a4308_0
  - pywavelets=1.4.1=py38h5eee18b_0
  - pyyaml=6.0.1=py38h5eee18b_0
  - qt-main=5.15.2=h53bd1ea_10
  - re2=2022.04.01=h27087fc_0
  - readline=8.2=h5eee18b_0
  - requests=2.32.3=py38h06a4308_0
  - scikit-image=0.20.0=py38h6a678d5_0
  - scipy=1.8.1=py38h1ee437e_0
  - setuptools=72.1.0=py38h06a4308_0
  - sip=6.7.12=py38h6a678d5_0
  - six=1.16.0=pyh6c4a22f_0
  - snappy=1.2.1=h6a678d5_0
  - sqlite=3.45.3=h5eee18b_0
  - sympy=1.13.2=py38h06a4308_0
  - tbb=2021.8.0=hdb19cb5_0
  - tensorboard=2.17.1=pyhd8ed1ab_0
  - tensorboard-data-server=0.7.0=py38h52d8a92_1
  - threadpoolctl=3.5.0=pyhc1e730c_0
  - tifffile=2023.4.12=py38h06a4308_0
  - tk=8.6.14=h39e8969_0
  - tomli=2.0.1=py38h06a4308_0
  - toolz=0.12.0=py38h06a4308_0
  - torchaudio=2.1.1=py38_cu121
  - torchtriton=2.1.0=py38
  - torchvision=0.16.1=py38_cu121
  - tornado=6.4.1=py38h5eee18b_0
  - tqdm=4.66.5=pyhd8ed1ab_0
  - uc-micro-py=1.0.1=py38h06a4308_0
  - umap-learn=0.5.6=py38h578d9bd_1
  - unicodedata2=15.1.0=py38h5eee18b_0
  - urllib3=2.2.2=py38h06a4308_0
  - webencodings=0.5.1=py38_1
  - werkzeug=3.0.4=pyhd8ed1ab_0
  - wheel=0.43.0=py38h06a4308_0
  - xarray=2022.11.0=py38h06a4308_0
  - xyzservices=2022.9.0=py38h06a4308_1
  - xz=5.4.6=h5eee18b_1
  - yaml=0.2.5=h7b6447c_0
  - zfp=1.0.0=h6a678d5_0
  - zlib=1.2.13=h5eee18b_1
  - zstd=1.5.5=hc292b87_2
  - pip:
      - aiohappyeyeballs==2.4.0
      - aiohttp==3.10.5
      - aiosignal==1.3.1
      - anndata==0.9.2
      - async-timeout==4.0.3
      - attrs==24.2.0
      - beautifulsoup4==4.12.3
      - blessed==1.20.0
      - blosc2==2.0.0
      - contourpy==1.1.1
      - cython==3.0.11
      - deprecated==1.2.14
      - einops==0.8.0
      - filelock==3.15.4
      - flash-attn==2.6.3
      - fonttools==4.53.1
      - frozenlist==1.4.1
      - fsspec==2024.9.0
      - gdown==5.2.0
      - get-annotations==0.1.2
      - gpustat==1.1.1
      - h5py==3.11.0
      - huggingface-hub==0.0.8
      - idna==3.8
      - igraph==0.11.8
      - importlib-metadata==8.4.0
      - kiwisolver==1.4.7
      - legacy-api-wrap==1.4
      - leidenalg==0.10.2
      - llvmlite==0.41.1
      - local-attention==1.9.14
      - louvain==0.8.2
      - matplotlib==3.6.3
      - memory-profiler==0.61.0
      - msgpack==1.0.8
      - multidict==6.0.5
      - natsort==8.4.0
      - ninja==1.11.1.1
      - numexpr==2.8.6
      - nvidia-ml-py==12.560.30
      - patsy==0.5.6
      - protobuf==5.28.2
      - psutil==6.0.0
      - py-cpuinfo==9.0.0
      - pydot==3.0.2
      - pyparsing==3.1.4
      - regex==2024.7.24
      - sacremoses==0.1.1
      - scanpy==1.9.8
      - scib==1.1.5
      - scikit-learn==0.24.2
      - scikit-misc==0.2.0
      - seaborn==0.13.2
      - session-info==1.0.0
      - sinfo==0.3.4
      - soupsieve==2.6
      - statsmodels==0.14.1
      - stdlib-list==0.10.0
      - tables==3.8.0
      - tensorboardx==2.6.2.2
      - texttable==1.7.0
      - tokenizers==0.10.3
      - torchdata==0.7.1
      - torchtext==0.16.1
      - transformers==4.6.1
      - typing-extensions==4.12.2
      - tzdata==2024.1
      - wcwidth==0.2.13
      - wrapt==1.16.0
      - yarl==1.9.10
      - zipp==3.20.1
prefix: /your/conda/envs/epiFoundation
```
