## Supplementary material for "EpiFoundation: A Foundation Model for Single-Cell ATAC-seq via Peak-to-Gene Alignment": Code for EpiFoundation: eval.py

```
# -*- coding: utf-8 -*-

import os
import gc
import argparse
import json
import random
import math
import random
from functools import reduce
import numpy as np
import pandas as pd
from scipy import sparse
from sklearn.model_selection import train_test_split
import torch
from torch import nn
from torch.optim import Adam
from torch.nn import functional as F
from tensorboardX import SummaryWriter
from torch.utils.data import DataLoader, Dataset
from torch.utils.data.distributed import DistributedSampler
from torch.nn.parallel import DistributedDataParallel as DDP
import torch.distributed as dist
from model import EpiFoundation
from loss.loss import MaskedMSELoss
from data.dataloader import *
from tokenizer import GeneVocab
import scanpy as sc
import anndata as ad
from utils import *
from memory_profiler import profile

import yaml

torch.autograd.set_detect_anomaly(True)

parser = argparse.ArgumentParser()
parser.add_argument("--config", type=str, default='./configs/eval/baseline.yml', help='Config file.')
parser.add_argument("--backend", type=str, default='flash', help='Fast Transformer backend.')
args = parser.parse_args()

# @profile(precision=4, stream=open("memory_profiler.log", "w+"))
def main():
    # read and parse config file
    local_rank = int(os.environ["LOCAL_RANK"])
    with open(args.config, 'r') as f:
        config = yaml.load(f, Loader=yaml.FullLoader)
        
    
    train_config = config['train']
    valid_config = config['valid']
    data_config = config['data']
    vocab_config = config['vocab']
    task_name = config['task_name']
    task_floder = './result/{}'.format(task_name)
    ckpt_dir = os.path.join(task_floder, 'ckpts')
    

    
    random_seed = train_config['seed']
    EPOCHS = train_config['epochs']
    BATCH_SIZE = train_config['batch_size']
    GRADIENT_ACCUMULATION = train_config['gradient_accumulation_steps']
    LEARNING_RATE = float(train_config['lr'])

    model_name = train_config['model']['encoder']
    
    save_ckpt_freq = train_config['save_ckpt_freq'] if 'save_ckpt_freq' in train_config else 5
    resume = train_config['resume'] if 'resume' in train_config else False
    
    # special tokens
    pad = vocab_config['special_tokens']['pad']
    mask = vocab_config['special_tokens']['mask']
    cls = vocab_config['special_tokens']['cls']
    
    # distibuted setting
    dist.init_process_group(backend='nccl')
    torch.cuda.set_device(local_rank)
    device = torch.device("cuda", local_rank)
    world_size = torch.distributed.get_world_size()
    seed_all(random_seed + torch.distributed.get_rank())
    is_master = (local_rank == 0)
    
    # init loggers
    logger = set_log(log_dir= os.path.join(task_floder, 'logs'))
    tb_logger = SummaryWriter(os.path.join(task_floder, 'tb_logs'))
    if is_master:
        logger.info(dict2str(config))
    
    
    rna_vocab = GeneVocab.from_file(vocab_config['rna_path'])
    atac_vocab = GeneVocab.from_file(vocab_config['atac_path'])
    cell_vocab = GeneVocab.from_file(vocab_config['cell_type_path'])
    batch_vocab = GeneVocab.from_file(vocab_config['batch_path'])
    chr_vocab = GeneVocab.from_file(vocab_config['chr_path'])
    if is_master:
        logger.info(f'Rna vocab size: {len(rna_vocab)}')
        logger.info(f'Atac vocab size: {len(atac_vocab)}')
        
    if is_master:
        logger.info('loading training data')


    
    if is_master:
        logger.info('loading validation data')
    val_set = PairedSCDataset(
        rna_file = data_config['test']['rna_path'],
        atac_file= data_config['test']['atac_path'],
        rna_key = data_config['test']['rna_key'],
        atac_key = data_config['test']['atac_key'],
        rna_vocab = rna_vocab,
        atac_vocab = atac_vocab,
        cell_vocab = cell_vocab,
        batch_vocab= batch_vocab,
        chr_vocab = chr_vocab,
        gene2chr_file= vocab_config['gene2chr_path'],
        rna_max_len = train_config['model']['rna_max_len'],
        atac_max_len = train_config['model']['atac_max_len'],
        pad_token = pad['token'],
        rna_pad_value = pad['value'],
        cls_token = cls['token'],
        logger = logger,
    )
    gc.collect()

    val_sampler = SequentialDistributedSampler(val_set, batch_size=BATCH_SIZE, world_size=world_size)
    val_loader = DataLoader(val_set, batch_size=BATCH_SIZE, sampler=val_sampler, prefetch_factor=4, num_workers=4)
    # create non distributed loader for evaluation
    if is_master:
        # for evaluation, batch size should be 1
        val_non_dist_loader = DataLoader(val_set, batch_size= 1, shuffle=False, num_workers=4) 
    
    if is_master:
        logger.info('Creating model')
    
    model = EpiFoundation(
        num_class_cell = len(cell_vocab),
        num_rnas = len(rna_vocab),
        num_atacs = len(atac_vocab),
        num_values= data_config['bin_num'],
        num_chrs= len(chr_vocab),
        embed_dim = train_config['model']['embedding_dim'],
        depth = train_config['model']['num_layers'],
        heads = train_config['model']['head_num'],
        head_dim = train_config['model']['head_dim'],
        encoder = model_name,
        dropout = train_config['model']['dropout'],
        pad_token_idx_rna = rna_vocab[pad['token']],
        pad_token_idx_atac = atac_vocab[pad['token']],
        cell_emb_style = train_config['model']['cell_emb_style'],
        mvc_arch_style = train_config['model']['mvc_arch_style'],
        use_batch_labels = train_config['model']['use_batch_labels'],
        batch_label_num= len(batch_vocab),
        use_chr_labels= train_config['model']['use_chr_labels'],
        transformer_backend = args.backend,
    ).to(device)
    
    # optimizer
    optimizer = Adam(model.parameters(), lr=LEARNING_RATE)
    
    # learning rate scheduler
    scheduler = CosineAnnealingWarmupRestarts(
        optimizer,
        first_cycle_steps=15,
        cycle_mult=2,
        max_lr=LEARNING_RATE,
        min_lr=1e-6,
        warmup_steps=5,
        gamma=0.9
    )
    
    if is_master and train_config['metric'] == True:
        non_dist_model = model
    model = DDP(model, device_ids=[local_rank], output_device=local_rank)
    
    # scaler = torch.amp.GradScaler(enabled=train_config['amp'].amp)
    scaler = torch.cuda.amp.GradScaler(enabled=train_config['amp'])
    
    # masked_mse_loss = MaskedMSELoss().to(local_rank)
    cross_entropy_loss = nn.CrossEntropyLoss(reduction='mean').to(local_rank)
    atac_cross_entropy_loss = nn.CrossEntropyLoss(reduction='mean', ignore_index = pad['value']).to(local_rank)
    
    softmax = nn.Softmax(dim=-1)
    
    steps = 0
    if train_config['model']['pretrained'] is not None:
        if is_master:
            logger.info('Loading pretrained model from: {}'.format(train_config['model']['pretrained']))
        checkpoint = torch.load(train_config['model']['pretrained'], map_location=device)
        
        # # do not load batch_emb and cls_decoder parameters (when finetuning on different dataset)
        pretrained_dict = {k: v for k, v in checkpoint['model'].items() if 'batch_emb' not in k and 'cls_decoder' not in k and 'mvc_decoder' not in k}
        # pretrained_dict = {k: v for k, v in checkpoint['model'].items() if 'batch_emb' not in k }
        model_dict = model.module.state_dict()
        model_dict.update(pretrained_dict)
        model.module.load_state_dict(model_dict)
        if is_master and train_config['metric'] == True:
            non_dist_model.load_state_dict(checkpoint['model'])
        # optimizer.load_state_dict(checkpoint['optimizer'])
        scheduler.load_state_dict(checkpoint['scheduler'])
        scaler.load_state_dict(checkpoint['scaler'])
        del checkpoint
        gc.collect()
    
    dist.barrier()
    
    if train_config['metric'] == True:
        if is_master:
            non_dist_model.eval()
            model.eval()
            logger.info('Start evaluation with scib metrices')
            test_adata = sc.read_h5ad(data_config['test']['atac_path'])
            
            batch_labels = []
            embeddings = []
            cell_labels = []
            cell_pred = []
            
            cell_acc = 0.0
            tbar = tqdm(val_non_dist_loader, desc='Eval')
            for index, batch in enumerate(val_non_dist_loader):
                tbar.update(1)
                index += 1
                # if index > 10:
                #     break
                
                rna_values = batch['rna_values'].to(device)
                rna_ids = batch['rna_ids'].to(device)
                atac_ids = batch['atac_ids'].to(device)
                cell_ids = batch['cell_ids'].to(device)
                batch_ids = batch['batch_ids'].to(device)
                rna_chrs = batch['rna_chrs'].to(device)
                atac_chrs = batch['atac_chrs'].to(device)
                
                padding_positions = atac_ids.eq(atac_vocab[pad['token']])
                with torch.cuda.amp.autocast(enabled=train_config['amp'], dtype= torch.bfloat16):
                    output = non_dist_model(atac = atac_ids, rna = rna_ids, src_key_padding_mask = padding_positions, batch_id = batch_ids, rna_chrs = rna_chrs, atac_chrs = atac_chrs)
                pred = softmax(output['cell_pred']).argmax(dim=-1)
                cell_pred.append(pred.cpu().numpy())
                
                cell_labels.append(cell_ids.cpu().numpy())
                
                # print("GT and Pred: ", cell_ids.cpu().numpy(), pred.cpu().numpy())
                
                embeddings.append(output['cell_emb'].detach().cpu().numpy())
                batch_labels.append(batch_ids.cpu().numpy())
            tbar.close()
            # concatenate the results and transform to numpy array
            cell_labels = np.concatenate(cell_labels)
            embeddings = np.concatenate(embeddings)
            batch_labels = np.concatenate(batch_labels)
            
            
            embeddings = embeddings / np.linalg.norm(embeddings, axis=1, keepdims=True)
            
            data_cell_ids = np.array(cell_vocab(test_adata.obs['annot'].tolist()))
            data_batch_ids = test_adata.obs['batch'].tolist()
            data_batch_ids = np.array(batch_vocab(data_batch_ids))
            
            cell_pred = np.concatenate(cell_pred)
            cell_acc = (cell_pred == cell_labels).sum() / len(cell_labels)
            logger.info(f'Cell type accuracy: {cell_acc:.4f}')
            
            
            # data_cell_ids must be same as cell_labels, ensure the order
            assert np.all(data_cell_ids == cell_labels)
            assert np.all(data_batch_ids == batch_labels)
            # now cell_labels have shape (len(data_cell_ids), 1), convert to (len(data_cell_ids),)


            test_adata.obsm['embedding'] = embeddings
            test_adata.obs['celltype'] = test_adata.obs['annot'].astype('category')
            test_adata.obs['str_batch'] = test_adata.obs['batch'].astype(str)
            eval_scib_metrics(test_adata, logger, batch_key='str_batch', label_key='celltype')
            
            del non_dist_model, val_non_dist_loader, test_adata
            gc.collect()
    
    dist.barrier()
    # if eval cell type acc, first finetune the model with cell type
    if train_config['cell_type_epochs'] > 0:
        if is_master:
            logger.info("Init Training set, train model with cell type")
        train_set = PairedSCDataset(
            rna_file = data_config['train']['rna_path'],
            atac_file= data_config['train']['atac_path'],
            rna_key = data_config['train']['rna_key'],
            atac_key = data_config['train']['atac_key'],
            rna_vocab = rna_vocab,
            atac_vocab = atac_vocab,
            cell_vocab = cell_vocab,
            batch_vocab= batch_vocab,
            chr_vocab = chr_vocab,
            gene2chr_file= vocab_config['gene2chr_path'],
            rna_max_len = train_config['model']['rna_max_len'],
            atac_max_len = train_config['model']['atac_max_len'],
            pad_token = pad['token'],
            rna_pad_value = pad['value'],
            cls_token = cls['token'],
            logger = logger,
        )
                                
        gc.collect()
        train_sampler = DistributedSampler(train_set)
        train_loader = DataLoader(train_set, batch_size=BATCH_SIZE, sampler=train_sampler, prefetch_factor=4, num_workers=4)
        
        for i in range(1, train_config['cell_type_epochs'] + 1):
            train_loader.sampler.set_epoch(i)
            if is_master:
                logger.info('Training with {} samples, steps: {}'.format(len(train_loader.dataset), len(train_loader)))
            model.train()
            dist.barrier()
            running_loss = {'mvc': 0.0, 'cell': 0.0, 'total': 0.0}
            cum_acc_cell = 0.0
            cum_acc_value = 0.0
            for index, batch in enumerate(train_loader):
                index += 1
                steps += 1
                
                rna_values = batch['rna_values'].to(device)
                rna_ids = batch['rna_ids'].to(device)
                atac_ids = batch['atac_ids'].to(device)
                cell_ids = batch['cell_ids'].to(device)
                batch_ids = batch['batch_ids'].to(device)
                rna_chrs = batch['rna_chrs'].to(device)
                atac_chrs = batch['atac_chrs'].to(device)
                
                padding_positions = atac_ids.eq(atac_vocab[pad['token']])
            
                if index % GRADIENT_ACCUMULATION != 0 and index != len(train_loader):
                    with model.no_sync():
                        with torch.cuda.amp.autocast(enabled=train_config['amp'], dtype= torch.bfloat16):
                            # finetue using all expression values, do not mask
                            output = model(atac = atac_ids, rna = rna_ids, src_key_padding_mask = padding_positions, batch_id = batch_ids, rna_chrs = rna_chrs, atac_chrs = atac_chrs)

                            mvc_loss = atac_cross_entropy_loss(output['mvc_pred'].transpose(1, 2), rna_values)
                            cell_loss = cross_entropy_loss(output['cell_pred'], cell_ids)
                            # only train the cell loss
                            loss =  cell_loss + mvc_loss * 0.0
                            
                            running_loss['mvc'] += mvc_loss.item()
                            running_loss['cell'] += cell_loss.item()
                            running_loss['total'] += loss.item()
                            
                            loss = loss / GRADIENT_ACCUMULATION
                        scaler.scale(loss).backward()
                else:
                    with torch.cuda.amp.autocast(enabled=train_config['amp'], dtype= torch.bfloat16):
                        output = model(atac = atac_ids, rna = rna_ids, src_key_padding_mask = padding_positions, batch_id = batch_ids, rna_chrs = rna_chrs, atac_chrs = atac_chrs)
                        
                        mvc_loss = atac_cross_entropy_loss(output['mvc_pred'].transpose(1, 2), rna_values)
                        
                        cell_loss = cross_entropy_loss(output['cell_pred'], cell_ids)
                        loss =  cell_loss + mvc_loss * 0.0
                        
                        running_loss['mvc'] += mvc_loss.item()
                        running_loss['cell'] += cell_loss.item()
                        running_loss['total'] += loss.item()
                        if is_master:
                            tb_logger.add_scalar('train/mvc_loss', mvc_loss.item(), steps)
                            tb_logger.add_scalar('train/cell_loss', cell_loss.item(), steps)
                            tb_logger.add_scalar('train/total_loss', loss.item(), steps)
                            logger.info(f'Epoch: {i} | Step: {index} | MVC Loss: {mvc_loss:.4f} | Cell Type Loss: {cell_loss:.4f} | Total Loss: {loss:.4f}')
                        loss = loss / GRADIENT_ACCUMULATION
                    scaler.scale(loss).backward()
                    scaler.unscale_(optimizer)
                    torch.nn.utils.clip_grad_norm_(model.parameters(), int(1e2))
                    scaler.step(optimizer)
                    scaler.update()
                    optimizer.zero_grad()
                # cell type accuracy
                type_pred = softmax(output['cell_pred'])
                type_pred = type_pred.argmax(dim=-1)
                cum_acc_cell += (type_pred.eq(cell_ids)).sum().item() / len(cell_ids)
                
                value_pred = softmax(output['mvc_pred']).argmax(dim=-1)
                # expression value accuracy
                non_pad_idx = rna_values.ne(pad['value'])
                non_pad_pred = value_pred[non_pad_idx]
                non_pad_label = rna_values[non_pad_idx]
                cum_acc_value += (non_pad_pred.eq(non_pad_label).sum().item()) / non_pad_label.size(0)
                
            cum_acc_cell = 100 * cum_acc_cell / index
            cum_acc_cell = get_reduced(cum_acc_cell, local_rank, 0, world_size)
            
            cum_acc_value = 100 * cum_acc_value / index
            cum_acc_value = get_reduced(cum_acc_value, local_rank, 0, world_size)
            for key in running_loss:
                running_loss[key] = running_loss[key] / index
                running_loss[key] = get_reduced(running_loss[key], local_rank, 0, world_size)
            if is_master:
                    logger.info(f'Epoch: {i} | MVC Loss: {running_loss["mvc"]:.4f} | Cell Type Loss: {running_loss["cell"]:.4f} | Total Loss: {running_loss["total"]:.4f} | Cell Type Accuracy: {cum_acc_cell:.2f} | Expression Value Accuracy: {cum_acc_value:.2f}')
            dist.barrier()
            scheduler.step()
            
            # del train_set, train_sampler, train_loader
            if is_master and i % save_ckpt_freq == 0:
                logger.info("Saving the finetuned model")
                save_ckpt(i, steps,  model, optimizer, scheduler, scaler, running_loss["total"], task_name, ckpt_dir)
            
            if i % valid_config['freq'] == 0:
                model.eval()
                dist.barrier()
                running_loss = {'mvc': 0.0, 'cell': 0.0, 'total': 0.0}
                
                cum_acc_cell = 0.0
                cum_acc_value = 0.0
                if is_master:
                    logger.info('Start validation')
                with torch.no_grad():
                    for index, batch in enumerate(val_loader):
                        index += 1
                        
                        rna_values = batch['rna_values'].to(device)
                        rna_ids = batch['rna_ids'].to(device)
                        atac_ids = batch['atac_ids'].to(device)
                        cell_ids = batch['cell_ids'].to(device)
                        batch_ids = batch['batch_ids'].to(device)
                        rna_chrs = batch['rna_chrs'].to(device)
                        atac_chrs = batch['atac_chrs'].to(device)

                        padding_positions = atac_ids.eq(atac_vocab[pad['token']])
                        with torch.cuda.amp.autocast(enabled=train_config['amp'], dtype= torch.bfloat16):
                            output = model(atac = atac_ids, rna = rna_ids, src_key_padding_mask = padding_positions, batch_id = batch_ids, rna_chrs = rna_chrs, atac_chrs = atac_chrs)
                            
                            mvc_loss = atac_cross_entropy_loss(output['mvc_pred'].transpose(1, 2), rna_values)
                            cell_loss = cross_entropy_loss(output['cell_pred'], cell_ids)
                            loss =  cell_loss + mvc_loss * 0.0
                            
                        running_loss['mvc'] += mvc_loss.item()
                        running_loss['cell'] += cell_loss.item()
                        running_loss['total'] += loss.item()
                        
                        type_pred = softmax(output['cell_pred'])
                        type_pred = type_pred.argmax(dim=-1)
                        cum_acc_cell += (type_pred.eq(cell_ids)).sum().item() / len(cell_ids)
                        
                        value_pred = softmax(output['mvc_pred']).argmax(dim=-1)
                        # expression value accuracy
                        non_pad_idx = rna_values.ne(pad['value'])
                        non_pad_pred = value_pred[non_pad_idx]
                        non_pad_label = rna_values[non_pad_idx]
                        cum_acc_value += (non_pad_pred.eq(non_pad_label).sum().item()) / non_pad_label.size(0)
                        # break   
                for key in running_loss:
                    running_loss[key] = running_loss[key] / index
                    running_loss[key] = get_reduced(running_loss[key], local_rank, 0, world_size)
                cum_acc_cell = 100 * cum_acc_cell / index
                cum_acc_cell = get_reduced(cum_acc_cell, local_rank, 0, world_size)
                
                cum_acc_value = 100 * cum_acc_value / index
                cum_acc_value = get_reduced(cum_acc_value, local_rank, 0, world_size)
                # del val_set, val_sampler, val_loader
                if is_master:
                    logger.info(f'MVC Loss: {running_loss["mvc"]:.4f} | Cell Type Loss: {running_loss["cell"]:.4f} | Total Loss: {running_loss["total"]:.4f} | Cell Type Accuracy: {cum_acc_cell:.2f} | Expression Value Accuracy: {cum_acc_value:.2f}')

if __name__ == '__main__':
    main()
    
```
