## Supplementary material for "EpiFoundation: A Foundation Model for Single-Cell ATAC-seq via Peak-to-Gene Alignment": Code for EpiFoundation: loss.py

```
import torch
from   torch import nn
import torch.nn.functional as F


class MaskedMSELoss(nn.Module):
    def __init__(self):
        super().__init__()
        
    def masked_mse_loss(
        self, input: torch.Tensor, target: torch.Tensor, mask: torch.Tensor
    ) -> torch.Tensor:
        """
        Compute the masked MSE loss between input and target.
        """
        mask = mask.float()
        loss = F.mse_loss(input * mask, target * mask, reduction="sum")
        return loss / mask.sum()

    def forward(self, input: torch.Tensor, target: torch.Tensor, mask: torch.Tensor) -> torch.Tensor:
        return self.masked_mse_loss(input, target, mask)
```
