## Supplementary material for "EpiFoundation: A Foundation Model for Single-Cell ATAC-seq via Peak-to-Gene Alignment": Code for EpiFoundation: __init__.py

```
from EpiFoundation.model.EpiFoundation import EpiFoundation
import logging
import sys

logger = logging.getLogger("scMultiomics")
# check if logger has been initialized
if not logger.hasHandlers() or len(logger.handlers) == 0:
    logger.propagate = False
    logger.setLevel(logging.INFO)
    handler = logging.StreamHandler(sys.stdout)
    handler.setLevel(logging.INFO)
    formatter = logging.Formatter(
        "%(name)s - %(levelname)s - %(message)s", datefmt="%H:%M:%S"
    )
    handler.setFormatter(formatter)
    logger.addHandler(handler)
```
