## Supplementary material for "EpiFoundation: A Foundation Model for Single-Cell ATAC-seq via Peak-to-Gene Alignment": Code for EpiFoundation: EpiFoundation.py

```
import math
import numpy as np
import torch
import torch.nn.functional as F
from torch import nn, Tensor
from torch.cuda.amp import autocast
from einops import rearrange, repeat

from functools import partial
from contextlib import contextmanager

from local_attention import LocalAttention
from typing import Dict, Mapping, Optional, Tuple, Any, Union
from model.performer import Performer, cast_tuple
from model.transformer import TransformerModel


class ClsDecoder(nn.Module):
    """
    Decoder for classification task.
    """
    def __init__(
        self,
        d_model: int,
        n_cls: int,
        nlayers: int = 3,
        activation: callable = nn.ReLU,
    ):
        super().__init__()
        # module list
        self._decoder = nn.ModuleList()
        for i in range(nlayers - 1):
            self._decoder.append(nn.Linear(d_model, d_model))
            self._decoder.append(activation())
            self._decoder.append(nn.LayerNorm(d_model))
        self.out_layer = nn.Linear(d_model, n_cls)

    def forward(self, x: Tensor) -> Tensor:
        """
        Args:
            x: Tensor, shape [batch_size, embsize]
        """
        for layer in self._decoder:
            x = layer(x)
        return self.out_layer(x)

class GeneEncoder(nn.Module):
    def __init__(
        self,
        num_embeddings: int,
        embedding_dim: int,
        padding_idx: Optional[int] = None,
    ):
        super().__init__()
        self.embedding = nn.Embedding(
            num_embeddings, embedding_dim, padding_idx=padding_idx
        )
        self.enc_norm = nn.LayerNorm(embedding_dim)

    def forward(self, x: Tensor) -> Tensor:
        x = self.embedding(x)  # (batch, seq_len, embsize)
        x = self.enc_norm(x)
        return x

# positional embeddings
class AbsolutePositionalEmbedding(nn.Module):
    def __init__(self, max_seq_len, embed_dim):
        super().__init__()
        self.emb = nn.Embedding(max_seq_len, embed_dim)

    def forward(self, x):
        # x is of shape (batch, seq_len)
        # torch.arange is for generating a range of values from 0 to max_seq_len
        t = torch.arange(x.shape[1], device=x.device)
        return self.emb(t)
    
    
class Always(nn.Module):
    def __init__(self, val):
        super().__init__()
        self.val = val

    def forward(self, *args, **kwargs):
        return self.val
    
    
class ExprDecoder(nn.Module):
    def __init__(
        self,
        d_model: int,
        explicit_zero_prob: bool = False,
        use_batch_labels: bool = False,
        # catagory_num: Optional[int] = None,
    ):
        super().__init__()
        d_in = d_model * 2 if use_batch_labels else d_model
        self.fc = nn.Sequential(
            nn.Linear(d_in, d_model),
            nn.LeakyReLU(),
            nn.Linear(d_model, d_model),
            nn.LeakyReLU(),
            nn.Linear(d_model, 1),
        )
        self.explicit_zero_prob = explicit_zero_prob
        if explicit_zero_prob:
            self.zero_logit = nn.Sequential(
                nn.Linear(d_in, d_model),
                nn.LeakyReLU(),
                nn.Linear(d_model, d_model),
                nn.LeakyReLU(),
                nn.Linear(d_model, 1),
            )

    def forward(self, x: Tensor) -> Dict[str, Tensor]:
        """x is the output of the transformer, (batch, seq_len, d_model)"""
        pred_value = self.fc(x).squeeze(-1)  # (batch, seq_len)

        # if not self.explicit_zero_prob:
        #     return dict(pred=pred_value)
        # zero_logits = self.zero_logit(x).squeeze(-1)  # (batch, seq_len)
        # zero_probs = torch.sigmoid(zero_logits)
        return pred_value
        # TODO: note that the return currently is only for training. Since decoder
        # is not used in the test setting for the integration task, the eval/inference
        # logic is not implemented yet. However, remember to implement it when
        # the decoder is used in any test setting. The inference logic will need
        # to sample from the bernoulli distribution with the zero_probs.
        
        
class PretrainDecoder(nn.Module):
    """
    Decoder for the masked value prediction for cell embeddings.
    """

    def __init__(
        self,
        d_model: int,
        arch_style: str = "inner product",
        query_activation: nn.Module = nn.Sigmoid,
        hidden_activation: nn.Module = nn.PReLU,
        explicit_zero_prob: bool = False,
        use_batch_labels: bool = False,
        catagory_num: Optional[int] = 2,
    ) -> None:
        """
        Args:
            d_model (:obj:`int`): dimension of the gene embedding.
            arch_style (:obj:`str`): architecture style of the decoder, choice from
                1. "inner product" or 2. "concat query" or 3. "sum query".
            query_activation (:obj:`nn.Module`): activation function for the query
                vectors.
            hidden_activation (:obj:`nn.Module`): activation function for the hidden
                layers.
        """
        super().__init__()
        d_in = d_model * 2 if use_batch_labels else d_model
        if arch_style in ["inner product", "inner product, detach"]:
            self.gene2query = nn.Linear(d_model, d_model)
            self.query_activation = query_activation()
            self.W = nn.Linear(d_model, d_in, bias=False)
            if explicit_zero_prob:  # by default, gene-wise prob rate
                self.W_zero_logit = nn.Linear(d_model, d_in)
        elif arch_style == "concat query":
            self.gene2query = nn.Linear(d_model, 128)
            self.query_activation = query_activation()
            self.fc1 = nn.Linear(d_in + 128, 128)
            self.hidden_activation = hidden_activation()
            # self.fc2 = nn.Linear(64, 1)
            # for rna value prediction
            self.fc2 = nn.Linear(128, catagory_num)
        elif arch_style == "sum query":
            self.gene2query = nn.Linear(d_model, d_model)
            self.query_activation = query_activation()
            self.fc1 = nn.Linear(d_in, 128)
            self.hidden_activation = hidden_activation()
            self.fc2 = nn.Linear(128, catagory_num)
        else:
            raise ValueError(f"Unknown arch_style: {arch_style}")

        self.arch_style = arch_style
        self.do_detach = arch_style.endswith("detach")
        self.explicit_zero_prob = explicit_zero_prob

    def forward(
        self, cell_emb: Tensor, gene_embs: Tensor
    ) -> Union[Tensor, Dict[str, Tensor]]:
        """
        Args:
            cell_emb: Tensor, shape (batch, embsize=d_model)
            gene_embs: Tensor, shape (batch, seq_len, embsize=d_model)
        """
        gene_embs = gene_embs.detach() if self.do_detach else gene_embs
        if self.arch_style in ["inner product", "inner product, detach"]:
            query_vecs = self.query_activation(self.gene2query(gene_embs))
            cell_emb = cell_emb.unsqueeze(2)  # (batch, embsize, 1)
            # the pred gene expr values, # (batch, seq_len)
            pred_value = torch.bmm(self.W(query_vecs), cell_emb).squeeze(2)
            if not self.explicit_zero_prob:
                return pred_value
            # zero logits need to based on the cell_emb, because of input exprs
            zero_logits = torch.bmm(self.W_zero_logit(query_vecs), cell_emb).squeeze(2)
            zero_probs = torch.sigmoid(zero_logits)
            return pred_value
        elif self.arch_style == "concat query":
            query_vecs = self.query_activation(self.gene2query(gene_embs))
            # expand cell_emb to (batch, seq_len, embsize)
            cell_emb = cell_emb.unsqueeze(1).expand(-1, gene_embs.shape[1], -1)

            h = self.hidden_activation(
                self.fc1(torch.cat([cell_emb, query_vecs], dim=2))
            )
            if self.explicit_zero_prob:
                raise NotImplementedError
            return self.fc2(h).squeeze(2)  # (batch, seq_len)
        elif self.arch_style == "sum query":
            query_vecs = self.query_activation(self.gene2query(gene_embs))
            cell_emb = cell_emb.unsqueeze(1)

            h = self.hidden_activation(self.fc1(cell_emb + query_vecs))
            if self.explicit_zero_prob:
                raise NotImplementedError
            return self.fc2(h).squeeze(2)  # (batch, seq_len)
    
class CategoryValueEncoder(nn.Module):
    def __init__(
        self,
        num_embeddings: int,
        embedding_dim: int,
        padding_idx: Optional[int] = None,
    ):
        super().__init__()
        self.embedding = nn.Embedding(
            num_embeddings, embedding_dim, padding_idx=padding_idx
        )
        self.enc_norm = nn.LayerNorm(embedding_dim)

    def forward(self, x: Tensor) -> Tensor:
        x = x.long()
        x = self.embedding(x)  # (batch, seq_len, embsize)
        x = self.enc_norm(x)
        return x

# sinusoidal positional embeddings

class Gene2VecPositionalEmbedding(nn.Module):
    def __init__(self, max_seq_len, embed_dim):
        super().__init__()
        gene2vec_weight = np.load('../data/gene2vec_16906.npy')
        gene2vec_weight = np.concatenate((gene2vec_weight, np.zeros((1, gene2vec_weight.shape[1]))), axis=0)
        gene2vec_weight = torch.from_numpy(gene2vec_weight)
        self.emb = nn.Embedding.from_pretrained(gene2vec_weight)

    def forward(self, x):
        t = torch.arange(x.shape[1], device=x.device)
        return self.emb(t)

class BatchLabelEncoder(nn.Module):
    def __init__(
        self,
        num_embeddings: int,
        embedding_dim: int,
        padding_idx: Optional[int] = None,
    ):
        super().__init__()
        self.embedding = nn.Embedding(
            num_embeddings, embedding_dim, padding_idx=padding_idx
        )
        self.enc_norm = nn.LayerNorm(embedding_dim)

    def forward(self, x: Tensor) -> Tensor:
        x = self.embedding(x)  # (batch, embsize)
        x = self.enc_norm(x)
        return x

class EpiFoundation(nn.Module):
    def __init__(
        self,
        num_class_cell,                     # num of cell categories    
        num_rnas,                           # num of genes (or atac peaks)
        num_atacs,                          # num of genes (or atac peaks)
        num_values,                         # num of values
        num_chrs,                           # num of chromosomes
        embed_dim,                          # embed_dim of tokens
        depth,                              # layers
        heads,                              # num of heads
        head_dim = 64,                      # embed_dim of heads
        encoder:str = 'transformer',        # encoder type, performer or transformer
        dropout = 0.2,
        pad_token_idx_atac = 0,             # padding token index , shoule be vocab[pad_token], set to 0 for debugging
        pad_token_idx_rna = 0,              # padding token index , shoule be vocab[pad_token], set to 0 for debugging
        cell_emb_style = "cls",             # cell embedding style
        mvc_arch_style = "inner product",   # mvc decoder architecture style 
        use_batch_labels = False,           # whether to use batch labels
        batch_label_num = 13,                # num of batch labels
        use_chr_labels = False,             # whether to use chr labels
        transformer_backend = 'flash',    # backend of transformer, pytorch or einsum
        stage = 'pretrain',                 # stage of the model, pretrain or finetune
    ):
        super().__init__()

        self.stage = stage
        # self.express_emb = nn.Embedding(num_tokens, embed_dim)
        self.encoder_type = encoder
        self.cell_emb_style = cell_emb_style
        self.embed_dim = embed_dim

        # determine positional embedding
        self.rna_emb = GeneEncoder(num_rnas, embed_dim, padding_idx=pad_token_idx_rna)
        self.atac_emb = GeneEncoder(num_atacs, embed_dim, padding_idx=pad_token_idx_atac)
        
        if use_batch_labels:
            self.batch_emb = BatchLabelEncoder(batch_label_num, embed_dim)
        else:
            self.batch_emb = None
            
        if use_chr_labels:
            self.chr_emb = GeneEncoder(num_chrs, embed_dim)
        else:  
            self.chr_emb = None

        self.dropout_rna = nn.Dropout(dropout)
        self.dropout_atac = nn.Dropout(dropout)

        if encoder == 'performer':
            self.encoder = Performer(embed_dim, depth, heads, head_dim)
        elif encoder == 'transformer':
            self.encoder = TransformerModel(d_model=embed_dim, nhead=heads, nlayers=depth, d_hid= head_dim, dropout=dropout, fast_transformer_backend=transformer_backend)
        # self.encoder = Performer(embed_dim, depth, heads, head_dim, local_attn_heads, local_window_size, causal, ff_mult, nb_features, feature_redraw_interval, reversible, ff_chunks, generalized_attention, kernel_fn, use_scalenorm, use_rezero, ff_glu, ff_dropout, attn_dropout, cross_attend, no_projection, auto_check_redraw, qkv_bias)
        self.norm = nn.LayerNorm(embed_dim)
        # self.to_out = nn.Linear(embed_dim, 1) 
        self.cls_decoder = ClsDecoder(embed_dim, num_class_cell)
        self.mvc_decoder = PretrainDecoder(embed_dim, arch_style = mvc_arch_style, use_batch_labels=use_batch_labels, catagory_num = num_values)
        self.bn_atac = nn.BatchNorm1d(embed_dim, eps=6.1e-5)
        self.bn_rna = nn.BatchNorm1d(embed_dim, eps=6.1e-5)
        if stage == 'value_finetune':
            for param in self.cls_decoder.parameters():
                param.requires_grad = False
            for param in self.mvc_decoder.parameters():
                param.requires_grad = False
            self.value_decoder = PretrainDecoder(embed_dim, arch_style = mvc_arch_style, use_batch_labels=use_batch_labels, catagory_num = 1)
    
    def _get_cell_emb_from_layer(
        self, layer_output: Tensor, weights: Tensor = None
    ) -> Tensor:
        """
        Args:
            layer_output(:obj:`Tensor`): shape (batch, seq_len, embsize)
            weights(:obj:`Tensor`): shape (batch, seq_len), optional and only used
                when :attr:`self.cell_emb_style` is "w-pool".

        Returns:
            :obj:`Tensor`: shape (batch, embsize)
        """
        if self.cell_emb_style == "cls":
            cell_emb = layer_output[:, 0, :]  # (batch, embsize)
        elif self.cell_emb_style == "avg-pool":
            cell_emb = torch.mean(layer_output, dim=1)
        elif self.cell_emb_style == "w-pool":
            if weights is None:
                raise ValueError("weights is required when cell_emb_style is w-pool")
            if weights.dim() != 2:
                raise ValueError("weights should be 2D")
            cell_emb = torch.sum(layer_output * weights.unsqueeze(2), dim=1)
            cell_emb = F.normalize(cell_emb, p=2, dim=1)  # (batch, embsize)
        return cell_emb

    def forward(self, atac, rna, src_key_padding_mask: Optional[Tensor] = None, **kwargs):
        atac_emb = self.atac_emb(atac)
        if self.chr_emb is not None:
            chr_emb = self.chr_emb(kwargs['atac_chrs'])
            atac_emb = atac_emb + chr_emb
        atac_emb = self.dropout_atac(atac_emb)
        atac_emb = self.bn_atac(atac_emb.permute(0, 2, 1)).permute(0, 2, 1)
        
        x = self.encoder(atac_emb, src_key_padding_mask = src_key_padding_mask)
        transformer_output = self.norm(x) # (batch, seq_len, embsize)
        
        rna_emb = self.rna_emb(rna)
        if self.chr_emb is not None:
            chr_emb = self.chr_emb(kwargs['rna_chrs'])
            rna_emb = rna_emb + chr_emb
        rna_emb = self.dropout_rna(rna_emb)
        rna_emb = self.bn_rna(rna_emb.permute(0, 2, 1)).permute(0, 2, 1)
        
        output = {}
        cell_emb = self._get_cell_emb_from_layer(transformer_output)
        
        if self.batch_emb is not None:
            batch_emb = self.batch_emb(kwargs['batch_id'])
            cell_emb_w_batch = torch.cat((cell_emb, batch_emb), dim = 1)
            output["mvc_pred"] = self.mvc_decoder(cell_emb_w_batch, rna_emb)
            if self.stage == 'value_finetune':
                output["value_pred"] = self.value_decoder(cell_emb_w_batch, rna_emb)
        else:
            output["mvc_pred"] = self.mvc_decoder(cell_emb, rna_emb)
            if self.stage == 'value_finetune':
                output["value_pred"] = self.value_decoder(cell_emb, rna_emb)
        
        output["cell_emb"] = cell_emb
        output["cell_pred"] = self.cls_decoder(cell_emb) # (batch, n_cls)
        
        return output
```
