## Supplementary material for "EpiFoundation: A Foundation Model for Single-Cell ATAC-seq via Peak-to-Gene Alignment": Code for EpiFoundation: flashDiff.py

```
import math
import torch
import torch.nn.functional as F
from torch import nn

# from .kernel.rotary import apply_rotary_emb
from flash_attn import flash_attn_func

def init_method(tensor, **kwargs):
    nn.init.kaiming_uniform_(tensor, a=math.sqrt(5))

def repeat_kv(x: torch.Tensor, n_rep: int) -> torch.Tensor:
    """torch.repeat_interleave(x, dim=1, repeats=n_rep)"""
    bs, n_kv_heads, slen, head_dim = x.shape
    if n_rep == 1:
        return x
    return (
        x[:, :, None, :, :]
        .expand(bs, n_kv_heads, n_rep, slen, head_dim)
        .reshape(bs, n_kv_heads * n_rep, slen, head_dim)
    )

def lambda_init_fn(depth):
    return 0.8 - 0.6 * math.exp(-0.3 * depth)

class RMSNorm(nn.Module):
    def __init__(self, dim: int, eps: float = 1e-6, elementwise_affine=True, memory_efficient=False):
        super().__init__()
        self.dim = dim
        self.eps = eps
        self.elementwise_affine = elementwise_affine
        if self.elementwise_affine:
            self.weight = nn.Parameter(torch.ones(dim))
        else:
            self.register_parameter('weight', None)

    def _norm(self, x):
        return x * torch.rsqrt(x.pow(2).mean(-1, keepdim=True) + self.eps)

    def forward(self, x):
        output = self._norm(x.float()).type_as(x)
        if self.weight is not None:
            output = output * self.weight
        return output

    def extra_repr(self) -> str:
        return f'dim={self.dim}, eps={self.eps}, elementwise_affine={self.elementwise_affine}'


class MultiheadFlashDiff(nn.Module):
    """
    (Recommended)
    DiffAttn implemented with FlashAttention, for packages that support different qk/v dimensions
    e.g., our customized-flash-attention (https://aka.ms/flash-diff) and xformers (https://github.com/facebookresearch/xformers)
    """
    def __init__(
        self,
        embed_dim,
        num_heads,
        attention_dropout=0.0,
        lambda_init=0.8,
    ):
        super().__init__()
        self.embed_dim = embed_dim
        # num_heads set to half of Transformer's #heads
        # self.num_heads = num_heads // args.model_parallel_size
        self.num_heads = num_heads
        self.num_kv_heads = num_heads
        # self.num_kv_heads = args.decoder_kv_attention_heads // args.model_parallel_size if args.decoder_kv_attention_heads is not None else num_heads // args.model_parallel_size
        self.n_rep = self.num_heads // self.num_kv_heads
        self.dropout = attention_dropout
        
        self.head_dim = embed_dim // num_heads // 2
        self.scaling = self.head_dim ** -0.5
        
        self.q_proj = nn.Linear(embed_dim, embed_dim, bias=False)
        self.k_proj = nn.Linear(embed_dim, embed_dim // self.n_rep, bias=False)
        self.v_proj = nn.Linear(embed_dim, embed_dim // self.n_rep, bias=False)
        self.out_proj = nn.Linear(embed_dim, embed_dim, bias=False)

        self.lambda_init = lambda_init
        self.lambda_q1 = nn.Parameter(torch.zeros(self.head_dim, dtype=torch.float32).normal_(mean=0,std=0.1))
        self.lambda_k1 = nn.Parameter(torch.zeros(self.head_dim, dtype=torch.float32).normal_(mean=0,std=0.1))
        self.lambda_q2 = nn.Parameter(torch.zeros(self.head_dim, dtype=torch.float32).normal_(mean=0,std=0.1))
        self.lambda_k2 = nn.Parameter(torch.zeros(self.head_dim, dtype=torch.float32).normal_(mean=0,std=0.1))

        self.subln = RMSNorm(2 * self.head_dim, eps=1e-5, elementwise_affine=True)
        
    def forward(
        self,
        x,
    ):
        bsz, tgt_len, embed_dim = x.size()
        src_len = tgt_len

        q = self.q_proj(x)
        k = self.k_proj(x)
        v = self.v_proj(x)

        q = q.view(bsz, tgt_len, 2 * self.num_heads, self.head_dim)
        k = k.view(bsz, src_len, 2 * self.num_kv_heads, self.head_dim)
        v = v.view(bsz, src_len, self.num_kv_heads, 2, self.head_dim)


        offset = src_len - tgt_len
        q = q.reshape(bsz, tgt_len, self.num_heads, 2, self.head_dim)
        k = k.reshape(bsz, src_len, self.num_kv_heads, 2, self.head_dim)
        q1, q2 = q[:, :, :, 0], q[:, :, :, 1]
        k1, k2 = k[:, :, :, 0], k[:, :, :, 1]
        v1, v2 = v[:, :, :, 0], v[:, :, :, 1]

        attn11 = flash_attn_func(q1, k1, v1, dropout_p=self.dropout)
        attn12 = flash_attn_func(q1, k1, v2, dropout_p=self.dropout)
        attn1 = torch.cat([attn11, attn12], dim=-1)
        
        attn21 = flash_attn_func(q2, k2, v1, dropout_p=self.dropout)
        attn22 = flash_attn_func(q2, k2, v2, dropout_p=self.dropout)
        attn2 = torch.cat([attn21, attn22], dim=-1)
        
        lambda_1 = torch.exp(torch.sum(self.lambda_q1 * self.lambda_k1, dim=-1).float()).type_as(q)
        lambda_2 = torch.exp(torch.sum(self.lambda_q2 * self.lambda_k2, dim=-1).float()).type_as(q)
        lambda_full = lambda_1 - lambda_2 + self.lambda_init
        attn = attn1 - lambda_full * attn2

        attn = self.subln(attn)
        attn = attn * (1 - self.lambda_init)
        attn = attn.reshape(bsz, tgt_len, self.num_heads * 2 * self.head_dim)
        
        attn = self.out_proj(attn)
        return attn
    
```
