## Supplementary material for "EpiFoundation: A Foundation Model for Single-Cell ATAC-seq via Peak-to-Gene Alignment": Code for EpiFoundation: flashMHA.py

```
import math
import torch
import torch.nn as nn
import torch.nn.functional as F

from einops import rearrange

from flash_attn.flash_attn_interface import flash_attn_varlen_qkvpacked_func
from flash_attn.bert_padding import unpad_input, pad_input


class FlashAttention(nn.Module):
    """Implement the scaled dot product attention with softmax.
    Arguments
    ---------
        softmax_scale: The temperature to use for the softmax attention.
                    (default: 1/sqrt(d_keys) where d_keys is computed at
                    runtime)
        attention_dropout: The dropout rate to apply to the attention
                        (default: 0.0)
    """
    def __init__(self, softmax_scale=None, attention_dropout=0.0):
        super().__init__()
        self.softmax_scale = softmax_scale
        self.dropout_p = attention_dropout
        
    def get_attention_weights(self, qkv):
        batch_size, seqlen, _, nheads, head_dim = qkv.shape
        q = qkv[:, :, 0, :, :]
        k = qkv[:, :, 1, :, :]
        
        q = rearrange(q, 'b s h d -> b h s d')
        k = rearrange(k, 'b s h d -> b h s d')
        
        scale = head_dim ** -0.5
        attn_weights = torch.matmul(q, k.transpose(-2, -1)) * scale
        attn_weights = F.softmax(attn_weights, dim=-1)
        return attn_weights

    def forward(self, qkv, key_padding_mask=None, causal=False, cu_seqlens=None,
                max_s=None, need_weights=False):
        """Implements the multihead softmax attention.
        Arguments
        ---------
            qkv: The tensor containing the query, key, and value. (B, S, 3, H, D) if key_padding_mask is None
                if unpadded: (nnz, 3, h, d)
            key_padding_mask: a bool tensor of shape (B, S)
        """
        # assert not need_weights
        assert qkv.dtype in [torch.float16, torch.bfloat16]
        assert qkv.is_cuda
        if need_weights:
            attn_weights = self.get_attention_weights(qkv)
            # get the attention weights for the first token
            attn_weights = attn_weights[:, :, 0, :]
        else:
            attn_weights = None

        if cu_seqlens is None:
            batch_size = qkv.shape[0]
            seqlen = qkv.shape[1]
            if key_padding_mask is None:
                qkv = rearrange(qkv, 'b s ... -> (b s) ...')
                max_s = seqlen
                cu_seqlens = torch.arange(0, (batch_size + 1) * seqlen, step=seqlen, dtype=torch.int32,
                                        device=qkv.device)
                output = flash_attn_varlen_qkvpacked_func(
                    qkv, cu_seqlens, max_s, self.dropout_p if self.training else 0.0,
                    softmax_scale=self.softmax_scale, causal=causal
                )
                output = rearrange(output, '(b s) ... -> b s ...', b=batch_size)
            else:
                nheads = qkv.shape[-2]
                x = rearrange(qkv, 'b s three h d -> b s (three h d)')
                x_unpad, indices, cu_seqlens, max_s = unpad_input(x, key_padding_mask)
                x_unpad = rearrange(x_unpad, 'nnz (three h d) -> nnz three h d', three=3, h=nheads)
                output_unpad = flash_attn_varlen_qkvpacked_func(
                    x_unpad, cu_seqlens, max_s, self.dropout_p if self.training else 0.0,
                    softmax_scale=self.softmax_scale, causal=causal
                )
                output = rearrange(pad_input(rearrange(output_unpad, 'nnz h d -> nnz (h d)'),
                                            indices, batch_size, seqlen),
                                'b s (h d) -> b s h d', h=nheads)
        else:
            assert max_s is not None
            output = flash_attn_varlen_qkvpacked_func(
                qkv, cu_seqlens, max_s, self.dropout_p if self.training else 0.0,
                softmax_scale=self.softmax_scale, causal=causal
            )
        return output, attn_weights


class FlashMHA(nn.Module):

    def __init__(self, embed_dim, num_heads, bias=True, batch_first=True, attention_dropout=0.0,
                causal=False, device=None, dtype=None) -> None:
        assert batch_first
        factory_kwargs = {'device': device, 'dtype': dtype}
        super().__init__()
        self.embed_dim = embed_dim
        self.causal = causal

        self.num_heads = num_heads
        assert self.embed_dim % num_heads == 0, "self.kdim must be divisible by num_heads"
        self.head_dim = self.embed_dim // num_heads
        assert self.head_dim % 8 == 0 and self.head_dim <= 128, "Only support head_dim <= 128 and divisible by 8"

        self.Wqkv = nn.Linear(embed_dim, 3 * embed_dim, bias=bias, **factory_kwargs)
        self.inner_attn = FlashAttention(attention_dropout=attention_dropout)
        self.out_proj = nn.Linear(embed_dim, embed_dim, bias=bias, **factory_kwargs)

    def forward(self, x, key_padding_mask=None, need_weights=False):
        """x: (batch, seqlen, hidden_dim) (where hidden_dim = num heads * head dim)
        key_padding_mask: bool tensor of shape (batch, seqlen)
        """
        qkv = self.Wqkv(x)
        qkv = rearrange(qkv, 'b s (three h d) -> b s three h d', three=3, h=self.num_heads)
        context, attn_weights = self.inner_attn(qkv, key_padding_mask=key_padding_mask,
                                                need_weights=need_weights, causal=self.causal)
        return self.out_proj(rearrange(context, 'b s h d -> b s (h d)')), attn_weights
```
