## Supplementary material for "EpiFoundation: A Foundation Model for Single-Cell ATAC-seq via Peak-to-Gene Alignment": Code for EpiFoundation: performer.py

```
import math
import numpy as np
import torch
import torch.nn.functional as F
from torch import nn, Tensor
from torch.cuda.amp import autocast
from einops import rearrange, repeat

from functools import partial
from contextlib import contextmanager

from local_attention import LocalAttention
from model.reversible import ReversibleSequence, SequentialSequence
from typing import Dict, Mapping, Optional, Tuple, Any, Union

try:
    from apex import amp
    APEX_AVAILABLE = True
except:
    APEX_AVAILABLE = False

# helpers

def exists(val):
    return val is not None

def empty(tensor):
    return tensor.numel() == 0

def default(val, d):
    return val if exists(val) else d

@contextmanager
def null_context():
    yield

def cast_tuple(val):
    return (val,) if not isinstance(val, tuple) else val

# def get_module_device(module):
#     return next(module.parameters).device

def get_module_device(module):
    try:
        return next(module.parameters()).device
    except StopIteration:
        # For nn.DataParallel compatibility in PyTorch 1.5
        def find_tensor_attributes(module):
            tuples = [(k, v) for k, v in module.__dict__.items() if torch.is_tensor(v)]
            return tuples
        gen = module._named_members(get_members_fn=find_tensor_attributes)
        first_tuple = next(gen)
        return first_tuple[1].device

def find_modules(nn_module, type):
    return [module for module in nn_module.modules() if isinstance(module, type)]


def rotate_every_two(x):
    x = rearrange(x, '... (d j) -> ... d j', j = 2)
    x1, x2 = x.unbind(dim = -1)
    x = torch.stack((-x2, x1), dim = -1)
    return rearrange(x, '... d j -> ... (d j)')

def apply_rotary_pos_emb(q, k, sinu_pos):
    sinu_pos = rearrange(sinu_pos, '() n (j d) -> n j d', j = 2)
    sin, cos = sinu_pos.unbind(dim = -2)
    sin, cos = map(lambda t: repeat(t, 'b n -> b (n j)', j = 2), (sin, cos))
    q, k = map(lambda t: (t * cos) + (rotate_every_two(t) * sin), (q, k))
    return q, k

# kernel functions

# transcribed from jax to pytorch from
# https://github.com/google-research/google-research/blob/master/performer/fast_attention/jax/fast_attention.py

def softmax_kernel(data, *, projection_matrix, is_query, normalize_data=True, eps=1e-4, device = None):
    b, h, *_ = data.shape

    data_normalizer = (data.shape[-1] ** -0.25) if normalize_data else 1.

    ratio = (projection_matrix.shape[0] ** -0.5)

    projection = repeat(projection_matrix, 'j d -> b h j d', b = b, h = h)
    projection = projection.type_as(data)

    data_dash = torch.einsum('...id,...jd->...ij', (data_normalizer * data), projection)

    diag_data = data ** 2
    diag_data = torch.sum(diag_data, dim=-1)
    diag_data = (diag_data / 2.0) * (data_normalizer ** 2)
    diag_data = diag_data.unsqueeze(dim=-1)

    if is_query:
        data_dash = ratio * (
            torch.exp(data_dash - diag_data -
                    torch.max(data_dash, dim=-1, keepdim=True).values) + eps)
    else:
        data_dash = ratio * (
            torch.exp(data_dash - diag_data - torch.max(data_dash)) + eps)

    return data_dash.type_as(data)

def generalized_kernel(data, *, projection_matrix, kernel_fn = nn.ReLU(), kernel_epsilon = 0.001, normalize_data = True, device = None):
    b, h, *_ = data.shape

    data_normalizer = (data.shape[-1] ** -0.25) if normalize_data else 1.

    if projection_matrix is None:
        return kernel_fn(data_normalizer * data) + kernel_epsilon

    projection = repeat(projection_matrix, 'j d -> b h j d', b = b, h = h)
    projection = projection.type_as(data)

    data_dash = torch.einsum('...id,...jd->...ij', (data_normalizer * data), projection)

    data_prime = kernel_fn(data_dash) + kernel_epsilon
    return data_prime.type_as(data)

def orthogonal_matrix_chunk(cols, device = None):
    unstructured_block = torch.randn((cols, cols), device = device)
    q, r = torch.qr(unstructured_block.cpu(), some = True)
    q, r = map(lambda t: t.to(device), (q, r))
    return q.t()

def gaussian_orthogonal_random_matrix(nb_rows, nb_columns, scaling = 0, device = None):
    nb_full_blocks = int(nb_rows / nb_columns)

    block_list = []

    for _ in range(nb_full_blocks):
        q = orthogonal_matrix_chunk(nb_columns, device = device)
        block_list.append(q)

    remaining_rows = nb_rows - nb_full_blocks * nb_columns
    if remaining_rows > 0:
        q = orthogonal_matrix_chunk(nb_columns, device = device)
        block_list.append(q[:remaining_rows])

    final_matrix = torch.cat(block_list)

    if scaling == 0:
        multiplier = torch.randn((nb_rows, nb_columns), device = device).norm(dim = 1)
    elif scaling == 1:
        multiplier = math.sqrt((float(nb_columns))) * torch.ones((nb_rows,), device = device)
    else:
        raise ValueError(f'Invalid scaling {scaling}')

    return torch.diag(multiplier) @ final_matrix

# linear attention classes with softmax kernel

# non-causal linear attention
def linear_attention(q, k, v):
    k_cumsum = k.sum(dim = -2)
    D_inv = 1. / torch.einsum('...nd,...d->...n', q, k_cumsum.type_as(q))
    context = torch.einsum('...nd,...ne->...de', k, v)
    out = torch.einsum('...de,...nd,...n->...ne', context, q, D_inv)
    return out

# efficient causal linear attention, created by EPFL
# TODO: rewrite EPFL's CUDA kernel to do mixed precision and remove half to float conversion and back
def causal_linear_attention(q, k, v, eps = 1e-6):
    from fast_transformers.causal_product import CausalDotProduct
    autocast_enabled = torch.is_autocast_enabled()
    is_half = isinstance(q, torch.cuda.HalfTensor)
    assert not is_half or APEX_AVAILABLE, 'half tensors can only be used if nvidia apex is available'
    cuda_context = null_context if not autocast_enabled else partial(autocast, enabled = False)

    causal_dot_product_fn = amp.float_function(CausalDotProduct.apply) if is_half else CausalDotProduct.apply

    k_cumsum = k.cumsum(dim=-2) + eps
    D_inv = 1. / torch.einsum('...nd,...nd->...n', q, k_cumsum.type_as(q))

    with cuda_context():
        if autocast_enabled:
            q, k, v = map(lambda t: t.float(), (q, k, v))

        out = causal_dot_product_fn(q, k, v)

    out = torch.einsum('...nd,...n->...nd', out, D_inv)
    return out

# inefficient causal linear attention, without cuda code, for reader's reference
# not being used
def causal_linear_attention_noncuda(q, k, v, chunk_size = 128):
    last_k_cumsum = 0
    last_context_cumsum = 0
    outs = []

    for q, k, v in zip(*map(lambda t: t.chunk(chunk_size, dim = -2), (q, k, v))):
        k_cumsum = last_k_cumsum + k.cumsum(dim=-2)

        D_inv = 1. / torch.einsum('...nd,...nd->...n', q, k_cumsum.type_as(q))
        context = torch.einsum('...nd,...ne->...nde', k, v)
        context_cumsum = last_context_cumsum + context.cumsum(dim=-3)
        out = torch.einsum('...nde,...nd,...n->...ne', context_cumsum, q, D_inv)

        last_k_cumsum = k_cumsum[:, :, -1:]
        last_context_cumsum = context_cumsum[:, :, -1:]
        outs.append(out)

    return torch.cat(outs, dim = -2)

def norm_tensor(tensor, dim=-1):
    return tensor / tensor.sum(dim=dim).unsqueeze(dim)

class FastAttention(nn.Module):
    def __init__(self, dim_heads, nb_features = None, ortho_scaling = 0, causal = False, generalized_attention = False, kernel_fn = nn.ReLU(), no_projection = False):
        super().__init__()
        nb_features = default(nb_features, int(dim_heads * math.log(dim_heads)))

        self.dim_heads = dim_heads
        self.nb_features = nb_features
        self.ortho_scaling = ortho_scaling

        self.create_projection = partial(gaussian_orthogonal_random_matrix, nb_rows = self.nb_features, nb_columns = dim_heads, scaling = ortho_scaling)
        projection_matrix = self.create_projection()
        self.register_buffer('projection_matrix', projection_matrix)

        self.generalized_attention = generalized_attention
        self.kernel_fn = kernel_fn

        # if this is turned on, no projection will be used
        # queries and keys will be softmax-ed as in the original efficient attention paper
        self.no_projection = no_projection

        self.causal = causal
        if causal:
            try:
                import fast_transformers.causal_product.causal_product_cuda
                self.causal_linear_fn = partial(causal_linear_attention)
            except ImportError:
                print('unable to import cuda code for auto-regressive Performer. will default to the memory inefficient non-cuda version')
                self.causal_linear_fn = causal_linear_attention_noncuda

    @torch.no_grad()
    def redraw_projection_matrix(self, device):
        projections = self.create_projection(device = device)
        self.projection_matrix.copy_(projections)
        del projections

    def forward(self, q, k, v, output_attentions = False):
        device = q.device
        # inds = [8060, 8064, 6243, 8575, 10342, 10913, 9366, 993, 7796, 5210, 5212, 5504, 6851, 6559, 5508, 13107, 13820]
        if self.no_projection:
            q = q.softmax(dim = -1)
            k = torch.exp(k) if self.causal else k.softmax(dim = -2)

        elif self.generalized_attention:
            create_kernel = partial(generalized_kernel, kernel_fn = self.kernel_fn, projection_matrix = self.projection_matrix, device = device)
            q, k = map(create_kernel, (q, k))

        else:
            create_kernel = partial(softmax_kernel, projection_matrix = self.projection_matrix, device = device)
            q = create_kernel(q, is_query = True)
            k = create_kernel(k, is_query = False)

        attn_fn = linear_attention if not self.causal else self.causal_linear_fn
        out = attn_fn(q, k, v)
        if output_attentions:
            v_diag = torch.eye(v.shape[-2]).to(device)
            v_diag = v_diag.unsqueeze(0).unsqueeze(0).repeat(v.shape[0],v.shape[1],1,1)
            # attn_weights = torch.zeros(1, 1, len(inds), len(inds)).to(device).to(torch.float16)
            # attn_weights = torch.zeros(1, q.shape[1], len(inds), len(inds)).to(device).to(torch.float16)
            attn_weights = torch.zeros(1, 1, q.shape[2], q.shape[2]).to(device).to(torch.float16)
            for head_dim in range(q.shape[1]):
                # attn_weights[0, head_dim] = torch.abs(attn_fn(q[:,head_dim].to(torch.float16), k[:,head_dim].to(torch.float16), v_diag[:,head_dim].to(torch.float16)))[0, inds][:, inds]
                attn_weights += torch.abs(attn_fn(q[:,head_dim].to(torch.float16), k[:,head_dim].to(torch.float16), v_diag[:,head_dim].to(torch.float16)))
                # attn_weights += norm_tensor(torch.abs(attn_fn(q[:,head_dim].to(torch.float16), k[:,head_dim].to(torch.float16), v_diag[:,head_dim].to(torch.float16))), dim=-1)
            attn_weights /= q.shape[1]
            return out, attn_weights
        else:
            return out

# classes

class ReZero(nn.Module):
    def __init__(self, fn):
        super().__init__()
        self.g = nn.Parameter(torch.tensor(1e-3))
        self.fn = fn

    def forward(self, x, **kwargs):
        return self.fn(x, **kwargs) * self.g

class PreScaleNorm(nn.Module):
    def __init__(self, dim, fn, eps=1e-5):
        super().__init__()
        self.fn = fn
        self.g = nn.Parameter(torch.ones(1))
        self.eps = eps

    def forward(self, x, **kwargs):
        n = torch.norm(x, dim=-1, keepdim=True).clamp(min=self.eps)
        x = x / n * self.g
        return self.fn(x, **kwargs)

class PreLayerNorm(nn.Module):
    def __init__(self, dim, fn):
        super().__init__()
        self.norm = nn.LayerNorm(dim)
        self.fn = fn
    def forward(self, x, **kwargs):
        return self.fn(self.norm(x), **kwargs)

class Chunk(nn.Module):
    def __init__(self, chunks, fn, along_dim = -1):
        super().__init__()
        self.dim = along_dim
        self.chunks = chunks
        self.fn = fn

    def forward(self, x, **kwargs):
        if self.chunks == 1:
            return self.fn(x, **kwargs)
        chunks = x.chunk(self.chunks, dim = self.dim)
        return torch.cat([self.fn(c, **kwargs) for c in chunks], dim = self.dim)

class FeedForward(nn.Module):
    def __init__(self, dim, mult = 4, dropout = 0., activation = None, glu = False):
        super().__init__()
        activation = default(activation, nn.GELU)

        self.glu = glu
        self.w1 = nn.Linear(dim, dim * mult * (2 if glu else 1))
        self.act = activation()
        self.dropout = nn.Dropout(dropout)
        self.w2 = nn.Linear(dim * mult, dim)

    def forward(self, x, **kwargs):
        if not self.glu:
            x = self.w1(x)
            x = self.act(x)
        else:
            x, v = self.w1(x).chunk(2, dim=-1)
            x = self.act(x) * v

        x = self.dropout(x)
        x = self.w2(x)
        return x

class SelfAttention(nn.Module):
    def __init__(
        self,
        dim,
        causal = False,
        heads = 8,
        dim_head = 64,
        local_heads = 0,
        local_window_size = 256,
        nb_features = None,
        feature_redraw_interval = 1000,
        generalized_attention = False,
        kernel_fn = nn.ReLU(),
        dropout = 0.,
        no_projection = False,
        qkv_bias = False
    ):
        super().__init__()
        assert dim % heads == 0, 'dimension must be divisible by number of heads'
        dim_head = default(dim_head, dim // heads)
        inner_dim = dim_head * heads
        self.fast_attention = FastAttention(dim_head, nb_features, causal = causal, generalized_attention = generalized_attention, kernel_fn = kernel_fn, no_projection = no_projection)

        self.heads = heads
        self.global_heads = heads - local_heads
        self.local_attn = LocalAttention(window_size = local_window_size, causal = causal, autopad = True, dropout = dropout, look_forward = int(not causal), rel_pos_emb_config = (dim_head, local_heads)) if local_heads > 0 else None

        self.to_q = nn.Linear(dim, inner_dim, bias = qkv_bias)
        self.to_k = nn.Linear(dim, inner_dim, bias = qkv_bias)
        self.to_v = nn.Linear(dim, inner_dim, bias = qkv_bias)
        self.to_out = nn.Linear(inner_dim, dim)
        self.dropout = nn.Dropout(dropout)

    def forward(self, x, pos_emb = None, context = None, mask = None, context_mask = None, output_attentions = False, **kwargs):
        b, n, _, h, gh = *x.shape, self.heads, self.global_heads

        cross_attend = exists(context)

        context = default(context, x)
        context_mask = default(context_mask, mask) if not cross_attend else context_mask

        q, k, v = self.to_q(x), self.to_k(context), self.to_v(context)

        q, k, v = map(lambda t: rearrange(t, 'b n (h d) -> b h n d', h = h), (q, k, v))
        (q, lq), (k, lk), (v, lv) = map(lambda t: (t[:, :gh], t[:, gh:]), (q, k, v))

        attn_outs = []

        if not empty(q):
            if exists(context_mask):
                global_mask = context_mask[:, None, :, None]
                v.masked_fill_(~global_mask, 0.)

            if exists(pos_emb) and not cross_attend:
                q, k, = apply_rotary_pos_emb(q, k, pos_emb)

            if output_attentions:
                out, attn_weights = self.fast_attention(q, k, v, output_attentions)
            else:
                out = self.fast_attention(q, k, v)
            attn_outs.append(out)

        if not empty(lq):
            assert not cross_attend, 'local attention is not compatible with cross attention'
            out = self.local_attn(lq, lk, lv, input_mask = mask)
            attn_outs.append(out)

        out = torch.cat(attn_outs, dim = 1)     # combine attn_out and cross_attn_out, here we have only attn_out, that means this line does nothing
        out = rearrange(out, 'b h n d -> b n (h d)')
        out =  self.to_out(out)
        if output_attentions:
            return self.dropout(out), attn_weights
        else:
            return self.dropout(out)
        
    # performer
class Performer(nn.Module):
    def __init__(
        self,
        dim,                                # dimension
        depth,                              # layers
        heads,                              # heads
        dim_head,                           # dim of head
        local_attn_heads = 0,               # num of local attention heads, (heads - local_attn_heads) is num of global performers
        local_window_size = 256,            # window size of local attention
        causal = False,                     # autoregressive or not
        ff_mult = 4,                        # dim of intermediate features after attention / dim of input features
        nb_features = None,                 # number of random features, if not set, will default to (d * log(d)), where d is the dimension of each head ?? what is random feature ??
        feature_redraw_interval = 1000,     # how frequently to redraw the projection matrix, the more frequent, the slower the training
        reversible = False,                 # reversible layers, from Reformer (save memory)
        ff_chunks = 1,                      # chunk feedforward layer, from Reformer
        generalized_attention = False,      # defaults to softmax approximation, but can be set to True for generalized attention ?? what is generalized attention ??
        kernel_fn = nn.ReLU(),              # the kernel function to be used, if generalized attention is turned on, defaults to Relu
        use_scalenorm = False,              # use scale norm, from 'Transformers without Tears' paper, a substitute for LayerNorm, priority: scalenorm.rezero.layernorm
        use_rezero = False,                 # use Rezero or not, from 'Rezero is all you need' paper, a substitute for LayerNorm, priority: scalenorm.rezero.layernorm
        ff_glu = False,                     # use GLU (Gated Linear Units) variant for feedforward
        ff_dropout = 0.,                    # feedforward dropout
        attn_dropout = 0.,                  # post-attention dropout
        cross_attend = False,               # ??
        no_projection = False,              # ??
        auto_check_redraw = True,           # ??
        qkv_bias = True,                    # ??
    ):
        super().__init__()
        layers = nn.ModuleList([])
        local_attn_heads = cast_tuple(local_attn_heads)
        local_attn_heads = local_attn_heads * depth if len(local_attn_heads) == 1 else local_attn_heads
        assert len(local_attn_heads) == depth, 'tuple specifying number of local attention heads per depth must be equal to the total depth'
        assert all(map(lambda n: n >= 0 and n <= heads, local_attn_heads)), 'local attention head value must be less than the total number of heads'

        if use_scalenorm:
            wrapper_fn = partial(PreScaleNorm, dim)
        elif use_rezero:
            wrapper_fn = ReZero
        else:
            wrapper_fn = partial(PreLayerNorm, dim)

        for _, local_heads in zip(range(depth), local_attn_heads):
            layers.append(nn.ModuleList([
                wrapper_fn(SelfAttention(dim, causal = causal, heads = heads, dim_head = dim_head, local_heads = local_heads, local_window_size = local_window_size, nb_features = nb_features, generalized_attention = generalized_attention, kernel_fn = kernel_fn, dropout = attn_dropout, no_projection = no_projection, qkv_bias = qkv_bias)),
                wrapper_fn(Chunk(ff_chunks, FeedForward(dim, mult = ff_mult, dropout = ff_dropout, glu = ff_glu), along_dim = 1))
            ]))
            # if no need cross_attend(decoder), begin next cycle
            if not cross_attend:
                continue
            layers.append(nn.ModuleList([
                wrapper_fn(SelfAttention(dim, heads = heads, dim_head = dim_head, nb_features = nb_features, generalized_attention = generalized_attention, kernel_fn = kernel_fn, dropout = attn_dropout, no_projection = no_projection)),
                wrapper_fn(Chunk(ff_chunks, FeedForward(dim, mult = ff_mult, dropout = ff_dropout, glu = ff_glu), along_dim = 1))
            ]))

        execute_type = ReversibleSequence if reversible else SequentialSequence

        route_attn = ((True, False),) * depth * (2 if cross_attend else 1)  # ((True, False), (True, False), (True, False), (True, False), (True, False), (True, False))
        route_context = ((False, False), (True, False)) * depth
        attn_route_map = {'mask': route_attn, 'pos_emb': route_attn}
        context_route_map = {'context': route_context, 'context_mask': route_context} if cross_attend else {}
        self.net = execute_type(layers, args_route = {**attn_route_map, **context_route_map})

        # keeping track of when to redraw projections for all attention layers
        self.auto_check_redraw = auto_check_redraw
        self.feature_redraw_interval = feature_redraw_interval
        self.register_buffer('calls_since_last_redraw', torch.tensor(0))

    def fix_projection_matrices_(self):
        self.feature_redraw_interval = None

    def check_redraw_projections(self):
        if not self.training:
            return

        if exists(self.feature_redraw_interval) and self.calls_since_last_redraw >= self.feature_redraw_interval:
            device = get_module_device(self)

            fast_attentions = find_modules(self, FastAttention)
            for fast_attention in fast_attentions:
                fast_attention.redraw_projection_matrix(device)

            self.calls_since_last_redraw.zero_()
            return

        self.calls_since_last_redraw += 1

    def forward(self, x, output_attentions = False, **kwargs):
        if self.auto_check_redraw:
            self.check_redraw_projections()
        out = self.net(x, output_attentions = output_attentions, **kwargs)
        return out
```
