## Supplementary material for "EpiFoundation: A Foundation Model for Single-Cell ATAC-seq via Peak-to-Gene Alignment": Code for EpiFoundation: reversible.py

```
import torch
import torch.nn as nn
from operator import itemgetter
from torch.autograd.function import Function
from torch.utils.checkpoint import get_device_states, set_device_states

# for routing arguments into the functions of the reversible layer
def route_args(router, args, depth):
    routed_args = [(dict(), dict()) for _ in range(depth)]
    matched_keys = [key for key in args.keys() if key in router]

    for key in matched_keys:
        val = args[key]
        for depth, ((f_args, g_args), routes) in enumerate(zip(routed_args, router[key])):
            new_f_args, new_g_args = map(lambda route: ({key: val} if route else {}), routes)
            routed_args[depth] = ({**f_args, **new_f_args}, {**g_args, **new_g_args})
    return routed_args

# following example for saving and setting rng here https://pytorch.org/docs/stable/_modules/torch/utils/checkpoint.html
class Deterministic(nn.Module):
    def __init__(self, net):
        super().__init__()
        self.net = net
        self.cpu_state = None
        self.cuda_in_fwd = None
        self.gpu_devices = None
        self.gpu_states = None

    def record_rng(self, *args):
        self.cpu_state = torch.get_rng_state()
        if torch.cuda._initialized:
            self.cuda_in_fwd = True
            self.gpu_devices, self.gpu_states = get_device_states(*args)

    def forward(self, *args, record_rng = False, set_rng = False, **kwargs):
        if record_rng:
            self.record_rng(*args)

        if not set_rng:
            return self.net(*args, **kwargs)

        rng_devices = []
        if self.cuda_in_fwd:
            rng_devices = self.gpu_devices

        with torch.random.fork_rng(devices=rng_devices, enabled=True):
            torch.set_rng_state(self.cpu_state)
            if self.cuda_in_fwd:
                set_device_states(self.gpu_devices, self.gpu_states)
            return self.net(*args, **kwargs)

# heavily inspired by https://github.com/RobinBruegger/RevTorch/blob/master/revtorch/revtorch.py
# once multi-GPU is confirmed working, refactor and send PR back to source
class ReversibleBlock(nn.Module):
    def __init__(self, f, g):
        super().__init__()
        self.f = Deterministic(f)
        self.g = Deterministic(g)

    def forward(self, x, f_args = {}, g_args = {}):
        x1, x2 = torch.chunk(x, 2, dim=2)
        y1, y2 = None, None

        with torch.no_grad():
            y1 = x1 + self.f(x2, record_rng=self.training, **f_args)
            y2 = x2 + self.g(y1, record_rng=self.training, **g_args)

        return torch.cat([y1, y2], dim=2)

    def backward_pass(self, y, dy, f_args = {}, g_args = {}):
        y1, y2 = torch.chunk(y, 2, dim=2)
        del y

        dy1, dy2 = torch.chunk(dy, 2, dim=2)
        del dy

        with torch.enable_grad():
            y1.requires_grad = True
            gy1 = self.g(y1, set_rng=True, **g_args)
            torch.autograd.backward(gy1, dy2)

        with torch.no_grad():
            x2 = y2 - gy1
            del y2, gy1

            dx1 = dy1 + y1.grad
            del dy1
            y1.grad = None

        with torch.enable_grad():
            x2.requires_grad = True
            fx2 = self.f(x2, set_rng=True, **f_args)
            torch.autograd.backward(fx2, dx1, retain_graph=True)

        with torch.no_grad():
            x1 = y1 - fx2
            del y1, fx2

            dx2 = dy2 + x2.grad
            del dy2
            x2.grad = None

            x = torch.cat([x1, x2.detach()], dim=2)
            dx = torch.cat([dx1, dx2], dim=2)

        return x, dx

class _ReversibleFunction(Function):
    @staticmethod
    def forward(ctx, x, blocks, args):
        ctx.args = args
        for block, kwarg in zip(blocks, args):
            x = block(x, **kwarg)
        ctx.y = x.detach()
        ctx.blocks = blocks
        return x

    @staticmethod
    def backward(ctx, dy):
        y = ctx.y
        args = ctx.args
        for block, kwargs in zip(ctx.blocks[::-1], args[::-1]):
            y, dy = block.backward_pass(y, dy, **kwargs)
        return dy, None, None

class SequentialSequence(nn.Module):
    def __init__(self, layers, args_route = {}):
        super().__init__()
        assert all(len(route) == len(layers) for route in args_route.values()), 'each argument route map must have the same depth as the number of sequential layers'
        self.layers = layers
        self.args_route = args_route

    def forward(self, x, output_attentions = False, **kwargs):
        args = route_args(self.args_route, kwargs, len(self.layers))
        layers_and_args = list(zip(self.layers, args))

        if output_attentions:
            attn_weights = []
        for (f, g), (f_args, g_args) in layers_and_args:
            if output_attentions:
                x = x + f(x, output_attentions = output_attentions, **f_args)[0]
                attn_weights.append(f(x, output_attentions = output_attentions, **f_args)[1].unsqueeze(0))
            else:
                x = x + f(x, **f_args)
            x = x + g(x, **g_args)
        if output_attentions:
            attn_weights = torch.transpose(torch.cat(attn_weights, dim=0), 0, 1)    # the final dim is (batch, layer, head, len, len)
            attn_weights = torch.mean(attn_weights, dim=1)                        # the dim is (batch, head, len, len)
            return x, attn_weights
        else:
            return x

class ReversibleSequence(nn.Module):
    def __init__(self, blocks, args_route = {}):
        super().__init__()
        self.args_route = args_route
        self.blocks = nn.ModuleList([ReversibleBlock(f=f, g=g) for f, g in blocks])

    def forward(self, x, **kwargs):
        x = torch.cat([x, x], dim=-1)

        blocks = self.blocks
        args = route_args(self.args_route, kwargs, len(blocks))
        args = list(map(lambda x: {'f_args': x[0], 'g_args': x[1]}, args))

        out =  _ReversibleFunction.apply(x, blocks, args)
        return torch.stack(out.chunk(2, dim=-1)).sum(dim=0)
```
