## Supplementary material for "EpiFoundation: A Foundation Model for Single-Cell ATAC-seq via Peak-to-Gene Alignment": Code for EpiFoundation: transformer.py

```
import gc
import math
from typing import Dict, Mapping, Optional, Tuple, Any, Union

import torch
import numpy as np
from torch import nn, Tensor
import torch.distributed as dist
import torch.nn.functional as F
from torch.nn import TransformerEncoder, TransformerEncoderLayer
from torch.distributions import Bernoulli
from tqdm import trange
from model.flashDiff import MultiheadFlashDiff

try:
    from model.flashMHA import FlashMHA
except ImportError:
    import warnings

    warnings.warn("flash_attn is not installed")
    flash_attn_available = False


class TransformerModel(nn.Module):
    def __init__(
        self,
        d_model: int,
        nhead: int,
        d_hid: int,
        nlayers: int,
        dropout: float = 0.5,
        use_fast_transformer: bool = True,
        fast_transformer_backend: str = "flash",
    ):
        super().__init__()


        if use_fast_transformer:
            if fast_transformer_backend == "linear":
                self.transformer_encoder = FastTransformerEncoderWrapper(
                    d_model, nhead, d_hid, nlayers, dropout
                )
            elif fast_transformer_backend == "flash":
                encoder_layers = FlashTransformerEncoderLayer(
                    d_model=d_model,
                    nhead=nhead,
                    dim_feedforward=d_hid,
                    dropout=dropout,
                    batch_first=True
                )
                self.transformer_encoder = TransformerEncoder(encoder_layers, nlayers, enable_nested_tensor= False)
            elif fast_transformer_backend == "diff":
                encoder_layers = DiffTransformerEncoderLayer(
                    d_model=d_model,
                    nhead=nhead,
                    dim_feedforward=d_hid,
                    dropout=dropout,
                    batch_first=True
                )
                self.transformer_encoder = TransformerEncoder(encoder_layers, nlayers, enable_nested_tensor= False)
        else:
            encoder_layers = TransformerEncoderLayer(
                d_model, nhead, d_hid, dropout, batch_first=True
            )
            self.transformer_encoder = TransformerEncoder(encoder_layers, nlayers)

    def forward(
        self,
        embs: Tensor,
        src_key_padding_mask: Optional[Tensor] = None,
        need_weights: Optional[bool] = False,
    ) -> Tensor:
        if need_weights:
            output, layer_weights = self.transformer_encoder(embs, src_key_padding_mask=src_key_padding_mask, need_weights=need_weights)
            return output, layer_weights
        else:
            output = self.transformer_encoder(embs, src_key_padding_mask=src_key_padding_mask)
            return output  # (batch, seq_len, embsize)


class FastTransformerEncoderWrapper(nn.Module):
    def __init__(
        self,
        d_model: int,
        nhead: int,
        d_hid: int,
        nlayers: int,
        dropout: float = 0.5,
    ):
        super().__init__()
        self.fast_transformer_encoder = self.build_fast_transformer_encoder(
            d_model, nhead, d_hid, nlayers, dropout
        )

    @staticmethod
    def build_fast_transformer_encoder(
        d_model: int, nhead: int, d_hid: int, nlayers: int, dropout: float
    ) -> nn.Module:
        from fast_transformers.builders import TransformerEncoderBuilder

        if d_model % nhead != 0:
            raise ValueError(
                f"d_model must be divisible by nhead, "
                f"got d_model={d_model} and nhead={nhead}"
            )
        builder = TransformerEncoderBuilder.from_kwargs(
            n_layers=nlayers,
            n_heads=nhead,
            query_dimensions=d_model // nhead,
            value_dimensions=d_model // nhead,
            feed_forward_dimensions=d_hid,
            attention_type="linear",
            attention_dropout=dropout,
            dropout=dropout,
            activation="gelu",
        )
        assert builder.attention_type == "linear"
        return builder.get()

    @staticmethod
    def build_length_mask(
        src: Tensor,
        src_key_padding_mask: torch.BoolTensor,
    ) -> "LengthMask":
        from fast_transformers.masking import LengthMask

        seq_len = src.shape[1]
        num_paddings = src_key_padding_mask.sum(dim=1)
        actual_seq_len = seq_len - num_paddings  # (N,)
        length_mask = LengthMask(actual_seq_len, max_len=seq_len, device=src.device)

        if src_key_padding_mask[length_mask.bool_matrix].sum() != 0:
            raise ValueError(
                "Found padding tokens in the middle of the sequence. "
                "src_key_padding_mask and length_mask are not compatible."
            )
        return length_mask

    def forward(
        self,
        src: Tensor,
        src_key_padding_mask: torch.BoolTensor,
    ) -> Tensor:
        """
        Args:
            src: Tensor, shape [N, seq_len, embsize]
            src_key_padding_mask: Tensor, shape [N, seq_len]

        Returns:
            output Tensor of shape [N, seq_len, embsize]
        """
        if src_key_padding_mask.shape != src.shape[:2]:
            raise ValueError(
                f"src_key_padding_mask shape {src_key_padding_mask.shape} "
                f"does not match first two dims of src shape {src.shape[:2]}"
            )

        if src_key_padding_mask.dtype != torch.bool:
            raise ValueError(
                f"src_key_padding_mask needs to be of type torch.bool, "
                f"got {src_key_padding_mask.dtype}"
            )

        length_mask = self.build_length_mask(src, src_key_padding_mask)
        output = self.fast_transformer_encoder(src, length_mask=length_mask)
        return output


class FlashTransformerEncoderLayer(nn.Module):
    r"""TransformerEncoderLayer is made up of self-attn and feedforward network.
    The class is modified from torch.nn.TransformerEncoderLayer to support the
    FlashAttention.

    Args:
        d_model: the number of expected features in the input (required).
        nhead: the number of heads in the multiheadattention models (required).
        dim_feedforward: the dimension of the feedforward network model (default=2048).
        dropout: the dropout value (default=0.1).
        activation: the activation function of intermediate layer, relu or gelu (default=relu).
        layer_norm_eps: the eps value in layer normalization components (default=1e-5).
        batch_first: If ``True``, then the input and output tensors are provided
            as (batch, seq, feature). Default: ``False``.

    Examples::
        >>> encoder_layer = nn.TransformerEncoderLayer(d_model=512, nhead=8)
        >>> src = torch.rand(10, 32, 512)
        >>> out = encoder_layer(src)

    Alternatively, when ``batch_first`` is ``True``:
        >>> encoder_layer = nn.TransformerEncoderLayer(d_model=512, nhead=8, batch_first=True)
        >>> src = torch.rand(32, 10, 512)
        >>> out = encoder_layer(src)
    """
    __constants__ = ["batch_first"]

    def __init__(
        self,
        d_model,
        nhead,
        dim_feedforward=2048,
        dropout=0.1,
        activation="relu",
        layer_norm_eps=1e-5,
        batch_first=True,
        device=None,
        dtype=None,
        norm_scheme="post",  # "pre" or "post"
    ) -> None:
        factory_kwargs = {"device": device, "dtype": dtype}
        super().__init__()
        self.self_attn = FlashMHA(
            embed_dim=d_model,
            num_heads=nhead,
            batch_first=batch_first,
            attention_dropout=dropout,
            **factory_kwargs,
        )
        # self.self_attn = FlashMHA(
        #     embed_dim=d_model,
        #     num_heads=nhead,
        #     dropout=dropout,
        #     use_flash_attn= True,
        #     **factory_kwargs,
        # )
        # Version compatibility workaround
        if not hasattr(self.self_attn, "batch_first"):
            self.self_attn.batch_first = batch_first
        # Implementation of Feedforward model
        self.linear1 = nn.Linear(d_model, dim_feedforward, **factory_kwargs)
        self.dropout = nn.Dropout(dropout)
        self.linear2 = nn.Linear(dim_feedforward, d_model, **factory_kwargs)

        self.norm1 = nn.LayerNorm(d_model, eps=layer_norm_eps, **factory_kwargs)
        self.norm2 = nn.LayerNorm(d_model, eps=layer_norm_eps, **factory_kwargs)
        self.dropout1 = nn.Dropout(dropout)
        self.dropout2 = nn.Dropout(dropout)

        self.activation = self._get_activation_fn(activation)
        self.norm_scheme = norm_scheme
        if self.norm_scheme not in ["pre", "post"]:
            raise ValueError(f"norm_scheme should be pre or post, not {norm_scheme}")

    @staticmethod
    def _get_activation_fn(activation):
        if activation == "relu":
            return F.relu
        elif activation == "gelu":
            return F.gelu

        raise RuntimeError("activation should be relu/gelu, not {}".format(activation))

    def __setstate__(self, state):
        if "activation" not in state:
            state["activation"] = F.relu
        super().__setstate__(state)

    def forward(
        self,
        src: Tensor,
        src_mask: Optional[Tensor] = None,
        src_key_padding_mask: Optional[Tensor] = None,
        need_weights: Optional[bool] = False,
        **kwargs,
    ) -> Tensor:
        r"""Pass the input through the encoder layer.

        Args:
            src: the sequence to the encoder layer (required).
            src_mask: the mask for the src sequence (optional).
            src_key_padding_mask: the mask for the src keys per batch (optional).

        Shape:
            see the docs in Transformer class.
        """
        if src_mask is not None:
            raise ValueError("FlashTransformerEncoderLayer does not support src_mask")

        if src_key_padding_mask is not None:
            if not src_key_padding_mask.any().item():
                # no padding tokens in src
                src_key_padding_mask_ = None
            else:
                if src_key_padding_mask.dtype != torch.bool:
                    src_key_padding_mask = src_key_padding_mask.bool()
                # NOTE: the FlashMHA uses mask 0 for padding tokens, which is the opposite
                src_key_padding_mask_ = ~src_key_padding_mask
        else:
            src_key_padding_mask_ = None

        if self.norm_scheme == "pre":
            src = self.norm1(src)
            src2, atten_weight = self.self_attn(src, key_padding_mask=src_key_padding_mask_, need_weights = need_weights)

            src = src + self.dropout1(src2)
            src = self.norm2(src)
            src2 = self.linear2(self.dropout(self.activation(self.linear1(src))))
            src = src + self.dropout2(src2)
        else:
            src2, atten_weight = self.self_attn(src, key_padding_mask=src_key_padding_mask_, need_weights = need_weights)
            src = src + self.dropout1(src2)
            src = self.norm1(src)
            src2 = self.linear2(self.dropout(self.activation(self.linear1(src))))
            src = src + self.dropout2(src2)
            src = self.norm2(src)

        if need_weights:
            return src, atten_weight
        else:
            return src


class DiffTransformerEncoderLayer(nn.Module):
    __constants__ = ["batch_first"]

    def __init__(
        self,
        d_model,
        nhead,
        dim_feedforward=2048,
        dropout=0.1,
        lambda_init = 0.8,
        activation="relu",
        layer_norm_eps=1e-5,
        batch_first=True,
        device=None,
        dtype=None,
        norm_scheme="post",  # "pre" or "post"
    ) -> None:
        factory_kwargs = {"device": device, "dtype": dtype}
        super().__init__()
        self.self_attn = MultiheadFlashDiff(
            embed_dim=d_model,
            num_heads=nhead,
            attention_dropout=dropout,
            lambda_init = lambda_init,
        )
        # self.self_attn = FlashMHA(
        #     embed_dim=d_model,
        #     num_heads=nhead,
        #     dropout=dropout,
        #     use_flash_attn= True,
        #     **factory_kwargs,
        # )
        # Version compatibility workaround
        if not hasattr(self.self_attn, "batch_first"):
            self.self_attn.batch_first = batch_first
        # Implementation of Feedforward model
        self.linear1 = nn.Linear(d_model, dim_feedforward, **factory_kwargs)
        self.dropout = nn.Dropout(dropout)
        self.linear2 = nn.Linear(dim_feedforward, d_model, **factory_kwargs)

        self.norm1 = nn.LayerNorm(d_model, eps=layer_norm_eps, **factory_kwargs)
        self.norm2 = nn.LayerNorm(d_model, eps=layer_norm_eps, **factory_kwargs)
        self.dropout1 = nn.Dropout(dropout)
        self.dropout2 = nn.Dropout(dropout)

        self.activation = self._get_activation_fn(activation)
        self.norm_scheme = norm_scheme
        if self.norm_scheme not in ["pre", "post"]:
            raise ValueError(f"norm_scheme should be pre or post, not {norm_scheme}")

    @staticmethod
    def _get_activation_fn(activation):
        if activation == "relu":
            return F.relu
        elif activation == "gelu":
            return F.gelu

        raise RuntimeError("activation should be relu/gelu, not {}".format(activation))

    def __setstate__(self, state):
        if "activation" not in state:
            state["activation"] = F.relu
        super().__setstate__(state)

    def forward(
        self,
        src: Tensor,
        src_mask: Optional[Tensor] = None,
        src_key_padding_mask: Optional[Tensor] = None,
        need_weights: Optional[bool] = False,
        **kwargs,
    ) -> Tensor:
        r"""Pass the input through the encoder layer.

        Args:
            src: the sequence to the encoder layer (required).
            src_mask: the mask for the src sequence (optional).
            src_key_padding_mask: the mask for the src keys per batch (optional).

        Shape:
            see the docs in Transformer class.
        """
        if src_mask is not None:
            raise ValueError("FlashTransformerEncoderLayer does not support src_mask")

        if src_key_padding_mask is not None:
            if not src_key_padding_mask.any().item():
                # no padding tokens in src
                src_key_padding_mask_ = None
            else:
                if src_key_padding_mask.dtype != torch.bool:
                    src_key_padding_mask = src_key_padding_mask.bool()
                # NOTE: the FlashMHA uses mask 0 for padding tokens, which is the opposite
                src_key_padding_mask_ = ~src_key_padding_mask
        else:
            src_key_padding_mask_ = None

        if self.norm_scheme == "pre":
            src = self.norm1(src)
            src2 = self.self_attn(src)

            src = src + self.dropout1(src2)
            src = self.norm2(src)
            src2 = self.linear2(self.dropout(self.activation(self.linear1(src))))
            src = src + self.dropout2(src2)
        else:
            src2 = self.self_attn(src)
            src = src + self.dropout1(src2)
            src = self.norm1(src)
            src2 = self.linear2(self.dropout(self.activation(self.linear1(src))))
            src = src + self.dropout2(src2)
            src = self.norm2(src)

        if need_weights:
            return src, atten_weight
        else:
            return src
```
