## Supplementary material for "EpiFoundation: A Foundation Model for Single-Cell ATAC-seq via Peak-to-Gene Alignment": Code for EpiFoundation: prepare_data.py

```
import scanpy as sc
from scanpy import AnnData
import scipy
import os
from model import logger
import numpy as np
from data.preprocess import Preprocessor
from tokenizer import GeneVocab
from data.dataloader import *
import yaml
from tqdm import tqdm
    

def divide_data(data: AnnData, radio: dict = {'train': 0.9, 'test': 0.05, 'valid': 0.05}):
    '''
    Divide the data into train, test, valid.
    return 3 AnnData object.
    '''
    assert sum(radio.values()) == 1.0, 'The sum of radio should be 1.0'
    data = data.copy()
    # get the number of cells
    n_cells = data.shape[0]
    # get the index of cells
    idx = np.arange(n_cells)
    np.random.shuffle(idx)
    # get the number of cells for each part
    n_train = int(n_cells * radio['train'])
    n_test = int(n_cells * radio['test'])
    n_valid = int(n_cells * radio['valid'])
    # divide the data
    train_data = data[idx[:n_train]]
    test_data = data[idx[n_train:n_train+n_test]]
    valid_data = data[idx[n_train+n_test:]]
    
    return train_data, test_data, valid_data

def preprocess():
    
    preprocess_config = {
        'path': '/home/jwu418/workspace/data/ours/',
        'raw_data': 'pbmc_rna_s1.h5ad',
        'use_key': 'X',
        'filter_gene_by_counts': False,
        'filter_cell_by_counts': False,
        'normalize_total': False,
        'result_normed_key': 'X_normed',
        'log1p': False,
        'result_log1p_key': 'X_log1p',
        'subset_hvg': False,
        'hvg_use_key': None,
        'hvg_flavor': 'seurat_v3',
        'binning': [2],
        'result_binned_key': 'X_binned',
        'batch_key': 'batch',
        'output_name': 'pbmc_rna_s1',
    }
    file = '{}/raw/{}'.format(preprocess_config['path'], preprocess_config['raw_data'])
    adata = sc.read_h5ad(file)
    # devide data into train, test, valid. with 0.8,0.1,0.1
    # adata._raw._var.rename(columns={'_index': 'genes'}, inplace=True)
    print(adata)
    
    train_data, test_data, valid_data = divide_data(adata)
    for binning in preprocess_config['binning']:
        logger.info('Binning: {}'.format(binning))
        processor = Preprocessor(use_key=preprocess_config['use_key'],
                                    filter_gene_by_counts=preprocess_config['filter_gene_by_counts'],
                                    filter_cell_by_counts=preprocess_config['filter_cell_by_counts'],
                                    normalize_total=preprocess_config['normalize_total'],
                                    result_normed_key=preprocess_config['result_normed_key'],
                                    log1p=preprocess_config['log1p'],
                                    result_log1p_key=preprocess_config['result_log1p_key'],
                                    subset_hvg=preprocess_config['subset_hvg'],
                                    hvg_use_key=preprocess_config['hvg_use_key'],
                                    hvg_flavor=preprocess_config['hvg_flavor'],
                                    binning=binning,
                                    result_binned_key=preprocess_config['result_binned_key'])
        
        
        
        output_name = f'{preprocess_config["output_name"]}_binning_{binning}'
        
        logger.info('Preprocessing Train Data')
        processor(train_data, batch_key= preprocess_config['batch_key'])
        print(train_data)
        train_data.write('{}/train/{}.h5ad'.format(preprocess_config['path'], output_name))

        logger.info('Preprocessing test Data')
        processor(test_data, batch_key= preprocess_config['batch_key'])
        print(test_data)
        test_data.write('{}/test/{}.h5ad'.format(preprocess_config['path'], output_name))
        
        logger.info('Preprocessing valid Data')
        processor(valid_data, batch_key= preprocess_config['batch_key'])
        print(valid_data)
        valid_data.write('{}/valid/{}.h5ad'.format(preprocess_config['path'], output_name))
    
    
    # save preprocess config as a yml file
    with open('/home/jwu418/workspace/data/ours/configs/{}.yml'.format(output_name), 'w') as file:
        yaml.dump(preprocess_config, file)
        
        
def reduce_data():
    path = '/home/jwu418/workspace/data/ours'
    rna_file = 'pbmc_rna_s1_binning_2.h5ad'
    
    stage = ['test', 'valid', 'train']
    for s in stage:
        adata = sc.read_h5ad('{}/{}/{}'.format(path, s, rna_file))
        print('Before:', adata)
        # remove the adata.raw
        adata.raw = None
        # save the X_binned as X
        adata.X = adata.layers['X_binned']
        # save adata.X as sparse matrix
        adata.X = scipy.sparse.csr_matrix(adata.X)
        # remove the X_binned layer
        adata.layers.pop('X_binned')
        # save the data as a new file
        adata.write('{}/{}/{}'.format(path, s, 'pbmc_rna_s1_binning_2_reduced.h5ad'))
        
def get_pair_data():
    path = '/home/jwu418/workspace/data/ours'
    rna_file = 'pbmc_rna_s1_binning_2.h5ad'
    atac_file = 'raw/pbmc_atac_s1.h5ad'
    
    output_name = 'pbmc_rna_s1_atac_paired.h5ad'
    
    stage = ['test', 'valid', 'train']
    
    atac = sc.read_h5ad('{}/{}'.format(path, atac_file))
    # breakpoint()
    for s in stage:
        rna = sc.read_h5ad('{}/{}/{}'.format(path, s, rna_file), backed='r')
        print('rna:', rna)
        # get the cell name of rna data

        rna_cell_name = rna.obs_names.tolist()
        # find the corresponding cell in atac data
        atac_cell = atac[rna_cell_name]
        # atac_cell._raw._var.rename(columns={'_index': 'peaks'}, inplace=True)
        # save the atac data as a new file
        atac_cell.write('{}/{}/{}'.format(path, s, output_name))
        print('atac cell:', atac_cell)
        
def generate_chr_vocab():
    file = '/home/jwu418/workspace/data/ours/meta/genes.csv'
    import pandas as pd
    # read the 'seqnames' column
    df = pd.read_csv(file)
    chr_names = df['seqnames'].tolist()
    chr_names = list(set(chr_names))
    vocab = GeneVocab(gene_list_or_vocab=chr_names,
                      special_first= True,
                       specials=['<pad>', '<mask>','<cls>', '<eos>'])
    vocab.save_json('/home/jwu418/workspace/data/ours/chr_vocab.json')

    
def generate_vocab():
    file = '/home/jwu418/workspace/data/ours/raw/mini_atlas_atac.h5ad'
    adata = sc.read_h5ad(file)
    # adata._raw._var.rename(columns={'_index': 'features'}, inplace=True)
    
    # get the gene names
    gene_names = adata.var_names.tolist()
    vocab = GeneVocab(gene_list_or_vocab=gene_names,
                        special_first= True,
                        specials=['<pad>', '<mask>','<cls>', '<eos>'])
    vocab.save_json('/home/jwu418/workspace/data/ours/atac_vocab.json')
    
def generate_cell_type_vocab():
    file = '/home/jwu418/workspace/data/ours/raw/pbmc_rna_s1.h5ad'
    adata = sc.read_h5ad(file)
    print(adata)
    # get the gene names
    gene_names = adata.obs['batch'].tolist()
    
    # remove duplicates
    gene_names = list(set(gene_names))
    print(gene_names)
    # get number of cell types
    print('Number of cell types:', len(gene_names))
    vocab = GeneVocab(gene_list_or_vocab=gene_names)
    vocab.save_json('/home/jwu418/workspace/data/ours/vocab/pbmc_s1_batch_vocab.json')
    
    

# if __name__ == '__main__':
    # preprocess()
    # reduce_data()
    # get_pair_data()
    # generate_vocab()
    # generate_cell_type_vocab()
    
'''
Proceesing the data:
1. preprocess()
2. get_pair_data()
3. generate_vocab()
4. generate_cell_type_vocab()
5. reduce_data()
'''
```
