## Supplementary material for "EpiFoundation: A Foundation Model for Single-Cell ATAC-seq via Peak-to-Gene Alignment": Code for EpiFoundation: pretrain_fsdp.py

```
# -*- coding: utf-8 -*-

import os
import gc
import argparse
import json
import random
import math
import random
from functools import reduce
import numpy as np
import pandas as pd
from scipy import sparse
from sklearn.model_selection import train_test_split
import torch
from torch import nn
from torch.optim import Adam
from torch.nn import functional as F
from tensorboardX import SummaryWriter
from torch.utils.data import DataLoader, Dataset
from torch.utils.data.distributed import DistributedSampler
import torch.distributed as dist

# DDP imports
from torch.distributed.fsdp import FullyShardedDataParallel as FSDP
# FSDP imports
from torch.distributed.fsdp.fully_sharded_data_parallel import (
    CPUOffload, MixedPrecision, BackwardPrefetch, ShardingStrategy
)
from torch.distributed.fsdp.wrap import (
    size_based_auto_wrap_policy,
    transformer_auto_wrap_policy,
    enable_wrap,
    wrap
)
from torch.distributed.fsdp import (
    FullStateDictConfig,
    StateDictType,
)


from model import EpiFoundation
from loss.loss import MaskedMSELoss
from data.dataloader import *
from tokenizer import GeneVocab
import scanpy as sc
import anndata as ad
from utils import *
from memory_profiler import profile

import yaml

torch.autograd.set_detect_anomaly(True)

parser = argparse.ArgumentParser()
parser.add_argument("--config", type=str, default='./configs/pretrain/atac_cross_debug.yml', help='Config file.')
args = parser.parse_args()

def main():
    dist.init_process_group(backend='nccl')
    # read and parse config file
    local_rank = int(os.environ["LOCAL_RANK"])
    rank = int(os.environ["RANK"])
    
    with open(args.config, 'r') as f:
        config = yaml.load(f, Loader=yaml.FullLoader)
        
    
    train_config = config['train']
    valid_config = config['valid']
    data_config = config['data']
    vocab_config = config['vocab']
    task_name = config['task_name']
    task_floder = './experiment/{}'.format(task_name)
    ckpt_dir = os.path.join(task_floder, 'ckpts')
    

    
    random_seed = train_config['seed']
    EPOCHS = train_config['epochs']
    BATCH_SIZE = train_config['batch_size']
    GRADIENT_ACCUMULATION = train_config['gradient_accumulation_steps']
    LEARNING_RATE = float(train_config['lr'])

    model_name = train_config['model']['encoder']
    
    save_ckpt_freq = train_config['save_ckpt_freq'] if 'save_ckpt_freq' in train_config else 5
    resume = train_config['resume'] if 'resume' in train_config else False
    
    # special tokens
    pad = vocab_config['special_tokens']['pad']
    mask = vocab_config['special_tokens']['mask']
    cls = vocab_config['special_tokens']['cls']
    
    # distibuted setting, use local_rank as device id
    torch.cuda.set_device(local_rank)
    device = torch.device("cuda", local_rank)
    
    world_size = dist.get_world_size()
    seed_all(random_seed + rank)
    # use rank to determine if it is master node, instead of local_rank
    is_master = (rank == 0)
    
    # init loggers
    logger = set_log(log_dir= os.path.join(task_floder, 'logs'))
    tb_logger = SummaryWriter(os.path.join(task_floder, 'tb_logs'))
    if is_master:
        logger.info(dict2str(config))
    
    
    rna_vocab = GeneVocab.from_file(vocab_config['rna_path'])
    atac_vocab = GeneVocab.from_file(vocab_config['atac_path'])
    cell_vocab = GeneVocab.from_file(vocab_config['cell_type_path'])
    batch_vocab = GeneVocab.from_file(vocab_config['batch_path'])
    chr_vocab = GeneVocab.from_file(vocab_config['chr_path'])
    
    if is_master:
        logger.info(f'Rna vocab size: {len(rna_vocab)}')
        logger.info(f'Atac vocab size: {len(atac_vocab)}')
        
    if is_master:
        logger.info('loading training data')
    
    train_set = PairedSCDataset(
        rna_file = data_config['train']['rna_path'],
        atac_file= data_config['train']['atac_path'],
        rna_key = data_config['train']['rna_key'],
        atac_key = data_config['train']['atac_key'],
        rna_vocab = rna_vocab,
        atac_vocab = atac_vocab,
        cell_vocab = cell_vocab,
        batch_vocab= batch_vocab,
        chr_vocab = chr_vocab,
        gene2chr_file= vocab_config['gene2chr_path'],
        rna_max_len = train_config['model']['rna_max_len'],
        atac_max_len = train_config['model']['atac_max_len'],
        pad_token = pad['token'],
        rna_pad_value = pad['value'],
        cls_token = cls['token'],
        logger = logger,
    )
                                
    gc.collect()
    train_sampler = DistributedSampler(train_set, num_replicas=world_size, rank=rank, shuffle=True)
    train_loader = DataLoader(train_set, batch_size=BATCH_SIZE, sampler=train_sampler, prefetch_factor=4, num_workers=4)

    
    if is_master:
        logger.info('loading validation data')
    val_set = PairedSCDataset(
        rna_file = data_config['test']['rna_path'],
        atac_file= data_config['test']['atac_path'],
        rna_key = data_config['test']['rna_key'],
        atac_key = data_config['test']['atac_key'],
        rna_vocab = rna_vocab,
        atac_vocab = atac_vocab,
        cell_vocab = cell_vocab,
        batch_vocab= batch_vocab,
        chr_vocab = chr_vocab,
        gene2chr_file= vocab_config['gene2chr_path'],
        rna_max_len = train_config['model']['rna_max_len'],
        atac_max_len = train_config['model']['atac_max_len'],
        pad_token = pad['token'],
        rna_pad_value = pad['value'],
        cls_token = cls['token'],
        logger = logger,
    )
    gc.collect()

    val_sampler = SequentialDistributedSampler(val_set, batch_size=BATCH_SIZE, world_size=world_size, rank=rank, num_replicas=world_size)
    val_loader = DataLoader(val_set, batch_size=BATCH_SIZE, sampler=val_sampler, prefetch_factor=4, num_workers=4)
    
    if is_master:
        logger.info('Creating model')
    # init model on CPU
    model = EpiFoundation(
        num_class_cell = len(cell_vocab),
        num_rnas = len(rna_vocab),
        num_atacs = len(atac_vocab),
        num_values= data_config['bin_num'],
        num_chrs= len(chr_vocab),
        embed_dim = train_config['model']['embedding_dim'],
        depth = train_config['model']['num_layers'],
        heads = train_config['model']['head_num'],
        head_dim = train_config['model']['head_dim'],
        encoder = model_name,
        dropout = train_config['model']['dropout'],
        pad_token_idx_rna = rna_vocab[pad['token']],
        pad_token_idx_atac = atac_vocab[pad['token']],
        cell_emb_style = train_config['model']['cell_emb_style'],
        mvc_arch_style = train_config['model']['mvc_arch_style'],
        use_batch_labels = train_config['model']['use_batch_labels'],
        batch_label_num= len(batch_vocab),
        use_chr_labels= train_config['model']['use_chr_labels'],
        transformer_backend='flash',
    )
    
    mixed_precision_policy = MixedPrecision(
        param_dtype=torch.bfloat16,
        reduce_dtype=torch.bfloat16,
        buffer_dtype=torch.bfloat16,
    )
    
    sharding_strategy = ShardingStrategy.FULL_SHARD
    
    model = FSDP(
        module=model,
        mixed_precision=mixed_precision_policy,
        sharding_strategy=sharding_strategy,
        device_id=torch.cuda.current_device(),
    )
    
    # optimizer
    optimizer = Adam(model.parameters(), lr=LEARNING_RATE)
    
    # learning rate scheduler
    scheduler = CosineAnnealingWarmupRestarts(
        optimizer,
        first_cycle_steps=15,
        cycle_mult=2,
        max_lr=LEARNING_RATE,
        min_lr=1e-6,
        warmup_steps=5,
        gamma=0.9
    )
    
    start_epoch = 1
    
    # model = DDP(model, device_ids=[local_rank], output_device=local_rank)
    # scaler = torch.amp.GradScaler(enabled=train_config['amp'].amp)
    scaler = torch.cuda.amp.GradScaler(enabled=train_config['amp'])
    
    # masked_mse_loss = MaskedMSELoss().to(local_rank)
    cross_entropy_loss = nn.CrossEntropyLoss(reduction='mean').to(device)
    
    if data_config['bin_num'] > 1:
        mvc_loss_fn = nn.CrossEntropyLoss(reduction='mean', ignore_index = pad['value']).to(device)
    else:
        mvc_loss_fn = nn.MSELoss(reduction='sum').to(device)
    
    mvc_weight = train_config['task_weight']['mvc']
    cell_type_weight = train_config['task_weight']['cell_type']
    
    softmax = nn.Softmax(dim=-1)
    
    steps = 0
    if train_config['model']['pretrained'] is not None:
        if is_master:
            logger.info('Loading pretrained model from: {}'.format(train_config['model']['pretrained']))
        checkpoint = torch.load(train_config['model']['pretrained'], map_location='cpu')
        
        with FSDP.state_dict_type(model, StateDictType.FULL_STATE_DICT):
            model.load_state_dict(checkpoint['model'])
        
        optimizer.load_state_dict(checkpoint['optimizer'])
        scheduler.load_state_dict(checkpoint['scheduler'])
        scaler.load_state_dict(checkpoint['scaler'])
        if resume:
            start_epoch = checkpoint['epoch'] + 1
            steps = checkpoint['steps']
        del checkpoint
        gc.collect()
    
    dist.barrier()
    if is_master:
        logger.info('Start finetuning from epoch: {}, steps: {}'.format(start_epoch, steps))
    for i in range(start_epoch, start_epoch + EPOCHS):
        train_loader.sampler.set_epoch(i)
        
        if is_master:
            logger.info('Training with {} samples, steps: {}'.format(len(train_loader.dataset), len(train_loader)))
        model.train()
        dist.barrier()
        running_loss = {'mvc': 0.0, 'cell': 0.0, 'total': 0.0}
        cum_acc_cell = 0.0
        cum_acc_value = 0.0
        for index, batch in enumerate(train_loader):
            index += 1
            steps += 1
            rna_values = batch['rna_values'].to(device)
            rna_ids = batch['rna_ids'].to(device)
            atac_ids = batch['atac_ids'].to(device)
            cell_ids = batch['cell_ids'].to(device)
            batch_ids = batch['batch_ids'].to(device)
            rna_chrs = batch['rna_chrs'].to(device)
            atac_chrs = batch['atac_chrs'].to(device)
            
            padding_positions = atac_ids.eq(atac_vocab[pad['token']])
            if index % GRADIENT_ACCUMULATION != 0 and index != len(train_loader):
                with model.no_sync():
                    # with torch.cuda.amp.autocast(enabled=train_config['amp'], dtype= torch.bfloat16):
                    #     # finetue using all expression values, do not mask
                    #     output = model(atac = atac_ids, rna = rna_ids, src_key_padding_mask = padding_positions, batch_id = batch_ids, rna_chrs = rna_chrs, atac_chrs = atac_chrs)

                    #     mvc_loss = mvc_loss_fn(output['mvc_pred'].transpose(1, 2), rna_values) * mvc_weight
                    #     cell_loss = cross_entropy_loss(output['cell_pred'], cell_ids) * cell_type_weight
                    #     loss =  mvc_loss + cell_loss
                        
                    #     running_loss['mvc'] += mvc_loss.item()
                    #     running_loss['cell'] += cell_loss.item()
                    #     running_loss['total'] += loss.item()
                        
                    #     loss = loss / GRADIENT_ACCUMULATION

                    output = model(atac = atac_ids, rna = rna_ids, src_key_padding_mask = padding_positions, batch_id = batch_ids, rna_chrs = rna_chrs, atac_chrs = atac_chrs)

                    mvc_loss = mvc_loss_fn(output['mvc_pred'].transpose(1, 2), rna_values) * mvc_weight
                    cell_loss = cross_entropy_loss(output['cell_pred'], cell_ids) * cell_type_weight
                    loss =  mvc_loss + cell_loss
                    
                    running_loss['mvc'] += mvc_loss.item()
                    running_loss['cell'] += cell_loss.item()
                    running_loss['total'] += loss.item()
                    
                    loss = loss / GRADIENT_ACCUMULATION
                    
                    ######
                    
                    scaler.scale(loss).backward()
            else:
                # with torch.cuda.amp.autocast(enabled=train_config['amp'], dtype= torch.bfloat16):
                #     output = model(atac = atac_ids, rna = rna_ids, src_key_padding_mask = padding_positions, batch_id = batch_ids, rna_chrs = rna_chrs, atac_chrs = atac_chrs)
                    
                #     mvc_loss = mvc_loss_fn(output['mvc_pred'].transpose(1, 2), rna_values) * mvc_weight
                #     cell_loss = cross_entropy_loss(output['cell_pred'], cell_ids) * cell_type_weight
                #     loss =  mvc_loss + cell_loss
                    
                #     running_loss['mvc'] += mvc_loss.item()
                #     running_loss['cell'] += cell_loss.item()
                #     running_loss['total'] += loss.item()
                #     if is_master:
                #         tb_logger.add_scalar('train/mvc_loss', mvc_loss.item(), steps)
                #         tb_logger.add_scalar('train/cell_loss', cell_loss.item(), steps)
                #         tb_logger.add_scalar('train/total_loss', loss.item(), steps)
                #         logger.info(f'Epoch: {i} | Step: {index} | MVC Loss: {mvc_loss:.4f} | Cell Type Loss: {cell_loss:.4f} | Total Loss: {loss:.4f}')
                #     loss = loss / GRADIENT_ACCUMULATION
                    
                output = model(atac = atac_ids, rna = rna_ids, src_key_padding_mask = padding_positions, batch_id = batch_ids, rna_chrs = rna_chrs, atac_chrs = atac_chrs)
                
                mvc_loss = mvc_loss_fn(output['mvc_pred'].transpose(1, 2), rna_values) * mvc_weight
                cell_loss = cross_entropy_loss(output['cell_pred'], cell_ids) * cell_type_weight
                loss =  mvc_loss + cell_loss
                
                running_loss['mvc'] += mvc_loss.item()
                running_loss['cell'] += cell_loss.item()
                running_loss['total'] += loss.item()
                if is_master:
                    tb_logger.add_scalar('train/mvc_loss', mvc_loss.item(), steps)
                    tb_logger.add_scalar('train/cell_loss', cell_loss.item(), steps)
                    tb_logger.add_scalar('train/total_loss', loss.item(), steps)
                    logger.info(f'Epoch: {i} | Step: {index} | MVC Loss: {mvc_loss:.4f} | Cell Type Loss: {cell_loss:.4f} | Total Loss: {loss:.4f}')
                loss = loss / GRADIENT_ACCUMULATION
                #######
                    
                scaler.scale(loss).backward()
                scaler.unscale_(optimizer)
                torch.nn.utils.clip_grad_norm_(model.parameters(), int(1e2))
                scaler.step(optimizer)
                scaler.update()
                optimizer.zero_grad()
            # cell type accuracy
            type_pred = softmax(output['cell_pred'])
            type_pred = type_pred.argmax(dim=-1)
            cum_acc_cell += (type_pred.eq(cell_ids)).sum().item() / len(cell_ids)
            
            value_pred = softmax(output['mvc_pred']).argmax(dim=-1)
            # expression value accuracy
            non_pad_idx = rna_values.ne(pad['value'])
            non_pad_pred = value_pred[non_pad_idx]
            non_pad_label = rna_values[non_pad_idx]
            cum_acc_value += (non_pad_pred.eq(non_pad_label).sum().item()) / non_pad_label.size(0)
            
        cum_acc_cell = 100 * cum_acc_cell / index
        cum_acc_cell = get_reduced(cum_acc_cell, rank, 0, world_size)
        
        cum_acc_value = 100 * cum_acc_value / index
        cum_acc_value = get_reduced(cum_acc_value, rank, 0, world_size)
        for key in running_loss:
            running_loss[key] = running_loss[key] / index
            running_loss[key] = get_reduced(running_loss[key], rank, 0, world_size)
        if is_master:
                logger.info(f'Epoch: {i} | MVC Loss: {running_loss["mvc"]:.4f} | Cell Type Loss: {running_loss["cell"]:.4f} | Total Loss: {running_loss["total"]:.4f} | Cell Type Accuracy: {cum_acc_cell:.2f} | Expression Value Accuracy: {cum_acc_value:.2f}')
        dist.barrier()
        scheduler.step()
        # del train_set, train_sampler, train_loader

        if i % valid_config['freq'] == 0:
            if is_master:
                logger.info('#### Validation ####')
            model.eval()
            dist.barrier()
            running_loss = {'mvc': 0.0, 'cell': 0.0, 'total': 0.0}
            
            cum_acc_cell = 0.0
            cum_acc_value = 0.0

            with torch.no_grad():
                for index, batch in enumerate(val_loader):
                    index += 1
                    
                    rna_values = batch['rna_values'].to(device)
                    rna_ids = batch['rna_ids'].to(device)
                    atac_ids = batch['atac_ids'].to(device)
                    cell_ids = batch['cell_ids'].to(device)
                    batch_ids = batch['batch_ids'].to(device)
                    rna_chrs = batch['rna_chrs'].to(device)
                    atac_chrs = batch['atac_chrs'].to(device)

                    padding_positions = atac_ids.eq(atac_vocab[pad['token']])
                    # with torch.cuda.amp.autocast(enabled=train_config['amp'], dtype= torch.bfloat16):
                    #     output = model(atac = atac_ids, rna = rna_ids, src_key_padding_mask = padding_positions, batch_id = batch_ids, rna_chrs = rna_chrs, atac_chrs = atac_chrs)
                        
                    #     mvc_loss = mvc_loss_fn(output['mvc_pred'].transpose(1, 2), rna_values) * mvc_weight
                    #     cell_loss = cross_entropy_loss(output['cell_pred'], cell_ids) * cell_type_weight
                    #     loss =  mvc_loss + cell_loss
                        
                    output = model(atac = atac_ids, rna = rna_ids, src_key_padding_mask = padding_positions, batch_id = batch_ids, rna_chrs = rna_chrs, atac_chrs = atac_chrs)
                    
                    mvc_loss = mvc_loss_fn(output['mvc_pred'].transpose(1, 2), rna_values) * mvc_weight
                    cell_loss = cross_entropy_loss(output['cell_pred'], cell_ids) * cell_type_weight
                    loss =  mvc_loss + cell_loss
                        
                    running_loss['mvc'] += mvc_loss.item()
                    running_loss['cell'] += cell_loss.item()
                    running_loss['total'] += loss.item()
                    
                    type_pred = softmax(output['cell_pred'])
                    type_pred = type_pred.argmax(dim=-1)
                    cum_acc_cell += (type_pred.eq(cell_ids)).sum().item() / len(cell_ids)
                    
                    value_pred = softmax(output['mvc_pred']).argmax(dim=-1)
                    # expression value accuracy
                    non_pad_idx = rna_values.ne(pad['value'])
                    non_pad_pred = value_pred[non_pad_idx]
                    non_pad_label = rna_values[non_pad_idx]
                    cum_acc_value += (non_pad_pred.eq(non_pad_label).sum().item()) / non_pad_label.size(0)
            for key in running_loss:
                running_loss[key] = running_loss[key] / index
                running_loss[key] = get_reduced(running_loss[key], rank, 0, world_size)
            cum_acc_cell = 100 * cum_acc_cell / index
            cum_acc_cell = get_reduced(cum_acc_cell, rank, 0, world_size)
            
            cum_acc_value = 100 * cum_acc_value / index
            cum_acc_value = get_reduced(cum_acc_value, rank, 0, world_size)
            
            # del val_set, val_sampler, val_loader
            if is_master:
                logger.info(f'MVC Loss: {running_loss["mvc"]:.4f} | Cell Type Loss: {running_loss["cell"]:.4f} | Total Loss: {running_loss["total"]:.4f} | Cell Type Accuracy: {cum_acc_cell:.2f} | Expression Value Accuracy: {cum_acc_value:.2f}')
                
        if i % save_ckpt_freq == 0 or i == EPOCHS:
            save_ckpt_fsdp(i, steps,  model, optimizer, scheduler, scaler, running_loss["total"], task_name, ckpt_dir, rank=rank)
            if is_master:
                logger.info('Model saved at epoch: {}'.format(i))
    

if __name__ == '__main__':
    main()
    
```
