## Supplementary material for "EpiFoundation: A Foundation Model for Single-Cell ATAC-seq via Peak-to-Gene Alignment": Code for EpiFoundation: README.md

# EpiFoundation: A Foundation Model for Single-Cell ATAC-seq via Peak-to-Gene Alignment

This repo contains official Pytorch implementation of **EpiFoundation** in our paper: [EpiFoundation: A Foundation Model for Single-Cell ATAC-seq via Peak-to-Gene Alignment]()

![image-20250204132627694](./assets/framework.png)

## Introduction

Foundation models exhibit strong capabilities for downstream tasks by learning generalized representations through self-supervised pre-training on large datasets. While several foundation models have been developed for single-cell RNA-seq (scRNA-seq) data, there is still a lack of models specifically tailored for single-cell ATAC-seq (scATAC-seq), which measures epigenetic information in individual cells. The principal challenge in developing such a model lies in the vast number of scATAC peaks and the significant sparsity of the data, which complicates the formulation of peak-to-peak correlations. To address this challenge, we introduce **EpiFoundation**, a foundation model for learning cell representations from the high-dimensional and sparse space of peaks. EpiFoundation relies on an innovative cross-modality pre-training procedure with two key technical innovations. First, EpiFoundation exclusively processes the non-zero peak set, thereby enhancing the density of cell-specific information within the input data. Second, EpiFoundation utilizes dense gene expression information to supervise the pre-training process, aligning peak-to-gene correlations. EpiFoundation can handle various types of downstream tasks, including cell-type annotation, batch correction, and gene expression prediction. To train and validate EpiFoundation, we curated **MiniAtlas**, a dataset of 100,000+ single cells with paired scRNA-seq and scATAC-seq data, along with diverse test sets spanning various tissues and cell types for robust evaluation. EpiFoundation demonstrates **state-of-the-art performance across multiple tissues and diverse downstream tasks**. 

![image-20250204133126719](./assets/batch_correlation.png)


## Data and model Access

- Our paired scRNA-seq and scATAC-seq data for pre-training and fine-tuning can be found at [MiniAtlas](https://huggingface.co/datasets/UCSC-VLAA/MiniAtlas).
- The pre-trained model weights can be found at [EpiFoundation](https://huggingface.co/UCSC-VLAA/EpiFoundation).

## Quick Start

0. prepare conda env from `env.yml`

1. Preprocess your data using `prepare_data.py`

   - call `preprocess()` to process scRNA-seq data, divide the data into train, valid and test set, and perform binning.
   - call `get_pair_data()` to get the paired scATAC-seq data
   - generation of vocabularies
   - Optional: call reduce_data() to transform data into sparse format and reduce the data size.


2. Prepare batch data:

   get the cell and gene vocab:

   ```python
   vocab = GeneVocab.from_file(vocab_config['path'])
   cell_vocab = GeneVocab.from_file(vocab_config['cell_type_path'])
   ```

   Init the paired dataset:

   ```python
   train_set = PairedSCDataset(
        rna_file = data_config['train']['rna_path'],
        atac_file= data_config['train']['atac_path'],
        rna_key = data_config['train']['rna_key'],
        atac_key = data_config['train']['atac_key'],
        rna_vocab = rna_vocab,
        atac_vocab = atac_vocab,
        cell_vocab = cell_vocab,
        batch_vocab= batch_vocab,
        chr_vocab = chr_vocab,
        gene2chr_file= vocab_config['gene2chr_path'],
        rna_max_len = train_config['model']['rna_max_len'],
        atac_max_len = train_config['model']['atac_max_len'],
        pad_token = pad['token'],
        rna_pad_value = pad['value'],
        cls_token = cls['token'],
        # reg_token= reg['token'],
        logger = logger,
    )
   ```

   Get the data loader

   ```python
   train_sampler = DistributedSampler(train_set)
   train_loader = DataLoader(train_set, batch_size=BATCH_SIZE, sampler=train_sampler, num_workers=4)
   ```

   Detailed code can be find in `pretrain_ddp.py`, `pretrain_fsdp.py` and `data/dataloader.py`

3. Construct the model using config from ./configs/test/rna_transformer.yml 

   ```python
   model = EpiFoundation(
        num_class_cell = len(cell_vocab),
        num_rnas = len(rna_vocab),
        num_atacs = len(atac_vocab),
        num_values= data_config['bin_num'],
        num_chrs= len(chr_vocab),
        embed_dim = train_config['model']['embedding_dim'],
        depth = train_config['model']['num_layers'],
        heads = train_config['model']['head_num'],
        head_dim = train_config['model']['head_dim'],
        encoder = model_name,
        dropout = train_config['model']['dropout'],
        pad_token_idx_rna = rna_vocab[pad['token']],
        pad_token_idx_atac = atac_vocab[pad['token']],
        cell_emb_style = train_config['model']['cell_emb_style'],
        mvc_arch_style = train_config['model']['mvc_arch_style'],
        use_batch_labels = train_config['model']['use_batch_labels'],
        batch_label_num= len(batch_vocab),
        use_chr_labels= train_config['model']['use_chr_labels'],
    ).to(device)
   ```


## Acknowledgment

We would like to thank the TPU Research Cloud (TRC) program and the Google Cloud Research Credits program for Research program for supporting our computing needs. W.H. and Z.J. are supported by the National Institute Of General Medical Sciences of the National Institutes of Health (NIH), under Award Number R35GM150887 and R35GM154865 respectively.

## Citation

```
```

## Contact

If you have any questions, please feel free to raise an issue or contact us directly: Juncheng Wu
