## Supplementary material for "EpiFoundation: A Foundation Model for Single-Cell ATAC-seq via Peak-to-Gene Alignment": Code for EpiFoundation: gene_tokenizer.py

```
import json
import pickle
from pathlib import Path
from collections import Counter, OrderedDict
from typing import Dict, Iterable, List, Optional, Tuple, Union
from typing_extensions import Self

import numpy as np
import pandas as pd
import torch
import torchtext.vocab as torch_vocab
from torchtext.vocab import Vocab

class GeneVocab(Vocab):
    """
    Vocabulary for genes.
    """

    def __init__(
        self,
        gene_list_or_vocab: Union[List[str], Vocab],
        specials: Optional[List[str]] = None,
        special_first: bool = True,
        default_token: Optional[str] = "<pad>",
    ) -> None:
        """
        Initialize the vocabulary.
        Note: add specials only works when init from a gene list.

        Args:
            gene_list_or_vocab (List[str] or Vocab): List of gene names or a
                Vocab object.
            specials (List[str]): List of special tokens.
            special_first (bool): Whether to add special tokens to the beginning
                of the vocabulary.
            default_token (str): Default token, by default will set to "<pad>",
                if "<pad>" is in the vocabulary.
        """
        if isinstance(gene_list_or_vocab, Vocab):
            _vocab = gene_list_or_vocab
            if specials is not None:
                raise ValueError(
                    "receive non-empty specials when init from a Vocab object."
                )
        elif isinstance(gene_list_or_vocab, list):
            _vocab = self._build_vocab_from_iterator(
                gene_list_or_vocab,
                specials=specials,
                special_first=special_first,
            )
        else:
            raise ValueError(
                "gene_list_or_vocab must be a list of gene names or a Vocab object."
            )
        super().__init__(_vocab.vocab)
        if default_token is not None and default_token in self:
            self.set_default_token(default_token)

    @classmethod
    def from_file(cls, file_path: Union[Path, str]) -> Self:
        """
        Load the vocabulary from a file. The file should be either a pickle or a
        json file of token to index mapping.
        """
        if isinstance(file_path, str):
            file_path = Path(file_path)
        if file_path.suffix == ".pkl":
            with file_path.open("rb") as f:
                vocab = pickle.load(f)
                return cls(vocab)
        elif file_path.suffix == ".json":
            with file_path.open("r") as f:
                token2idx = json.load(f)
                return cls.from_dict(token2idx)
        else:
            raise ValueError(
                f"{file_path} is not a valid file type. "
                "Only .pkl and .json are supported."
            )

    @classmethod
    def from_dict(
        cls,
        token2idx: Dict[str, int],
        default_token: Optional[str] = "<pad>",
    ) -> Self:
        """
        Load the vocabulary from a dictionary.

        Args:
            token2idx (Dict[str, int]): Dictionary mapping tokens to indices.
        """
        # initiate an empty vocabulary first
        _vocab = cls([])

        # add the tokens to the vocabulary, GeneVocab requires consecutive indices
        for t, i in sorted(token2idx.items(), key=lambda x: x[1]):
            _vocab.insert_token(t, i)

        if default_token is not None and default_token in _vocab:
            _vocab.set_default_token(default_token)

        return _vocab

    def _build_vocab_from_iterator(
        self,
        iterator: Iterable,
        min_freq: int = 1,
        specials: Optional[List[str]] = None,
        special_first: bool = True,
    ) -> Vocab:
        """
        Build a Vocab from an iterator. This function is modified from
        torchtext.vocab.build_vocab_from_iterator. The original function always
        splits tokens into characters, which is not what we want.

        Args:
            iterator (Iterable): Iterator used to build Vocab. Must yield list
                or iterator of tokens.
            min_freq (int): The minimum frequency needed to include a token in
                the vocabulary.
            specials (List[str]): Special symbols to add. The order of supplied
                tokens will be preserved.
            special_first (bool): Whether to add special tokens to the beginning

        Returns:
            torchtext.vocab.Vocab: A `Vocab` object
        """

        counter = Counter()
        counter.update(iterator)

        if specials is not None:
            for tok in specials:
                del counter[tok]

        sorted_by_freq_tuples = sorted(counter.items(), key=lambda x: x[0])
        sorted_by_freq_tuples.sort(key=lambda x: x[1], reverse=True)
        ordered_dict = OrderedDict(sorted_by_freq_tuples)

        if specials is not None:
            if special_first:
                specials = specials[::-1]
            for symbol in specials:
                ordered_dict.update({symbol: min_freq})
                ordered_dict.move_to_end(symbol, last=not special_first)

        word_vocab = torch_vocab.vocab(ordered_dict, min_freq=min_freq)
        return word_vocab

    @property
    def pad_token(self) -> Optional[str]:
        """
        Get the pad token.
        """
        if getattr(self, "_pad_token", None) is None:
            self._pad_token = None
        return self._pad_token

    @pad_token.setter
    def pad_token(self, pad_token: str) -> None:
        """
        Set the pad token. Will not add the pad token to the vocabulary.

        Args:
            pad_token (str): Pad token, should be in the vocabulary.
        """
        if pad_token not in self:
            raise ValueError(f"{pad_token} is not in the vocabulary.")
        self._pad_token = pad_token

    def save_json(self, file_path: Union[Path, str]) -> None:
        """
        Save the vocabulary to a json file.
        """
        if isinstance(file_path, str):
            file_path = Path(file_path)
        with file_path.open("w") as f:
            json.dump(self.get_stoi(), f, indent=2)

    def set_default_token(self, default_token: str) -> None:
        """
        Set the default token.

        Args:
            default_token (str): Default token.
        """
        if default_token not in self:
            raise ValueError(f"{default_token} is not in the vocabulary.")
        self.set_default_index(self[default_token])
```
