## Supplementary material for "EpiFoundation: A Foundation Model for Single-Cell ATAC-seq via Peak-to-Gene Alignment": Code for EpiFoundation: utils.py

```
# -*- coding: utf-8 -*-

from __future__ import print_function
import json
import os
import struct
import sys
import platform
import re
import time
import traceback
import requests
import socket
import random
import math
import numpy as np
import torch
import logging
import datetime
from torch.optim.lr_scheduler import _LRScheduler
from torch import nn
import torch.nn.functional as F
from torch.nn.modules.loss import _WeightedLoss
from typing import Dict, Optional

import torch.distributed as dist
from torch.nn.parallel import DistributedDataParallel as DDP
# DDP imports
from torch.distributed.fsdp import FullyShardedDataParallel as FSDP
# FSDP imports
from torch.distributed.fsdp.fully_sharded_data_parallel import (
    CPUOffload, MixedPrecision, BackwardPrefetch, ShardingStrategy
)
from torch.distributed.fsdp.wrap import (
    size_based_auto_wrap_policy,
    transformer_auto_wrap_policy,
    enable_wrap,
    wrap
)
from torch.distributed.fsdp import (
    FullStateDictConfig,
    StateDictType
)


def seed_all(seed_value, cuda_deterministic=False):
    """
    set all random seeds
    """
    random.seed(seed_value)
    os.environ['PYTHONHASHSEED'] = str(seed_value)
    # np.random.seed(seed_value)
    torch.manual_seed(seed_value)
    if torch.cuda.is_available():
        torch.cuda.manual_seed(seed_value)
        torch.cuda.manual_seed_all(seed_value)
    # Speed-reproducibility tradeoff https://pytorch.org/docs/stable/notes/randomness.html
    if cuda_deterministic:  # slower, more reproducible
        torch.backends.cudnn.deterministic = True
        torch.backends.cudnn.benchmark = False
    else:  # faster, less reproducible
        torch.backends.cudnn.deterministic = False
        torch.backends.cudnn.benchmark = True


def set_log(log_dir, rank = -1):
    """
    save log
    """
    time_now = datetime.datetime.now()
    log_file = os.path.join(log_dir, f'{time_now.year}_{time_now.month}_{time_now.day}_{time_now.hour}_{time_now.minute}.log')
    if not os.path.exists(log_dir):
        os.makedirs(log_dir)
    else:
        pass

    logging.basicConfig(level=logging.INFO if rank in [-1, 0] else logging.WARN,
        format='[%(asctime)s %(levelname)s %(filename)s line %(lineno)d %(process)d] %(message)s',
        datefmt='[%X]',
        handlers=[logging.FileHandler(log_file), logging.StreamHandler()]
    )
    logger = logging.getLogger()
    return logger

def dict2str(opt, indent_level=1):
    """dict to string for printing options.

    Args:
        opt (dict): Option dict.
        indent_level (int): Indent level. Default: 1.

    Return:
        (str): Option string for printing.
    """
    msg = '\n'
    for k, v in opt.items():
        if isinstance(v, dict):
            msg += ' ' * (indent_level * 2) + k + ':['
            msg += dict2str(v, indent_level + 1)
            msg += ' ' * (indent_level * 2) + ']\n'
        else:
            msg += ' ' * (indent_level * 2) + k + ': ' + str(v) + '\n'
    return msg


def save_ckpt_fsdp(epoch, steps, model, optimizer, scheduler, scaler, loss, task_name, ckpt_dir, rank):
    # Ensure all processes are synchronized
    dist.barrier()
    # Switch to FULL_STATE_DICT context to gather full state_dict
    with FSDP.state_dict_type(model, StateDictType.FULL_STATE_DICT):
        full_state_dict = model.state_dict()
    # Only save on the master process
    if rank == 0:
        if not os.path.exists(ckpt_dir):
            os.makedirs(ckpt_dir)
        ckpt = {
            'epoch': epoch,
            'steps': steps,
            'model': full_state_dict,
            'optimizer': optimizer.state_dict(),
            'scheduler': scheduler.state_dict(),
            'scaler': scaler.state_dict(),
            'loss': loss,
        }
        ckpt_path = os.path.join(ckpt_dir, f'checkpoint_{epoch}.pth')
        torch.save(ckpt, ckpt_path)

def save_ckpt(epoch, steps,  model, optimizer, scheduler, scaler, losses, model_name, ckpt_folder):
    """
    save checkpoint
    """
    if not os.path.exists(ckpt_folder):
        os.makedirs(ckpt_folder)
    torch.save(
        {
            'epoch': epoch,
            'steps': steps,
            'model': model.module.state_dict(),
            'optimizer': optimizer.state_dict(),
            'scheduler': scheduler.state_dict(),
            'scaler': scaler.state_dict(),
            'losses': losses,
        },
        f'{ckpt_folder}/Epoch_{epoch}_Step_{steps}_{model_name}.pth',
    )
    


def get_reduced(tensor, current_device, dest_device, world_size):
    """
    garther tensor from different GPUs to the main GPU
    """
    tensor = torch.tensor(tensor, device ='cuda')
    torch.distributed.reduce(tensor, dst=dest_device, op = dist.ReduceOp.SUM)
    tensor_mean = tensor.item() / world_size
    return tensor_mean

def get_ndtensor_reduced(tensor, current_device, dest_device, world_size):
    """
    garther tensor from different GPUs to the main GPU
    """
    tensor = tensor.clone().detach() if torch.is_tensor(tensor) else torch.tensor(tensor)
    tensor = tensor.to(current_device)
    torch.distributed.reduce(tensor, dst=dest_device)
    tensor_mean = torch.zeros(tensor.shape)
    if len(tensor.shape) == 2:
        for i in range(tensor.shape[0]):
            for j in range(tensor.shape[1]):
                tensor_mean[i,j] = tensor[i,j].item() / world_size
    elif len(tensor.shape) == 1:
        for i in range(tensor.shape[0]):
            tensor_mean[i] = tensor[i].item() / world_size
    return tensor_mean

def numel(m: torch.nn.Module, only_trainable: bool = False):
    """
    returns the total number of parameters used by `m` (only counting
    shared parameters once); if `only_trainable` is True, then only
    includes parameters with `requires_grad = True`
    """
    parameters = m.parameters()
    if only_trainable:
        parameters = list(p for p in parameters if p.requires_grad)
    unique = dict((p.data_ptr(), p) for p in parameters).values()
    return sum(p.numel() for p in unique)


def label_smooth(y, K, epsilon=0.1):
    """
    Label smoothing for multiclass labels
    One hot encode labels `y` over `K` classes. `y` should be of the form [1, 6, 3, etc.]
    """
    m = len(y)
    out = np.ones((m, K)) * epsilon / K
    for index in range(m):
        out[index][y[index] - 1] += 1 - epsilon
    return torch.tensor(out)


class SequentialDistributedSampler(torch.utils.data.sampler.Sampler):
    """
    Distributed Sampler that subsamples indicies sequentially,
    making it easier to collate all results at the end.
    Even though we only use this sampler for eval and predict (no training),
    which means that the model params won't have to be synced (i.e. will not hang
    for synchronization even if varied number of forward passes), we still add extra
    samples to the sampler to make it evenly divisible (like in `DistributedSampler`)
    to make it easy to `gather` or `reduce` resulting tensors at the end of the loop.
    """

    def __init__(self, dataset, batch_size, world_size, rank=None, num_replicas=None):
        if num_replicas is None:
            if not torch.distributed.is_available():
                raise RuntimeError("Requires distributed package to be available")
            num_replicas = world_size
        if rank is None:
            if not torch.distributed.is_available():
                raise RuntimeError("Requires distributed package to be available")
            rank = torch.distributed.get_rank()
        self.dataset = dataset
        self.num_replicas = num_replicas
        self.rank = rank
        self.batch_size = batch_size
        self.num_samples = int(math.ceil(len(self.dataset) * 1.0 / self.batch_size / self.num_replicas)) * self.batch_size
        self.total_size = self.num_samples * self.num_replicas

    def __iter__(self):
        indices = list(range(len(self.dataset)))
        # add extra samples to make it evenly divisible
        indices += [indices[-1]] * (self.total_size - len(indices))
        # subsample
        indices = indices[self.rank * self.num_samples : (self.rank + 1) * self.num_samples]
        return iter(indices)

    def __len__(self):
        return self.num_samples


def distributed_concat(tensor, num_total_examples, world_size):
    """
    merge the inference results of different processes
    """
    output_tensors = [tensor.clone() for _ in range(world_size)]
    torch.distributed.all_gather(output_tensors, tensor)
    concat = torch.cat(output_tensors, dim=0)
    # truncate the dummy elements added by SequentialDistributedSampler
    return concat[:num_total_examples]


class CosineAnnealingWarmupRestarts(_LRScheduler):
    """
        optimizer (Optimizer): Wrapped optimizer.
        first_cycle_steps (int): First cycle step size.
        cycle_mult(float): Cycle steps magnification. Default: -1.
        max_lr(float): First cycle's max learning rate. Default: 0.1.
        min_lr(float): Min learning rate. Default: 0.001.
        warmup_steps(int): Linear warmup step size. Default: 0.
        gamma(float): Decrease rate of max learning rate by cycle. Default: 1.
        last_epoch (int): The index of last epoch. Default: -1.
    """
    
    def __init__(self,
                 optimizer : torch.optim.Optimizer,
                 first_cycle_steps : int,
                 cycle_mult : float = 1.,
                 max_lr : float = 0.1,
                 min_lr : float = 0.001,
                 warmup_steps : int = 0,
                 gamma : float = 1.,
                 last_epoch : int = -1
        ):
        assert warmup_steps < first_cycle_steps
        
        self.first_cycle_steps = first_cycle_steps # first cycle step size
        self.cycle_mult = cycle_mult # cycle steps magnification
        self.base_max_lr = max_lr # first max learning rate
        self.max_lr = max_lr # max learning rate in the current cycle
        self.min_lr = min_lr # min learning rate
        self.warmup_steps = warmup_steps # warmup step size
        self.gamma = gamma # decrease rate of max learning rate by cycle
        
        self.cur_cycle_steps = first_cycle_steps # first cycle step size
        self.cycle = 0 # cycle count
        self.step_in_cycle = last_epoch # step size of the current cycle
        
        super(CosineAnnealingWarmupRestarts, self).__init__(optimizer, last_epoch)
        
        # set learning rate min_lr
        self.init_lr()

    def init_lr(self):
        self.base_lrs = []
        for param_group in self.optimizer.param_groups:
            param_group['lr'] = self.min_lr
            self.base_lrs.append(self.min_lr)
    
    def get_lr(self):
        if self.step_in_cycle == -1:
            return self.base_lrs
        elif self.step_in_cycle < self.warmup_steps:
            return [(self.max_lr - base_lr)*self.step_in_cycle / self.warmup_steps + base_lr for base_lr in self.base_lrs]
        else:
            return [base_lr + (self.max_lr - base_lr) \
                    * (1 + math.cos(math.pi * (self.step_in_cycle-self.warmup_steps) \
                                    / (self.cur_cycle_steps - self.warmup_steps))) / 2
                    for base_lr in self.base_lrs]

    def step(self, epoch=None):
        if epoch is None:
            epoch = self.last_epoch + 1
            self.step_in_cycle = self.step_in_cycle + 1
            if self.step_in_cycle >= self.cur_cycle_steps:
                self.cycle += 1
                self.step_in_cycle = self.step_in_cycle - self.cur_cycle_steps
                self.cur_cycle_steps = int((self.cur_cycle_steps - self.warmup_steps) * self.cycle_mult) + self.warmup_steps
        else:
            if epoch >= self.first_cycle_steps:
                if self.cycle_mult == 1.:
                    self.step_in_cycle = epoch % self.first_cycle_steps
                    self.cycle = epoch // self.first_cycle_steps
                else:
                    n = int(math.log((epoch / self.first_cycle_steps * (self.cycle_mult - 1) + 1), self.cycle_mult))
                    self.cycle = n
                    self.step_in_cycle = epoch - int(self.first_cycle_steps * (self.cycle_mult ** n - 1) / (self.cycle_mult - 1))
                    self.cur_cycle_steps = self.first_cycle_steps * self.cycle_mult ** (n)
            else:
                self.cur_cycle_steps = self.first_cycle_steps
                self.step_in_cycle = epoch
                
        self.max_lr = self.base_max_lr * (self.gamma**self.cycle)
        self.last_epoch = math.floor(epoch)
        for param_group, lr in zip(self.optimizer.param_groups, self.get_lr()):
            param_group['lr'] = lr


class DistanceLoss(_WeightedLoss):
    """
    CrossEntropyLoss with Distance Weighted
    """
    def __init__(self, weight=None, reduction='mean', ignore_index = None):
        super().__init__(weight=weight, reduction=reduction)
        self.weight = weight
        self.reduction = reduction
        self.ignore_index = ignore_index
    def forward(self, inputs, targets):
        if len(inputs.shape) > 2:
            inputs = inputs.reshape(-1, inputs.size(-1))
        if len(targets.shape) > 1:
            targets = targets.reshape(-1)
        if self.ignore_index is not None:
            keep_index = (targets != self.ignore_index).nonzero(as_tuple=True)[0]
            targets = torch.index_select(targets, 0, keep_index) #targets[targets != self.ignore_index]
            inputs = torch.index_select(inputs, 0, keep_index)
        lsm = F.log_softmax(inputs, -1)
        targets = torch.empty(size=(targets.size(0), inputs.size(-1)), device=targets.device).fill_(0).scatter_(1, targets.data.unsqueeze(1), 1)
        if self.weight is not None:
            lsm = lsm * self.weight.unsqueeze(0)
        loss = -(targets * lsm).sum(-1)
        inputs = nn.Softmax(dim=-1)(inputs)[..., 1:-1].argmax(dim=-1) + 1
        # print('inputs', inputs.device, inputs.shape)
        targets = nn.Softmax(dim=-1)(targets)[..., 1:-1].argmax(dim=-1) + 1
        # print('targets', targets.device, targets.shape)
        distance = abs(inputs - targets) + 1e-2
        # print('loss.shape', loss.shape)
        # print('distance.shape', distance.shape)
        loss = loss * distance
        if self.reduction == 'sum':
            loss = loss.sum()
        elif self.reduction == 'mean':
            loss = loss.mean()
        return loss


class LabelSmoothCrossEntropyLoss(_WeightedLoss):
    """
    CrossEntropyLoss with Label Somoothing
    """
    def __init__(self, weight=None, reduction='mean', smoothing=0.0):
        super().__init__(weight=weight, reduction=reduction)
        self.smoothing = smoothing
        self.weight = weight
        self.reduction = reduction

    @staticmethod
    def _smooth_one_hot(targets: torch.Tensor, n_classes: int, smoothing=0.0):
        assert 0 <= smoothing < 1
        with torch.no_grad():
            targets = torch.empty(size=(targets.size(0), n_classes),
                                  device=targets.device) \
                .fill_(smoothing / (n_classes - 1)) \
                .scatter_(1, targets.data.unsqueeze(1), 1. - smoothing)
        return targets

    def forward(self, inputs, targets):
        targets = LabelSmoothCrossEntropyLoss._smooth_one_hot(targets, inputs.size(-1),
                                                              self.smoothing)
        lsm = F.log_softmax(inputs, -1)

        if self.weight is not None:
            lsm = lsm * self.weight.unsqueeze(0)

        loss = -(targets * lsm).sum(-1)

        if self.reduction == 'sum':
            loss = loss.sum()
        elif self.reduction == 'mean':
            loss = loss.mean()

        return loss
    
    
def eval_scib_metrics(
    adata,
    logger,
    batch_key: str = "str_batch",
    label_key: str = "celltype",
    notes: Optional[str] = None
):
    import scib

    logger.info("Calculating metrics...")
    results = scib.metrics.metrics(
        adata,
        adata_int=adata,
        batch_key=batch_key,
        label_key=label_key,
        embed="embedding",
        isolated_labels_asw_=False,
        silhouette_=True,
        hvg_score_=False,
        graph_conn_=True,
        pcr_=True,
        isolated_labels_f1_=False,
        trajectory_=False,
        nmi_=True,  # use the clustering, bias to the best matching
        ari_=True,  # use the clustering, bias to the best matching
        cell_cycle_=False,
        kBET_=False,  # kBET return nan sometimes, need to examine
        ilisi_=False,
        clisi_=False,
    )

    if notes is not None:
        logger.info(f"{notes}")

    logger.info(f"{results}")

    result_dict = results[0].to_dict()
    logger.info(
        "Biological Conservation Metrics: \n"
        f"ASW (cell-type): {result_dict['ASW_label']:.4f}, graph cLISI: {result_dict['cLISI']:.4f}, "
        f"isolated label silhouette: {result_dict['isolated_label_silhouette']:.4f}, \n"
        "Batch Effect Removal Metrics: \n"
        f"PCR_batch: {result_dict['PCR_batch']:.4f}, ASW (batch): {result_dict['ASW_label/batch']:.4f}, "
        f"graph connectivity: {result_dict['graph_conn']:.4f}, graph iLISI: {result_dict['iLISI']:.4f}"
    )

    result_dict["avg_bio"] = np.mean(
        [
            result_dict["NMI_cluster/label"],
            result_dict["ARI_cluster/label"],
            result_dict["ASW_label"],
        ]
    )

    # remove nan value in result_dict
    result_dict = {k: v for k, v in result_dict.items() if not np.isnan(v)}

    return result_dict
```
