## Supplementary figures and images for "EpiFoundation: A Foundation Model for Single-Cell ATAC-seq via Peak-to-Gene Alignment"

### batch_correlation.png

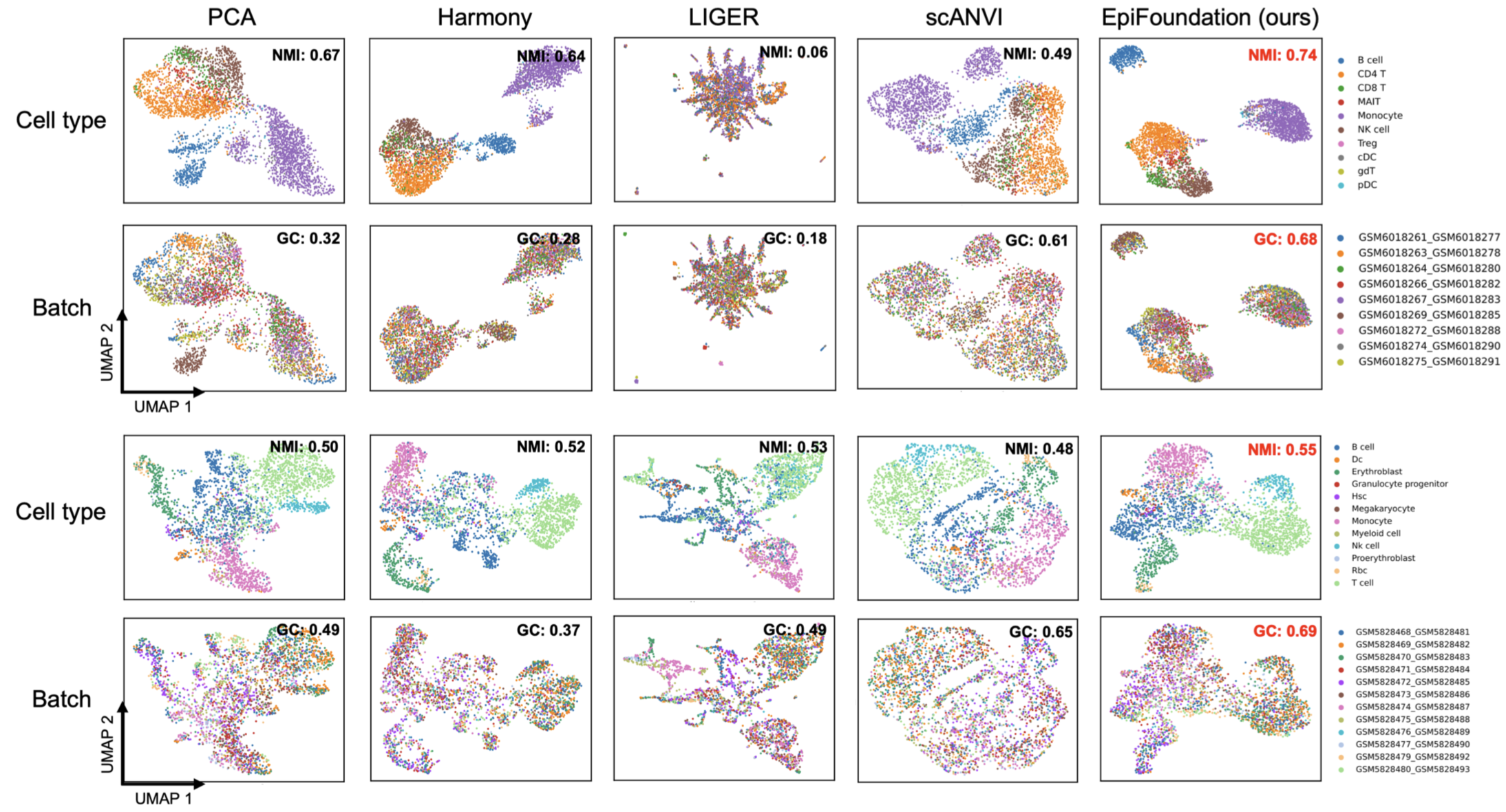

### framework.png

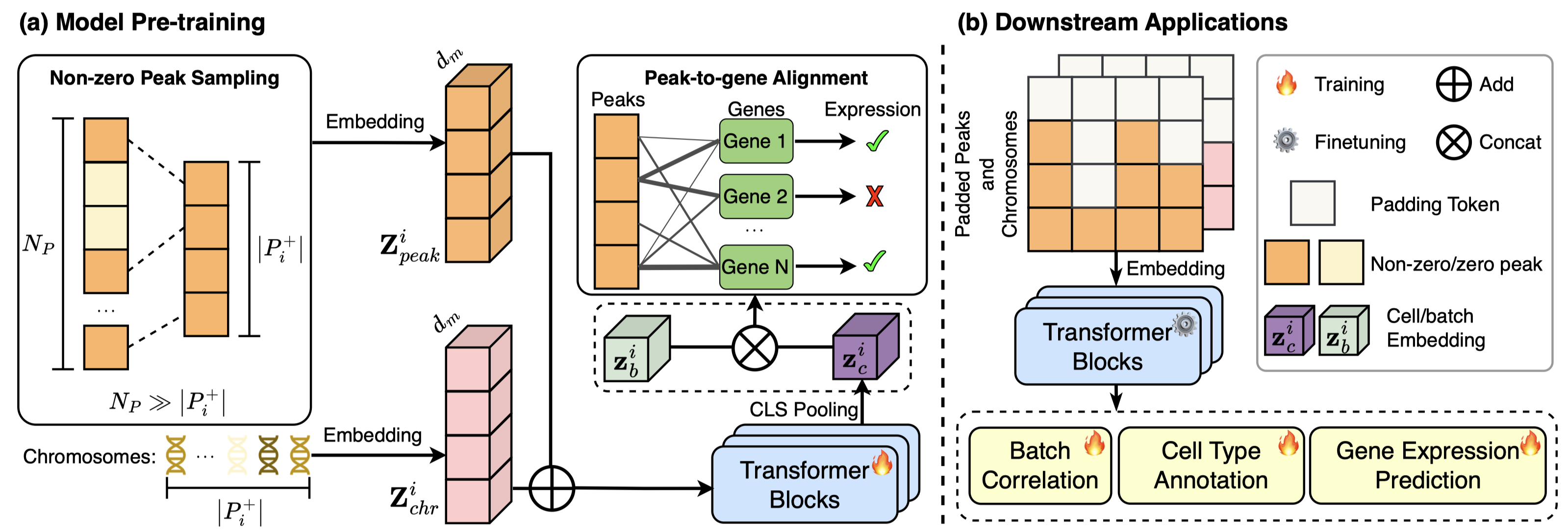
